# Population history shapes urban evolutionary dynamics: distinct genetic structure across urban and rural Europe in two lepidopterans

**DOI:** 10.1101/2025.01.27.635009

**Authors:** Sami M. Kivelä, Kyung Min Lee, George C. Adamidis, Simona Bonelli, Frederik Hendrickx, Peter Huemer, Tomáš Kadlec, Tuomas Kankaanpää, Guillaume Minard, Matthew E. Nielsen, Elena Piano, Toomas Tammaru, Konstantina Zografou, Thomas Merckx

## Abstract

Urbanisation is transforming environments globally. The altered abiotic conditions and biotic interactions in urban habitats impose divergent selection pressures on urban versus rural populations, while genetic drift may also be significant in typically small urban populations. A key question in urban evolution concerns the origin and spread of urban genotypes. Examples exist of both single and multiple origins of urban genotypes, but these have proven difficult to generalize. Here, we address genetic differentiation among urban populations, among rural populations, and between urban and rural populations. We conducted an extensive population genomic double digest restriction-site associated DNA sequencing analysis of two non-model grassland lepidopterans, *Coenonympha pamphilus* and *Chiasmia clathrata*, across Europe. The genetic population structures of the study species were strikingly different: *Co. pamphilus* showed strong population differentiation, while this was almost absent in *Ch. clathrata*, which instead showed signs of high current and past gene flow among populations. Results of *Co. pamphilus* are consistent with multiple origins of urban populations, and multiple origins seem plausible also in *Ch. clathrata*. These results suggest that past and large-scale population dynamics need to be integrated into urban evolution research, because population history affects urban evolutionary dynamics.

## Introduction

Urbanisation changes both the abiotic and biotic environment globally [1,2], and it is projected to extensively continue [2,3]. Consequently, wildlife is increasingly forced to face urban environments. Although urban settings filter out many species, other species do cope with urbanisation, and some even successfully exploit urban habitats [4-8]. Because both biotic interactions and abiotic conditions differ in urban environments from those of natural or rural habitats, it is likely that populations in urban habitats experience different selection pressures compared to those living in non-urban habitats. Accordingly, evidence of evolved adaptations to urban environments is accumulating (reviewed by [9-12]). Yet, urban populations are often small and isolated in green patches embedded within an urban matrix, where stochastic events such as habitat modification and destruction by human activities may be more frequent than in other contexts [13]. These conditions make genetic drift potentially an important evolutionary force too, resulting in genetic differentiation not attributed to selection between urban and rural populations [10,14]. Consequently, genetic differentiation of urban populations has been reported in some species [15-20] but not in all studied ones [19,21-24]. However, studies have typically focused on only a single or a few cities (but see [15,18,20,23]), which is why the large-scale dynamics of urban evolution remain insufficiently understood.

An important issue concerning evolution in urban environments is to identify whether urban genotypes have arisen only once and then spread across cities—which is a plausible mechanism in mobile species—or whether urban genotypes derive from independent evolution in different cities [20,22,23,25]. An example of the former scenario is the recent urbanisation-associated range expansion of the butterfly *Pieris mannii* in Central Europe, driven by a single genetic lineage [26]. Industrial melanism in the moth *Biston betularia* in Britain has been traced to a single evolutionary origin too [27]. On the contrary, genetic evidence suggests multiple origins of urban Blackbird (*Turdus merula*) genotypes [15], although the observed timing and spatial spread of urban colonisations erroneously implied a scenario where urban genotypes have a single origin and have then spread from one city to another [28]. In Great tits (*Parus major*), haplotypes under selection in urban populations mainly differ across cities, suggesting that urban adaptations evolve independently and repeatedly in different cities, although selection consistently acts on the same behaviour-linked genes in different urban populations [20]. The owl *Athene cunicularia* shows a clear genetic signature of multiple urban colonisations [29] and repeated adaptation in neural function likely affecting cognition [25]. Finally, in the frog *Lithobates sylvaticus*, bumblebee *Bombus lapidarius*, and damselfly *Ischnura elegans*, urban populations are not genetically differentiated from nearby rural ones, although a few genetic markers associated with urbanisation were identified [21,22,24], rendering inferences on the dynamics of urban evolution impossible. Owing to this diversity of patterns, we cannot draw generalizations about the origin and spread of urban genotypes or the genetic differentiation between urban and rural populations. However, a diversity of patterns is expected across species [14]. Hence, it remains unclear if any generalizations are possible at all. Spatially extensive studies including several cities, and replicated across species are, therefore, necessary for building understanding of the population genetic consequences of urbanisation.

Here, we conducted an extensive population genomic analysis across multiple European urban and rural populations of the butterfly *Coenonympha pamphilus* (L.) (Nymphalidae) and the moth *Chiasmia clathrata* (L.) (Geometridae) to find out the scale of genetic differentiation among urban and rural populations, and to address the origin of urban genotypes. Both species are small and sedentary lepidopterans, inhabit open meadow and grassland habitats in their (Western) Palearctic range and use very common and abundant larval host plants (grasses by *Co. pamphilus*, leguminous plants by *Ch. clathrata*) [30-37]. The species have similar phenologies and produce one generation per year in the northern parts of their distribution and two or more at lower latitudes where the growing season is longer [32,35,36]. Because there is evidence of genetic population structure at a small spatial scale in *Co. pamphilus* [38], we expected to find genetic differentiation also at a larger geographic scale and between urban and nearby rural populations. In *Ch. clathrata*, common garden experiments have demonstrated genetically-based urban adaptations in seasonal plasticity and heat tolerance [39-41] as well as local life-history adaptations along a subset of the climatic gradient included in the present study [42,43], suggesting genetic differentiation among populations.

Besides testing for genetic population differentiation and the genetic signature of urban adaptation, we also aimed to reveal whether urban genotypes all share a single evolutionary origin, or have each derived from a nearby rural population, or display an intermediate pattern between these extremes. The extreme case of a single origin of urban genotypes would manifest itself as shared genetic lineages across urban populations, with individuals from all urban populations belonging to the same genetic cluster, resulting in weaker isolation-by-distance among urban than among rural populations [15]. This pattern could arise if a single urban-adapted lineage colonizes several cities but fails to successfully interbreed with rural populations, either through maladaptation or by failing to reach these populations, such as in the case of human assisted dispersal [44-46]. Uniform selection pressures across urban environments (e.g., due to urban heat islands [23,41], recurrent anthropogenic disturbances [13], or similar cognitive and behavioural demands [20, 25]), combined with a high dispersal capacity, could result in such a pattern. At the other extreme, where each urban population is derived from a nearby rural population, urban individuals would cluster with geographically nearby rural ones. This scenario is possible even if all urban populations experience similar selection pressures because repeated adaptation may lead to similar urban phenotypes even if the underlying genetic changes differ among urban populations (see e.g. [20]). Under this scenario, isolation-by-distance would be similar across both urban and rural populations, and there would be a threshold-type increase in isolation by the degree of urbanisation of the environment.

Under intermediate scenarios, urban genotypes would display multiple origins but fewer than the total amount of sampled cities, which would be best observable by some urban populations clustering with other urban populations, but not all urban populations clustering together.

## Methods

### Sampling

We sampled *Co. pamphilus* with a hand net from 22 sites across Europe. *Ch. clathrata* was sampled from 28 sites across Europe, mainly with a hand net but a light trap was used in all Austrian sites, in two Czech sites (BIS, CHL; see Table S1 for an abbreviation key), and in two Estonian sites (KAR, TAR). Sampling sites for both species consisted almost exclusively of pairwise urban and rural locations associated with the same city (Fig. 1; Table S1).

**Figure 1.**
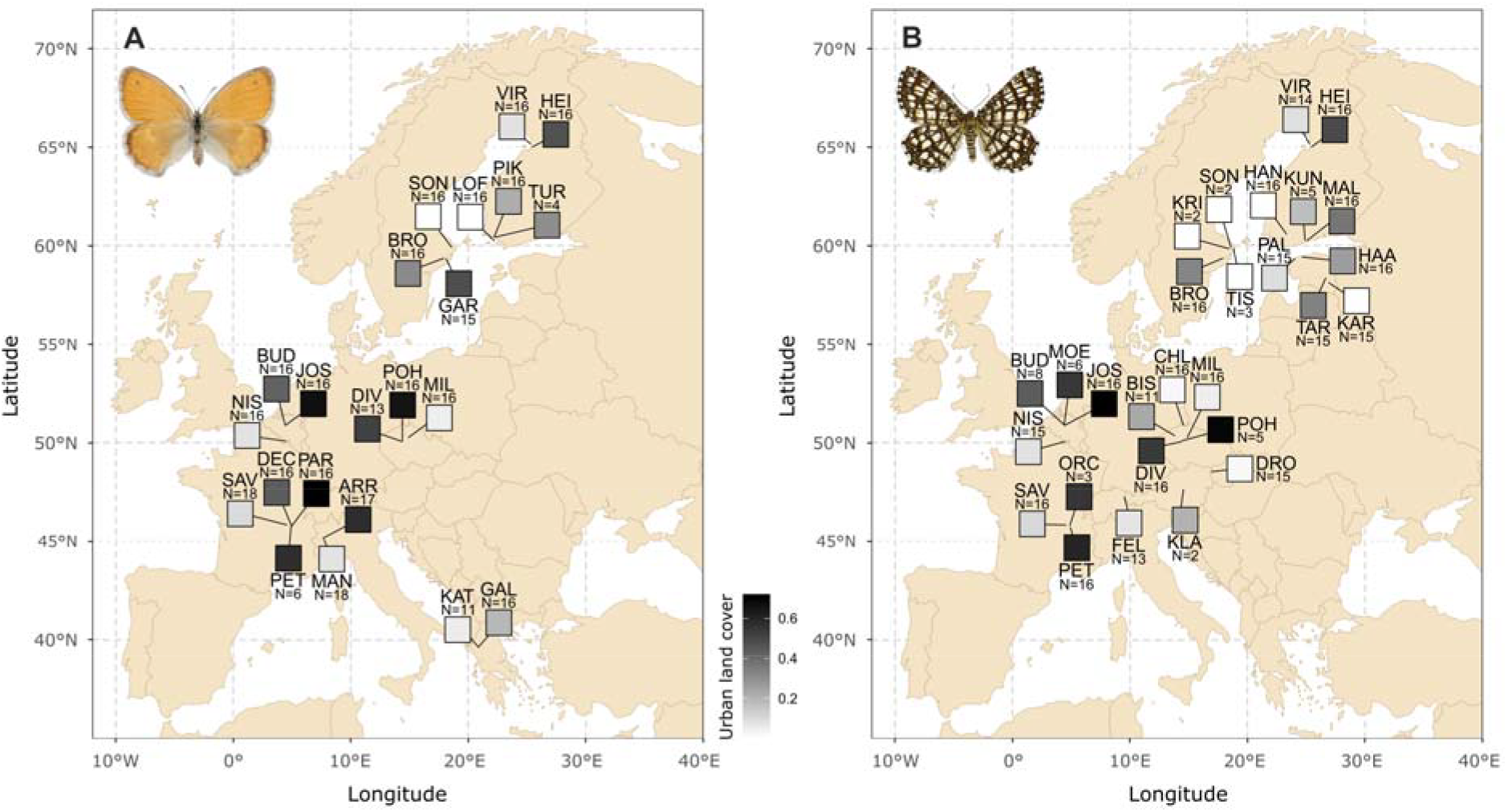
Maps showing the sampling sites for *Coenonympha pamphilus* (A) and *Chiasmia clathrata* (B). Squares indicate sampling sites, with lines connecting them to the exact sampling locations (the squares are spread out to improve readability). The colour of each square indicates the degree of urbanisation at the sampling sites (defined as the proportion of land covered by impervious structures within a 2500-m radius; darker colour means higher urbanisation). Population abbreviations and sample sizes (N) are given for each site. Pictures of the study species are shown at the top left corners of the maps (pictures sourced from the Finnish Biodiversity Information Facility [http://laji.fi] under the CC BY-SA 4.0 license, taken by Pekka Malinen, and accessed on 28 March 2024).

### DNA extraction and ddRADseq library preparation

Genomic DNA (gDNA) was extracted from leg and thorax tissues using the E.Z.N.A. Tissue DNA Kit (Omega Bio-tek), following the manufacturer’s protocol. The gDNA concentration was measured with a Quant-iT PicoGreen dsDNA assay kit (Molecular Probes). The ddRADseq [47] library preparation followed the protocol described by Lee et al. [48] with minor adjustments: PstI and MseI restriction enzymes (NEB) were used for digestion, and the size distribution was assessed via Bioanalyzer (Agilent Technologies). The final libraries were sequenced in two batches on the Illumina HiSeq X PE150 platform at Macrogen (Seoul, South Korea) and NovaSeq X Plus PE150 at Novogene (Cambridge, UK).

### Bioinformatics of ddRADseq reads

Raw paired-end reads were demultiplexed using their unique index and adapter sequences, with no mismatches tolerated, with *ipyrad* v.0.9.92 [49]. The quality of the raw demultiplexed reads was checked with FastQC (available at http://www.bionformatics.babraham.ac.uk/projects/fastq/). Poor-quality samples were removed after quality checks, with one sample excluded from the *Co. pamphilus* dataset, leaving 326 samples, and two samples excluded from the *Ch. clathrata* dataset, leaving 325 samples. The demultiplexed paired reads were merged into single-end reads using PEAR v.0.9.8 [50] with default settings and then processed through the *ipyrad* pipeline for further analysis.

All *ipyrad* analyses were performed on the Puhti supercomputer at the Centre for Scientific computing Finland (CSC), using default settings with the following modifications: datatype was set to “ddrad”, assembly method to “denovo”, restriction overhang to “TGCAG,TAA”, maximum low-quality bases to 6, minimum statistical depth to 8, clustering threshold to 0.9, and the minimum number of samples required for a locus set to 120. Loci with excessive shared polymorphic sites (>50% of samples) were excluded to avoid clustering of paralogs. We generated two types of final matrices: one including all variable sites (SNP) and another containing one random SNP from each putatively unlinked locus (uSNP).

### Data analyses

First, we ran a frequentist population admixture analysis by using sparse non-negative matrix factorization algorithms as implemented in the function ‘snmf’ from the package ‘LEA’ [51] in R version 4.3.2 [52]. The genotypic matrix was used as an input for the derivation of individual-specific ancestry coefficients, and we repeated the analysis with the number of genetic clusters (*K*) spanning from one to 16 in *Co. pamphilus* and from one to eight in *Ch. clathrata*. Higher values of *K* were included in *Co. pamphilus* because initial investigations indicated strong population structure in this species.

We further investigated population structure by employing SplitsTree v.4.19.2 to generate an unrooted genetic network using the Neighbor-Net algorithm with uncorrected p-distances [53]. For admixture histories, we applied population tree inference model implemented in TreeMix v.1.13 [54]. The analysis, performed on the uSNP dataset, progressively added up to eight migration events. Asymmetries in the covariance matrix of allele frequencies, relative to the ancestral population inferred from the ML trees, were used to determine the directionality of gene flow. Demographic history was investigated with the stairwayplot2 software [55,56] by running the analysis separately for each cluster suggested by the population admixture analysis. We derived site frequency spectra from the genotypic matrix where missing values were imputed based on the population admixture analysis. Sequence length was 139579, generation length 0.5 years, and the default settings were otherwise used.

For a further investigation of the weak population structure in *Ch. clathrata*, we derived a phylogenomic maximum likelihood (ML) tree for this species and investigated its demographic history. The tree was constructed by using the SNP dataset in IQ-TREE v.2.1.3 [57]. The integrated ModelFinder [58] within IQ-TREE tested 286 DNA substitution models, selecting ‘SYM+I+G4’ for the *Ch. clathrata* dataset. To reconstruct the phylogeny, ML analysis with ultrafast bootstrap approximation model (1,000 replicates) was applied [59]. The resulting tree was generated using FigTree v.1.4.2 [60] and further modified using CorelDRAW v24.

To get more insight into population differentiation and the role of urbanisation and climate in it, we ran a redundancy analysis (RDA) by using the R function ‘rda’ from the package ‘vegan’ [61] for both species and setting urbanisation and growing season length as environmental covariates, and conditioning for latitude and longitude (i.e. spatial structure). We measured urbanisation as the proportion of land covered by artificial surfaces and constructions (water bodies excluded) within 500-m, 2500-m, and 5000-m radii based on 10 m resolution land cover data from S2GLC Europe from 2017 (version v1.2; [62]). The radius of 2500 m best captured the urbanization gradient (maximized the range of derived values for proportion of urban land cover), so it was used in all analyses. Growing season length was measured as the number of days during which mean daily temperature was above 5 °C, with the conditions that mean daily temperature was above the threshold for seven consecutive days since the beginning date of the season and below the threshold for seven consecutive days after the end date of the season. We used interpolated (30[arc[second resolution) daily temperatures for 1990–2005 from CHELSA Europe (version v1.1; [63,64]) for the estimation of site-specific growing season lengths. Both urbanisation and growing season length were centred and scaled to mean of zero and standard deviation of one by subtracting the mean and dividing by standard deviation. Before the RDA analysis, we imputed missing values in the genotypic matrices by using the function ‘impute’ from the R package ‘LEA’ [51] and setting the number of genetic clusters to the value that was best supported in the snmf-based admixture analysis (nine clusters in *Co. pamphilus*, three in *Ch. clathrata*; see Results). We ran permutation tests for the effects of urbanisation and growing season length by using the ‘anova.cca’ function [61] with 1000 permutations.

Furthermore, we ran a genotype-environment association analyses with the latent factor mixed model approach to investigate whether any SNPs show an association with either urbanisation or background climate, measured as growing season length. We used the R function ‘lfmm2’ from the package ‘LEA’ [51] for fitting the latent factor mixed models. For both species, the response variables were the columns of the imputed genotypic matrix, and the environmental covariates included the above-mentioned urbanisation and growing season length metrics standardized to mean of zero and standard deviation of one. The number of latent factors was initially set to be equal to the number of genetic clusters used in missing value imputation, but we also fitted models with number of latent factors spanning from 7 to 15 in *Co. pamphilus* and from 2 to 5 in *Ch. clathrata* to assess if genomic inflation factors, or the distribution of *p*-values (a model goodness-of-fit indicator), depended on the number of latent factors. We report results for the models fitted with the numbers of latent factors that resulted in best model goodness-of-fit, yet inferences remained unaffected by the numbers of latent factors within the investigated ranges. We derived *p*-values separately for urbanisation and growing season length by using the ‘lfmm2.test’ function [51]. First, we adjusted the SNP-specific *p*-values of both covariates to population structure. We did this by calculating the genomic inflation factors (λ) as λ_*i*_ = median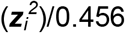, where ***z*** (*i* = {urbanisation, growing season length}) refers to a vector of *z*-scores of covariate *i*, and then deriving *p*-values for these approximately 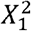 -distributed scores (see [65]) from the 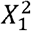 distribution. Second, we made an adjustment for multiple testing by using the Benjamini-Hochberg procedure [66] so that we derived statistical significance thresholds (adjusted *p*-values) corresponding to 1% and 5% false discovery rates. For inferences, we chose the model where the number of latent factors resulted in λ_*i*_ values closest to unity and appropriate distributions of *p*-values in both species.

Finally, we conducted isolation by distance (IBD) and isolation by (urbanisation of the) environment (IBE) analyses. For this purpose, we derived matrices of pairwise F_ST_ values, pairwise geographic distances (in kilometres), and pairwise differences in urbanisation between all populations for both species. Pairwise F_ST_ values (Tables S2 and S3) were calculated with Arlequin v.3.5.1.3 [67], employing 1000 permutations. In the IBD analysis, we tested if the pairwise F_ST_ values between populations are positively correlated with pairwise geographic distances between the populations. In the IBE analysis, we tested if the pairwise F_ST_ values between populations are positively correlated with pairwise differences in urbanisation or growing season length between the populations. By following Rousset [68], we scaled the proxies of genetic distance (F_ST_ values) as F_ST_/(1 - F_ST_), but we did not use a logarithmic transformation for geographic distances, because untransformed distances best linearized the relationship between genetic and geographic distances. We also used the raw differences in urbanisation as the distance measures in the IBE analysis [68]. We tested whether genetic distances were correlated with geographic (IBD) or urbanisation (IBE) distances separately for rural-rural and urban-urban population pairs with a Mantel test (based on 5000 permutations) to test the predictions concerning origin of potential urban-specific genotypes by using the R function ‘mantel’ from the package ‘vegan’ [61]. We used an urbanisation value of 0.2 (proportion of urban land cover within a 2500-m radius) as a cutoff when classifying populations as rural (urbanisation <0.2) or urban (urbanisation ≥0.2) in this analysis, which correctly categorized the locations as rural or urban along with the subjective categorization done during fieldwork.

## Results

48459 polymorphic SNPs for 326 *Co. pamphilus* individuals met our quality criteria for inclusion (see Methods), resulting in 41.1% missing data (Table S4). The corresponding numbers for *Ch. clathrata* were 17896 polymorphic SNPs for 325 individuals, and 37.8% missing data (Table S5). There was a small difference in the proportion of missing data per individual between the two sequencing batches in *Co. pamphilus* (95% bootstrapped percentile confidence intervals [CI] of the means; batch 1: 0.426 [0.414, 0.438]; batch 2: 0.391 [0.381, 0.402]), but not in *Ch. clathrata* (batch 1: 0.375 [0.351, 0.402]; batch 2: 0.378 [0.363, 0.395]). Despite being statistically supported, the difference (3.43 percentage points) in *Co. pamphilus* is so small that it is very unlikely to affect results.

Population admixture analysis identified nine genetic clusters as the best-supported number of clusters (*K*) in *Co. pamphilus*, yet numbers from seven to 14 were almost equally well-supported (Fig. 2A; see Fig. S1 for *K*=5—8). In this species, country-level genetic differentiation was evident, except that the Czech and Greek populations were assigned to the same cluster when *K*≤10 (Fig. 2A). This country-level genetic clustering was also recovered by a network analysis by SplitsTree (Fig. S2A). TreeMix (a tool designed for detecting historical population splits and gene flow among populations) analysis yielded similar results, with the only inferred migration events having taken place from the French to the Czech lineages, and from the Italian to the French lineages (Fig. S3A). Within-country genetic clustering was found in Finland where the northern and southern populations formed separate clusters (Fig. 2A; Fig. S2A). Urban populations generally clustered with nearby rural populations (Fig. 2A), as expected under a scenario of multiple origins of urban genotypes. However, in Belgium, and to a lesser degree in Czech Republic, some urban populations showed an ancestry different from the associated rural populations.

**Figure 2.**
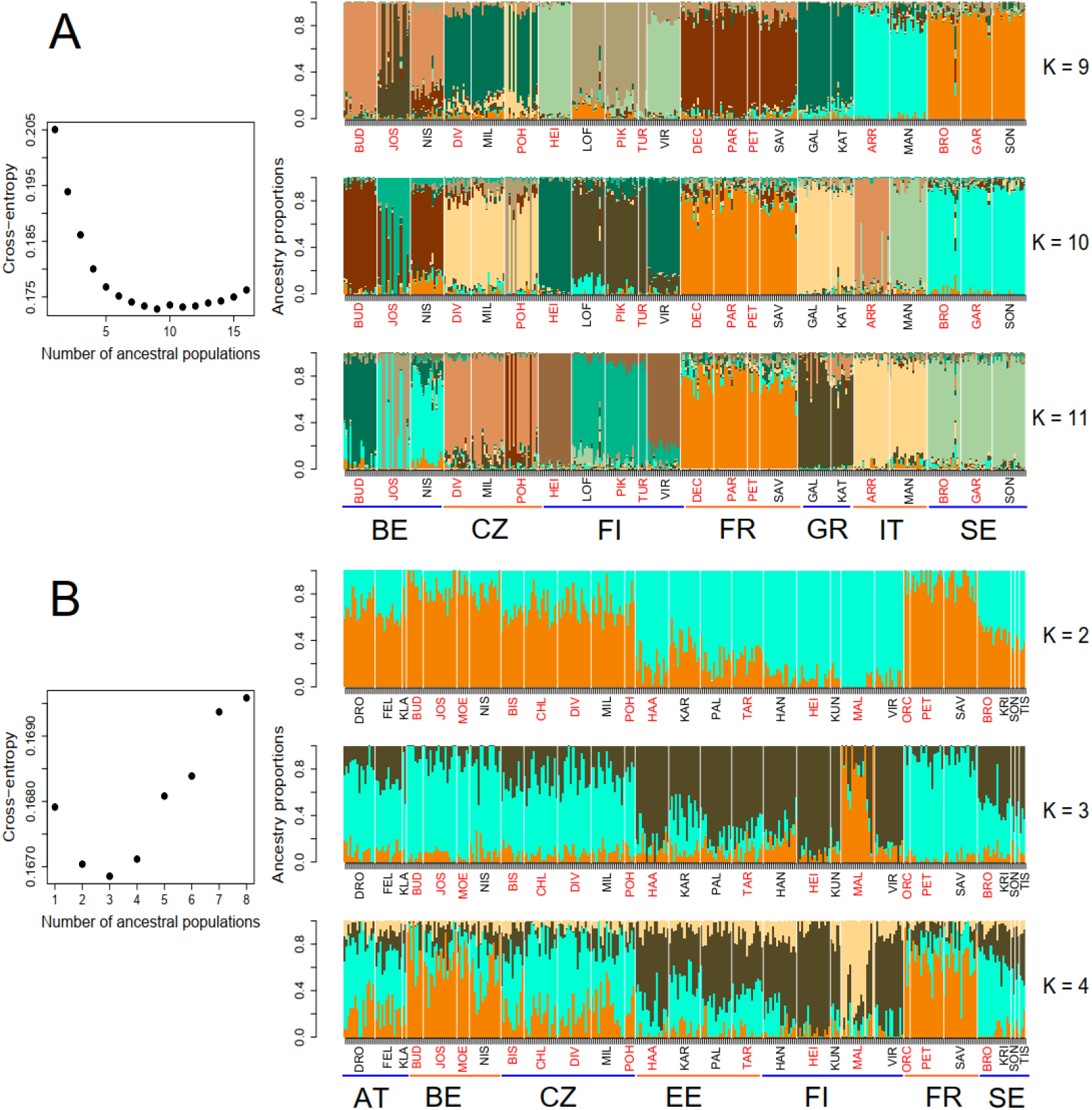
Results of population admixture analysis. Cross-entropy values (left column) and individual-specific ancestry coefficients (coloured vertical bars in the right column) derived from an analysis using sparse non-negative matrix factorization algorithms for *Coenonympha pamphilus* (A) and *Chiasmia clathrata* (B). The number of ancestral populations (*K*) that minimizes cross-entropy is the best-supported number of genetic clusters (nine in *Co. pamphilus*; three in *Ch. clathrata*). The right column shows the ancestry coefficients for three well-supported numbers of ancestral populations (*K* = {9, 10, 11} for *Co. pamphilus, K* = {2, 3, 4} for *Ch. clathrata*). The height of each colour within a vertical bar indicates the probability that the individual in question belongs to a particular genetic cluster. The order of individuals is kept the same across panels. White vertical lines separate populations, and population labels (see Fig. 1; Table S1) are given below the horizontal axis. Country abbreviations (see Table S1) are also given below the population labels in the bottom panels of both species, and the coloured horizontal lines above the country labels tell apart the populations from different countries. A rough rural-urban categorisation of the populations is indicated with font colour (black for rural [urban land cover <20%]; red for urban [urban land cover ≥20%]).

In *Ch. clathrata*, the genetic clustering analysis best supported the presence of three genetic clusters (Fig. 2B). One of the clusters consisted of individuals of the urban Malmi population from Southern Finland, another one of all the other Finnish and Estonian populations, and the third of all the remaining populations (Fig. 2B). The assignment of individuals to these clusters was not as clear as in *Co. pamphilus*, and the ancestry of many individuals was unclear. This weak population structure is not due to confounding effects of several populations with a low sample size, as a sensitivity analysis with these low-sample-size populations removed, recovered similar results (Fig S4). Lack of genetic population structure was also highlighted in a SplitsTree analysis (Fig. S2B) and in the ML tree (Fig. S5). Consequently, drawing inferences concerning the origin of urban genotypes is difficult in this species. Nonetheless, a TreeMix analysis recovered a geographically meaningful clustering, although the degree of genetic differentiation was minimal in comparison to *Co. pamphilus*, and several migration events between lineages were inferred (Fig. S3B). Investigation of historical population size changes with stairwayplot2 revealed a long history with a very high effective population size, except in the Malmi population (Fig. S6), suggesting high gene flow among populations.

To get more insight into population differentiation and the associated role of urbanisation and background climate, we ran a redundancy analysis (RDA). In *Co. pamphilus*, urbanisation and growing season length explained 3.71% of variation in the genotypic matrix (adjusted *R*^2^) when conditioning for spatial structure, with both urbanisation (*F*_1,321_=2.94, *p*=0.001) and growing season length (*F*_1,321_=13.7, *p*=0.001) being significant predictors of genotypic variation. Populations were clearly separated in the space defined by the first two RDA axes, with urban populations situated close to nearby rural ones (Fig. 3A). This indicates a strong population structure, aligning with the results of the population admixture analysis.

**Figure 3.**
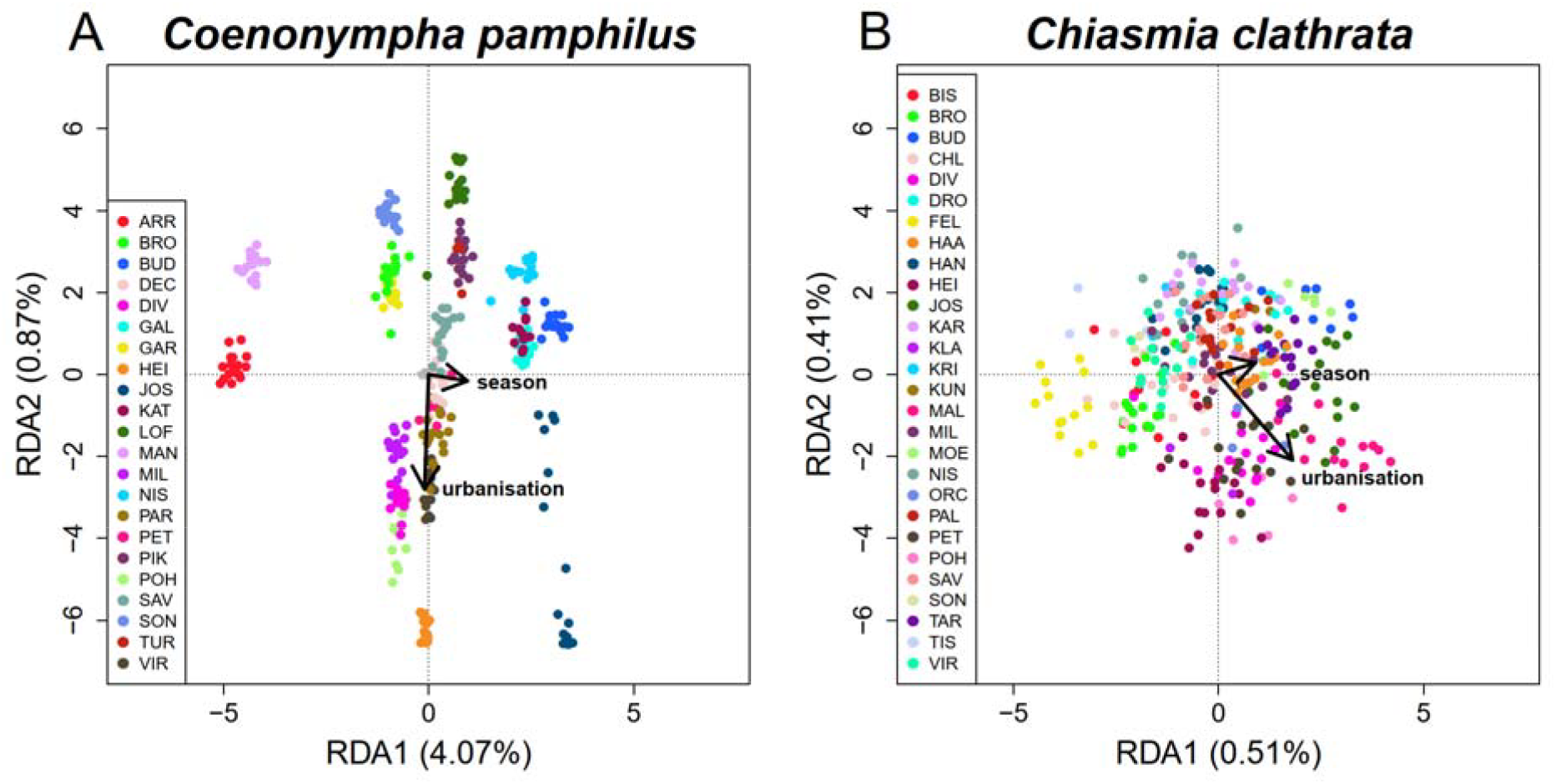
Visualization of results of the redundancy analysis (RDA) exploring the effects of urbanisation and climate (growing season length) on genotypic variation in *Coenonympha pamphilus* (A) and *Chiasmia clathrata* (B). Scaling is such that the lengths of the arrows—illustrating directions of increasing urbanisation and growing season length—show the effect sizes (arrow lengths are multiplied by three to improve readability), while the angles between the arrows indicate the correlation between the environmental predictors (almost no correlation here, because the angle is close to 90° in both species). The small grey points, extremely densely concentrated in the middle, depict the locus scores, while the coloured large points indicate the individual scores, with colours corresponding to the population of origin. Proportions of variance explained by the RDA axes are given in the axis labels. See Figure 1 and Table S1 for population abbreviations. Corresponding visualizations, scaled so that the distances among points depend on their similarity, are presented in Figure S7.

In *Ch. clathrata*, urbanisation and growing season length explained 2.94% of the genotypic variation in the RDA analysis (adjusted *R*^2^). Nonetheless, both urbanisation (*F*_1,320_=1.47, *p*=0.001) and growing season length (*F*_1,320_=1.52, *p*=0.001) were again significant predictors of this genotypic variation. Populations were not separated in a space defined by the first two RDA axes but were largely overlapping (Fig. 3B). This aligns well with the results of the population admixture analysis, where clear genetic clusters could not be identified.

Because a part of the genetic variation was associated with variation in urbanisation and climate in the RDA analysis, we ran a more detailed investigation of associations between SNPs and either urbanisation or growing season length; genotype-environment association analysis with the latent factor mixed model approach. In *Co. pamphilus*, the number of latent factors minimally affected model goodness-of-fit but it did not affect inferences. The model including 12 latent factors appeared to be best (see Fig. S8A, B) and we used it for drawing inferences concerning this species. That model found three candidate SNPs associated with variation in urbanisation with a 5% false discovery rate (FDR), none of which were significant at 1% FDR (Fig. 4A). In contrast, 4136 SNPs were associated with variation in growing season length at 5% FDR, 2547 of which were still significant at 1% FDR (Fig. 4B). Hence, there is only a weak signature that few of the investigated SNPs are involved in urban adaptation. However, there is a clear signal of climate adaptation across numerous loci.

**Figure 4.**
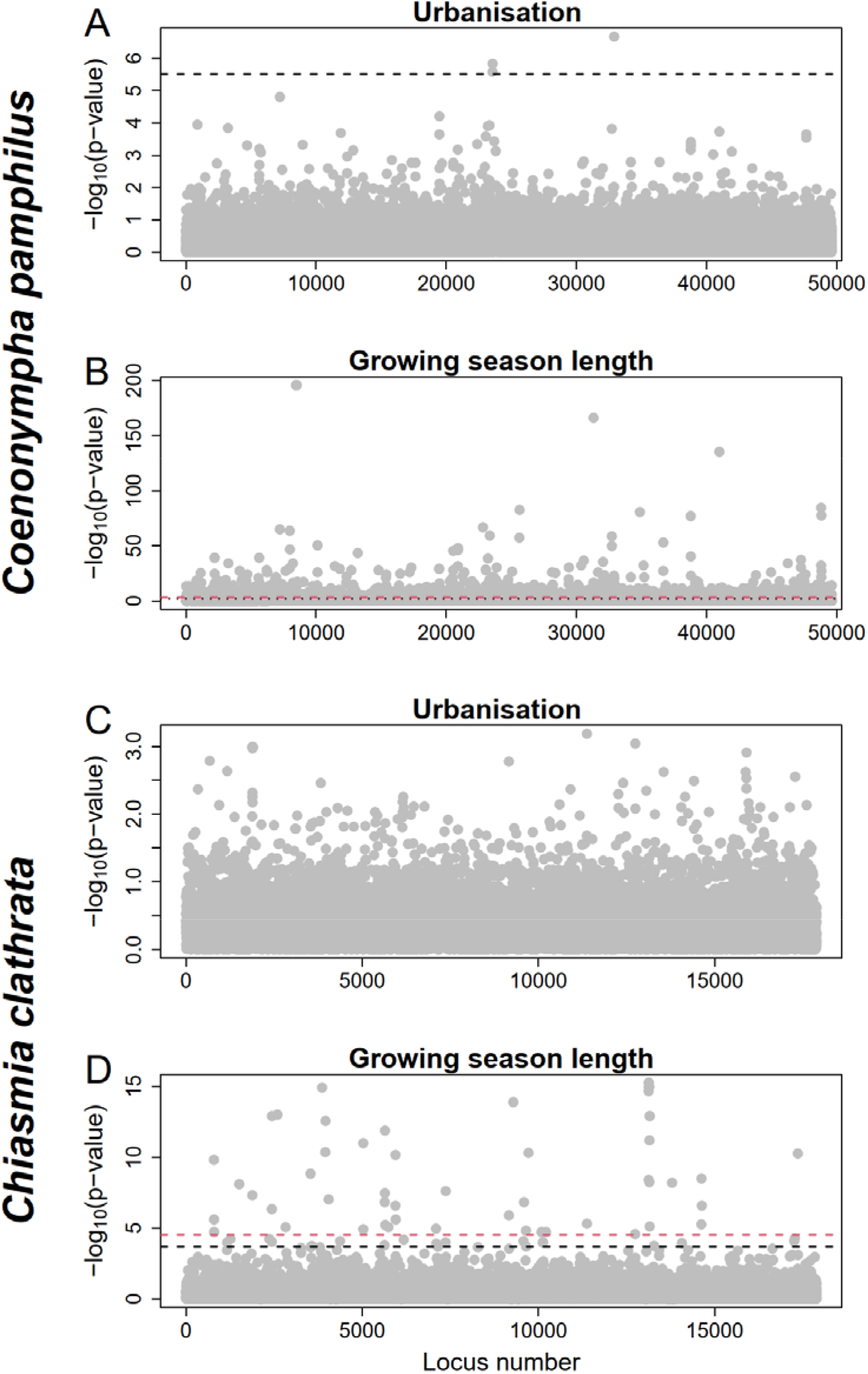
Manhattan plots showing the log_10_-transformed *p*-values for SNP-specific associations with urbanisation (A, C) and growing season length (B, D) in *Coenonympha pamphilus* (A, B) and *Chiasmia clathrata* (C, D). The red and black dashed lines indicate statistical significance thresholds adjusted for 1% and 5% false discovery rate, respectively (each threshold shown only when at least one SNP exceeds the given threshold). Note that the scale of the y-axis varies greatly among panels A–D.

In *Ch. clathrata*, the number of latent factors similarly had a minimal effect on model goodness-of-fit and no effect on inferences. The model including four latent factors seemed best (see Fig. S8C, D). Hence, we used that model for drawing inferences concerning this species. While there were no SNPs associated with variation in urbanisation, neither at 1% nor at 5% FDR (Fig. 4C), 73 SNPs were associated with variation in growing season length at 5% FDR, 53 of which remained significant at 1% FDR (Fig. 4D). *Chiasmia clathrata*, therefore, shows some genetic signature of climate adaptation, but no indication of urban adaptation. This is consistent with the known existence of local climate adaptations in this species but contradicts with the known presence of urban adaptations.

Finally, to test the predictions concerning single versus multiple origins of urban populations, we conducted isolation-by-distance (IBD) and isolation-by-(the degree of urbanisation of the) environment (IBE) analyses. We performed these analyses separately for rural and urban populations. In *Co. pamphilus*, there was clear IBD both among rural and urban populations (Fig. 5A) as well as across all possible population comparisons (Mantel test: *r*=0.667, *p*=0.0002), but IBE was neither present in these population comparison categories (Fig. 5C) nor overall (Mantel test: *r*=-0.0313, *p*=0.62). There was also no threshold-type increase in genetic distance at any urbanisation difference (Fig. 5C). Hence, the patterns in *Co. pamphilus* match most of the predictions for multiple origins of urban populations.

**Figure 5.**
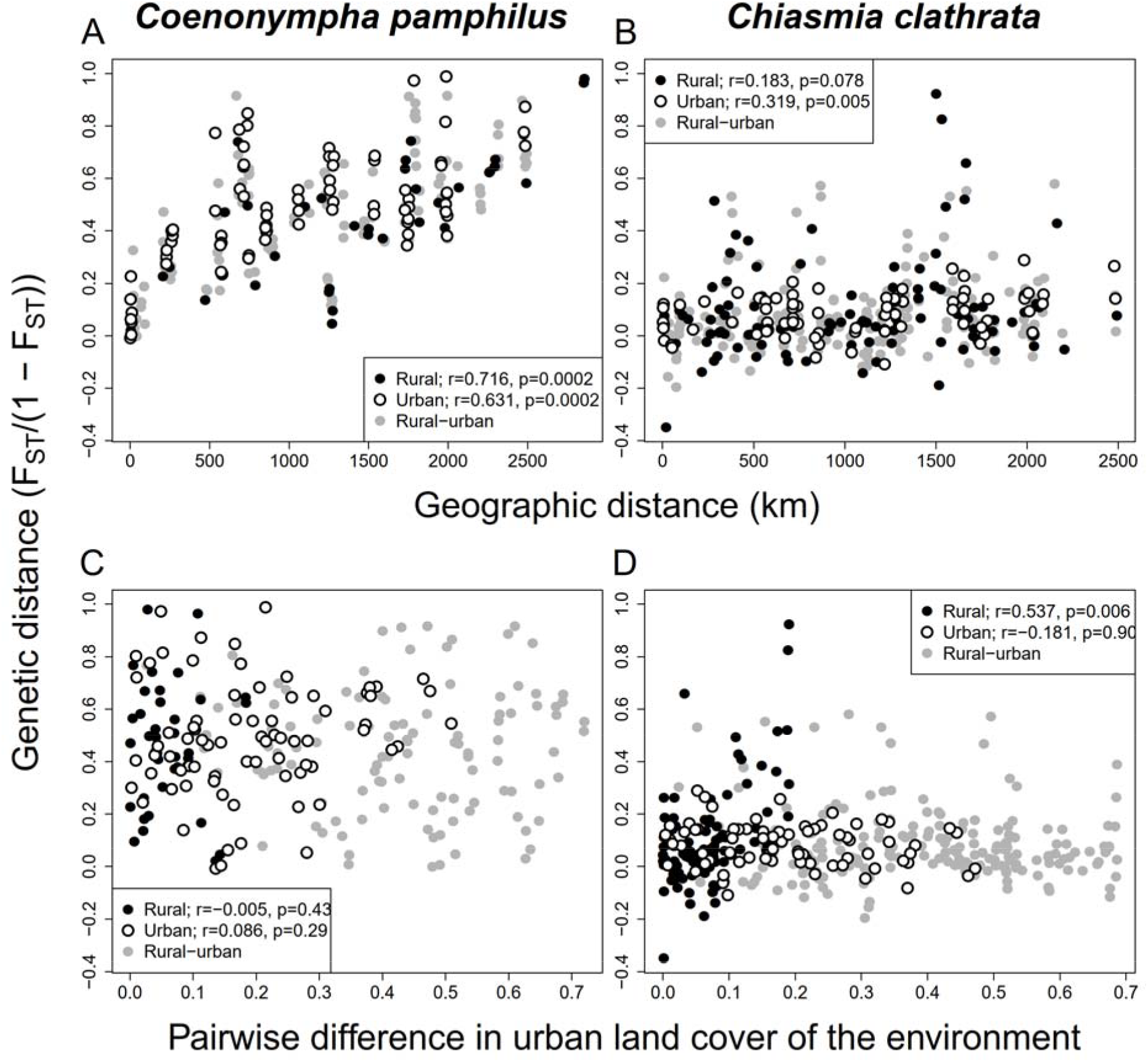
Isolation-by-distance (A, B) and isolation-by-environment (C, D) analyses for *Coenonympha pamphilus* (A, C) and *Chiasmia clathrata* (B, D). Genetic distances were based on transformation by Rousset [68], whereas difference in the urbanisation of the environment is the inter-population difference in proportion of land covered by impervious surfaces within 2500-m radii from the sampling locations. Black dots indicate rural-rural population comparisons, open circles urban-urban comparisons, and grey dots rural-urban comparisons. Pearson product-moment correlation coefficients, estimated based on Mantel tests with 5000 permutations, and associated *P*-values, are indicated in the legends for rural-rural and urban-urban comparisons. Note that the scale of the y-axis is kept constant across all panels to facilitate comparison between the species.

In *Ch. clathrata*, there was slight IBD among urban populations (Fig. 5B) and across all population comparisons (Mantel test: *r*=0.190, *p*=0.0036) but not when comparing only rural populations (Fig. 5B). In this species, there was also some IBE when comparing rural populations but neither among urban populations (Fig, 5D), nor across all population comparisons (Mantel test: *r*=-0.0320, *p*=0.59). These patterns contradict the predictions of both single and multiple origins of urban populations.

## Discussion

We found contrasting genetic population structures in two grassland lepidopterans. Such a difference between strong genetic population structure in *Co. pamphilus* and almost no structure in *Ch. clathrata* is striking, especially when considering that common-garden experiments on *Ch. clathrata* have demonstrated heritable phenotypic differences between nearby rural and urban populations [39-41] and among populations along a climatic gradient [42,43]. Genetic population structure in *Co. pamphilus* is also highlighted by relatively low effective population size estimates (Table S6) and is clearly consistent with multiple origins of urban genotypes. The lack of population structure in *Ch. clathrata* precludes rigorous inferences on the origins of urban genotypes. However, because the high past (Fig. S6) and current (Table S7) effective population size estimates and the star-like topology of the phylogeny of studied individuals (Figs S2 and S5) suggest high gene flow among all *Ch. clathrata* populations, a single origin of urban populations can be excluded.

Our results concerning *Co. pamphilus* generalize the small-scale population structure found by Wood and Pullin [38] to a continental scale. As the dispersal capacity of *Co. pamphilus* appears limited [30], and as urbanization reduces the species’ observed flight speed [69], population differentiation may eventually evolve even between geographically nearby populations due to genetic drift. We found little evidence of selection in urban populations, as only three SNPs were associated with urbanisation, and only at a less strict 5% FDR threshold. This aligns with a ddRAD study on a damselfly that did not find strong evidence of specific SNPs being under selection in urban environments [24], yet a study on a bumblebee identified urban adaptation in some loci despite weak genetic population structure [21]. Whole-genome analyses revealed clear evidence of loci under selection in urban environments in two bird species, although genetic population structure was weak in one of them [20,25]. These examples suggest that methodological limitations of the ddRAD approach may partially explain why we did not find convincing evidence of urban adaptation in *Co. pamphilus* despite strong evidence of local climatic adaptation.

Despite lacking identifiable SNPs associated with urbanisation, we know from common-garden experiments that adaptations to the urban environment have evolved in *Ch. clathrata* populations. There are five mutually non-exclusive explanations for this conundrum, and these explanations are also equally likely to explain the weak genetic signature of urban adaptation in *Co. pamphilus*. (1) Urban adaptation may be polygenic, with different loci and alleles underlying the observed phenotypic evolution in different urban populations (see e.g., [20,66]), making the association analysis conservative. This is plausible because the known urban adaptations concern polygenic (quantitative) traits in *Ch. clathrata* and many other species (e.g., [20,70-73]). Consistent with this hypothesis, pairwise F_ST_ values between nearby urban-rural population pairs suggest slight differentiation associated with the urban environment in *Co. pamphilus* (mean scaled F_ST_=0.109 [95% confidence interval: 0.0226, 0.195], *df*=7, *t*=2.99, *p*=0.02), yet this is not the case in *Ch. clathrata* (mean scaled F_ST_=-0.0100 [95% confidence interval: -0.0617, 0.0417], *df*=8, *t*=-0.448, *p*=0.67), which has very low population differentiation overall. (2) Low genetic variation between populations may have hindered the identification of adaptive markers. Accordingly, the number of polymorphic SNPs recovered in *Ch. clathrata* was only 37% of those recovered in *Co. pamphilus*. (3) Key SNPs linked to urban adaptation may not have been captured in the sampled data due to the random genome sampling inherent to the ddRADseq approach (see [47]). (4) Stringent bioinformatics quality control may have filtered out important low-quality SNPs, such as those with low read depth or high error rates. Finally, (5) small sample sizes of a few *Ch. clathrata* populations (e.g., KLA, ORC, KRI, SON, and TIS) and sampling imbalances between these and other populations may have constrained the detection of strong associations. Yet, the insensitivity of the population admixture analysis for the exclusion of the low-sample-size populations suggests that this explanation is unlikely. Nevertheless, there were SNPs clearly associated with the known adaptations to local climate in *Ch. clathrata*, indicating that our approach at least had power to detect genotype-environment associations originating from selection that has continued longer and over a larger spatial scale than the selection imposed by the urban environment.

A plausible explanation for the lack of population structure in *Ch. clathrata* is high gene flow, suggested by the high effective population size estimates. Although *Ch. clathrata* is considered as a sedentary species [31], long-distance dispersal could occur if adult moths are carried by strong winds. We consider such wind-assisted dispersal more likely in *Ch. clathrata* than in *Co. pamphilus* because the latter species more typically remains inactive in windy conditions. Yet, our findings also suggest some long-distance gene flow in *Co. pamphilus*, as individuals from the Lofsdal (LOF) population in southern Finland show some ancestry from the Swedish populations, which cannot be explained by gene flow through the continuum of populations around the Bothnian Bay because the northern (HEI, VIR) populations belonging to that continuum formed a separate genetic cluster (Fig. 2A). The TreeMix analysis also suggested migration from western to central Europe and over the Alps (Fig. S3A). Hence, the assumption that both study species are poor dispersers [30,31] needs to be revisited.

Urban populations of both species have likely originated when parts of a formerly uniform population have been trapped in green spaces in the urban matrix once cities have grown, although the trapping is currently incomplete for some studied urban populations. This is a similar process with similar genetic consequences as population subdivision due to habitat fragmentation in the butterfly *Zerynthia polyxena* [74]. However, colonization of urban habitat patches is also possible, either by flying adults or by anthropogenic transportation of juveniles. It is known that dispersal by human transport occurs in the spider *Latrodectus hesperus* [44] and some lepidopteran pests [45,46]. Anthropogenic dispersal remains a possibility for *Ch. clathrata* too, if eggs or larvae are transported with leguminous fodder or mulching, but we consider such dispersal unlikely for *Co. pamphilus*. We also consider wind-assisted long-distance dispersal unlikely for *Co. pamphilus*. Anyway, unlikely or unexpected dispersal possibilities need to be considered, as exemplified by the plant *Lepidium virginicum* where genetic evidence suggests city-to-city colonization over long distances, contrary to *a priori* expectations [75].

Our study adds diversity to the reported patterns of genetic population structure in the urban evolutionary ecology context and highlights that caution is needed when generalizing results based on a few species. The contrasting patterns reported here seem to derive from different levels of inter-population gene flow in the two species, different demographic histories, and different phylogeographic backgrounds. Phylogeography also affects the origin of urban populations, so the biogeographical and historical contexts need to be considered in urban evolution studies, independently of whether the focus is on a single or multiple urban populations. Therefore, continent-scale population genetic analyses should be integrated as a part of the urban evolutionary ecology research program. Currently, there is a paucity of such studies, but the existing ones suggest that urban adaptation occurs and is largely repeated [20,23].

## Data availability

The ddRAD raw demultiplexed reads are deposited at NCBI SRA under BioProject no. PRJNA1177370 and BioSample accession numbers SAMN44419406-SAMN44419730 for *Ch. clathrata* and SAMN44419731-SAMN44420056 for *Co. pamphilus*. The data (bioinformatics output files and environmental data) that support the findings of this study are openly available in Zenodo at https://doi.org/10.5281/zenodo.17162242.

## Code availability

The R-scripts of the analyses are available at Zenodo (DOI: 10.5281/zenodo.17162242).

## Acknowledgements

We are grateful to Ulrich Hiermann, Toni Mayr, Christin Wieser, and Manuel Vilgut for providing us with some *C. clathrata* specimens, to Anna Antinoja, Francesca Cochis, and Irene Piccini for helping us during fieldwork, to Laura Törmälä for carrying out ddRAD library preparation in the laboratory, and to Soile Alatalo and Hannele Parkkinen for assistance in DNA extraction. We thank two anonymous reviewers for constructive feedback that helped to improve the manuscript. We also acknowledge the use of image sources from the Finnish Biodiversity Information Facility (FinBIF/laji.fi). The IT Center for Science (CSC) provided computational resources for bioinformatics and data storage services for the study.

## Author contributions

S.M.K conceived the study. S.M.K., G.C.A., S.B., P.H., T. Kadlec, T. Kankaanpää, G.M., M.E.N., E.P., T.T., K.Z., and T.M. collected samples, with the sampling being coordinated by S.M.K. and T.M., K.M.L. did bioinformatics, and data were analysed by S.M.K., K.M.L., and F.H., with T. Kankaanpää extracting and handling land cover and climate data and helping in data visualization. The manuscript was initially drafted by S.M.K., K.M.L., F.H., and T.M. with input from all the other authors.

## Ethics

National legislation and international regulations were followed in the study. As the study included sampling of non-endangered insects, there was no need for an animal experimentation permit. Estonia, Greece and France were the only countries where sampling required a permit. Sampling in Estonia was allowed by permission numbers 1-3/20/293 and 1-3/22/13, in Greece by a permission number 25518/938, and in France by a permission number TREL2206915S / 574. France was the only country included in the study that had regulations based on the Nagoya Protocol. Export of the sampled genetic material out of France was allowed by the certificate ABSCH-IRCC-FR-260797-1.

## Additional information

Supplementary information file.

