## Supplementary material for "Population history shapes urban evolutionary dynamics: distinct genetic structure across urban and rural Europe in two lepidopterans"

### Supplementary Figures

**Figure S1.** Additional results of the population admixture analysis in *Co. pamphilus*.

**Figure S2.** Phylogenetic relationships among the sequenced individuals.

**Figure S3.** Country-level analysis of population splits and mixture.

**Figure S4.** Sensitivity analysis for the population admixture results in *Ch. clathrata*.

**Figure S5.** Maximum likelihood tree of *Ch. clathrata* individuals.

**Figure S6.** Estimated historical effective population sizes in *Ch. clathrata*.

**Figure S7.** Visualization of results of a redundancy analysis (RDA).

**Figure S8.** Histograms illustrating distributions of *p*-values of SNP-specific tests of association with urbanisation and growing season length.

### Supplementary Tables

**Table S1.** Sampling sites and basic site-specific information.

**Table S2.** Estimated pairwise $F_{ST}$ and Nei's genetic distance values between *Co. pamphilus* populations.

**Table S3.** Estimated pairwise $F_{ST}$ and Nei's genetic distance values between *Ch. clathrata* populations.

**Table S4.** Details of DNA sequencing data for *Co. pamphilus* individuals.

**Table S5.** Details of DNA sequencing data for *Ch. clathrata* individuals.

**Table S6.** Estimated effective sizes of the studied *Co. pamphilus* populations.

**Table S7.** Estimated effective sizes of the studied *Ch. clathrata* populations.

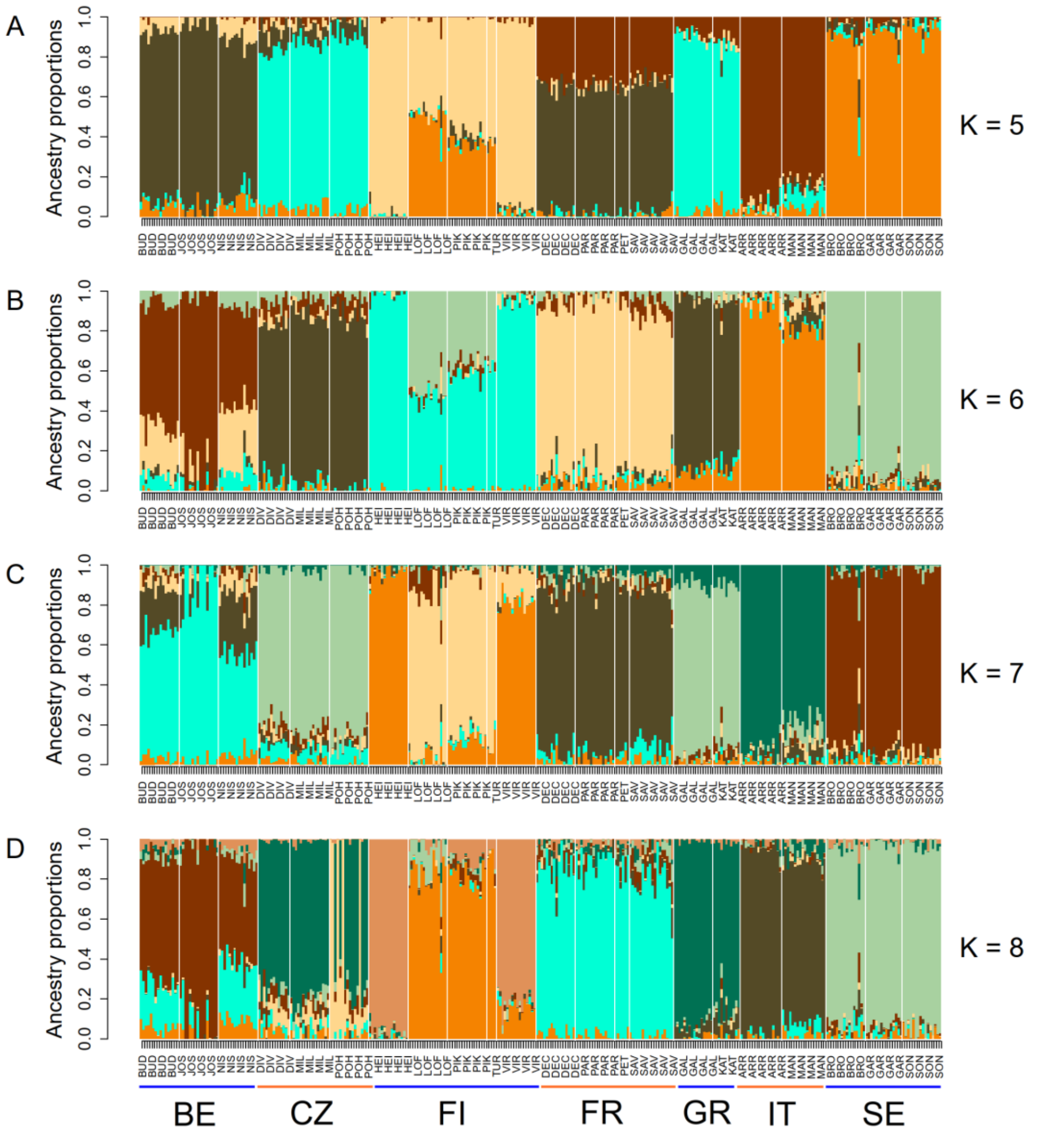


**Figure S1.** Results of the population admixture analysis in *Co. pamphilus* for the numbers of ancestral populations (*K*) being five (A), six (B), seven (C), and eight (D). See Figure 2 for other details.

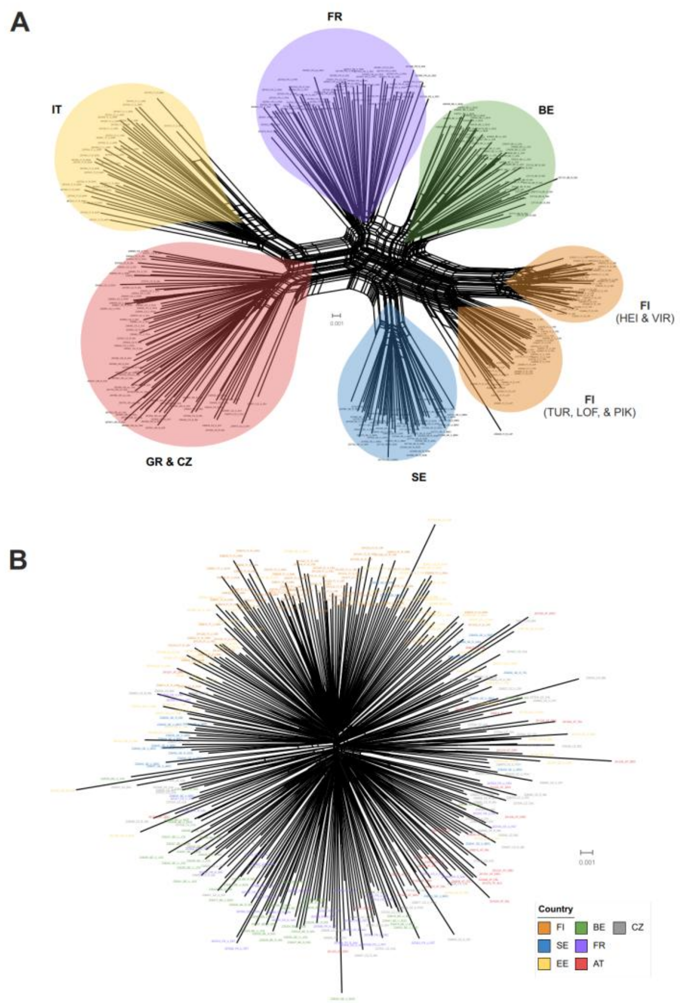


**Figure S2.** Phylogenetic relationships among the sequenced individuals in *Co. pamphilus* (A) and *Ch. clathrata* (B) inferred using a phylogenetic network analysis in the SplitsTree software [1]. The analysis is based on uncorrelated P-distances derived from the SNP dataset. In *Co. pamphilus*, the individuals form clusters that match geographically distinct sampling areas, with the exception that Czech and Greek individuals cluster together. No clustering was found in *Ch. clathrata* in this analysis.

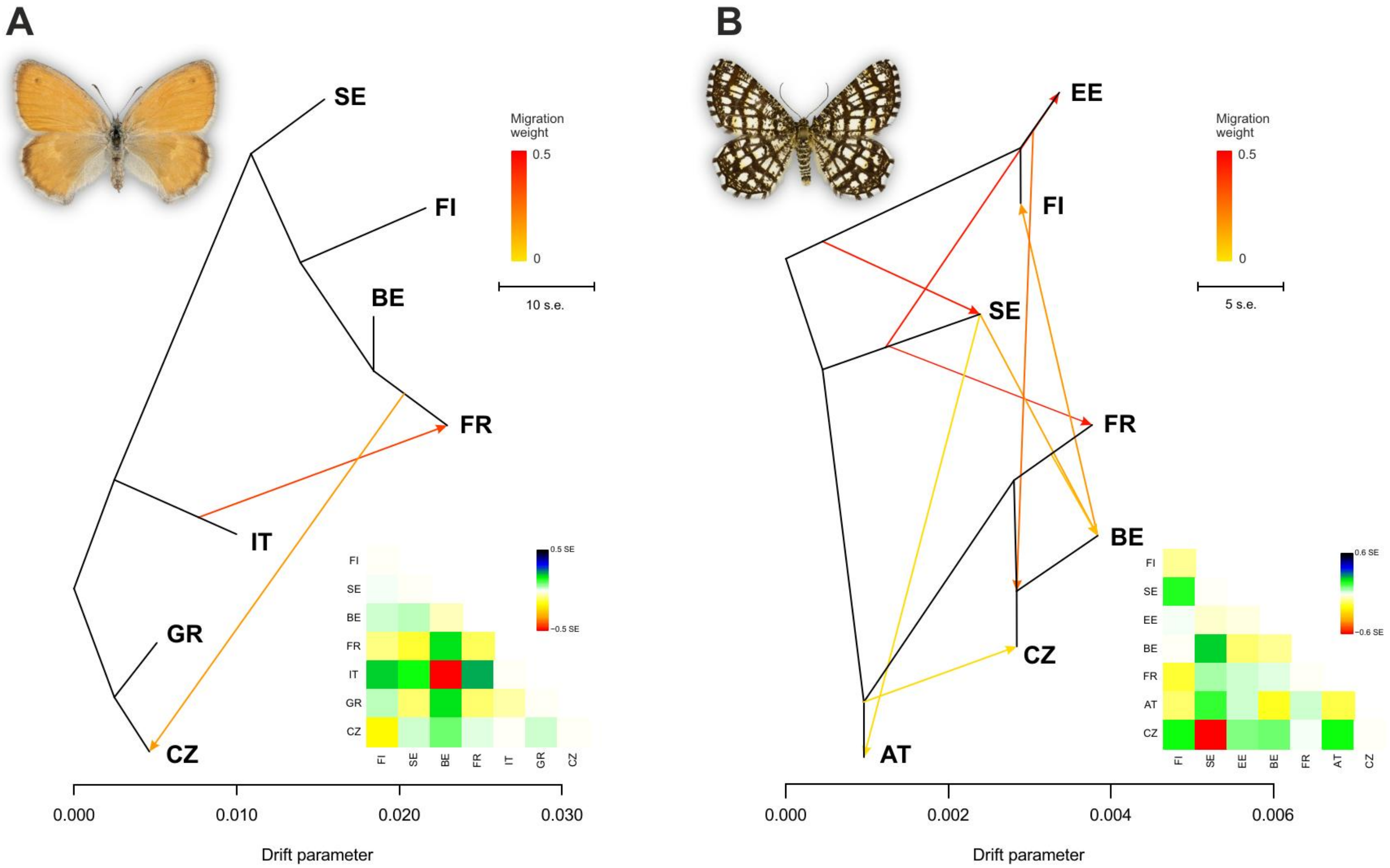


**Figure S3.** Country-level analysis of population splits and mixture in *Co. pamphilus* (A) and *Ch. clathrata* (B). The results are derived by using the TreeMix software [2] from an unlinked SNP (uSNP) dataset. For both species, a country-level tree is presented, with arrows indicating inferred historical migration events (arrow colour indicates migration weight). Scaled residuals of a maximum likelihood tree are illustrated for pairs of countries in the coloured matrices at the bottom right corners. These residuals are used for inferring gene flow events, with positive residuals indicating that populations are more closely related to each other than in the maximum likelihood tree, which could be a consequence of gene flow. Note that the scale for the drift parameter is much larger in A than in B, indicating that genetic differentiation among countries is much stronger in *Co. pamphilus* than in *Ch. clathrata*.

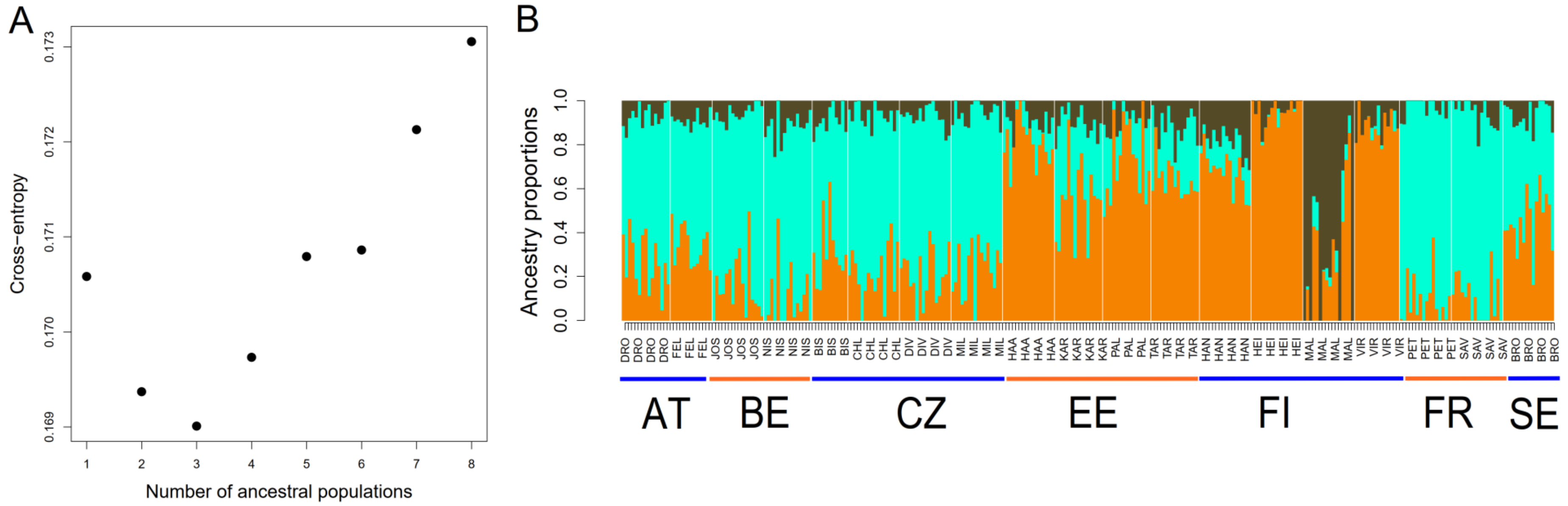


**Figure S4.** Sensitivity of the population admixture analysis in for inclusion of *Ch. clathrata* populations with low sample sizes. The cross-entropy values (A) and estimated ancestry proportions of individuals (B) are shown here for K=3, and when populations with a sample size less than ten individuals were removed from the analysis. See Figure 2 for other details.

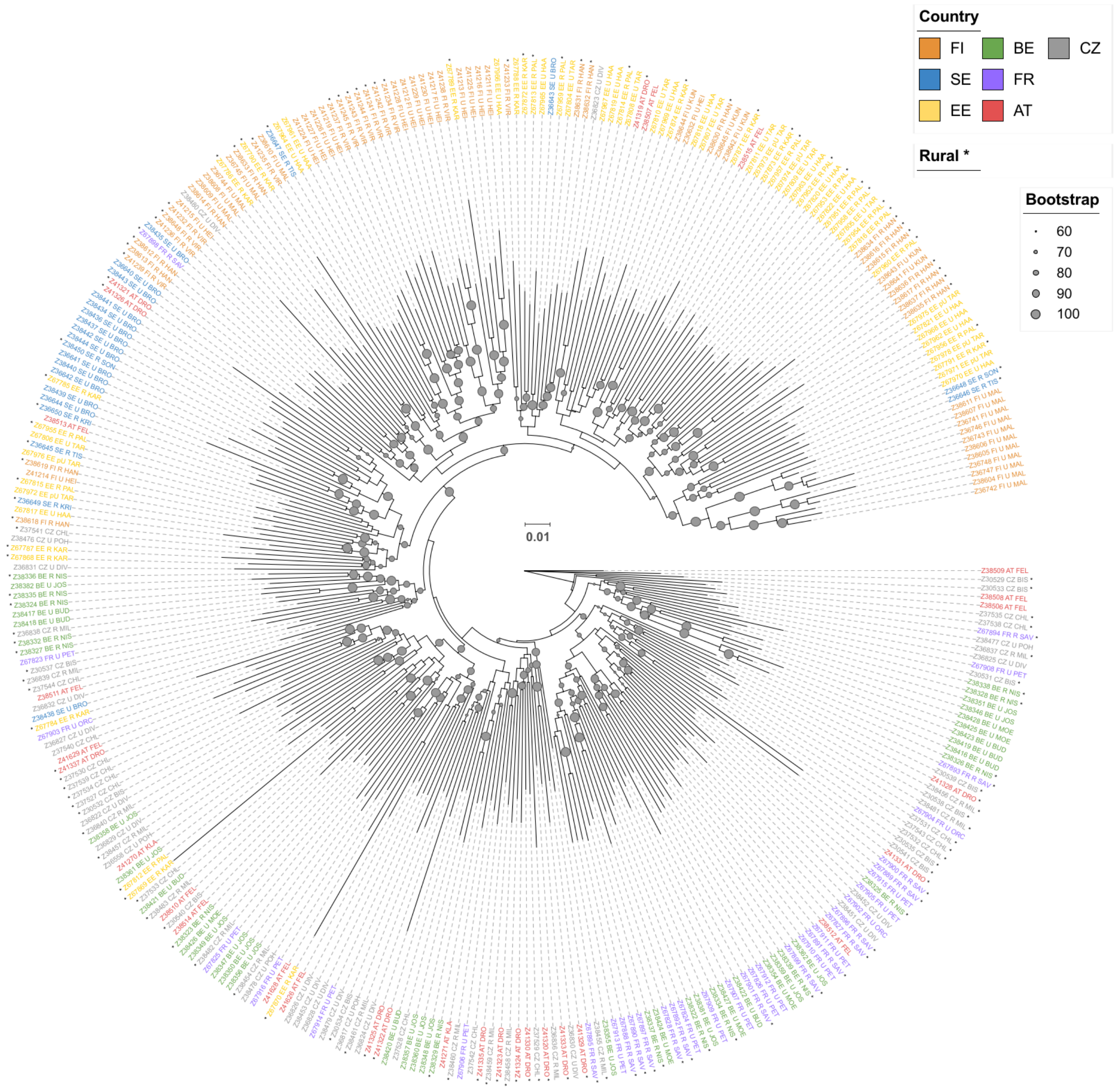


**Figure S5.** Maximum likelihood tree inferred from an IQ-TREE analysis based on the SNP dataset showing phylogenetic relationships among the studied *Ch. clathrata* individuals.

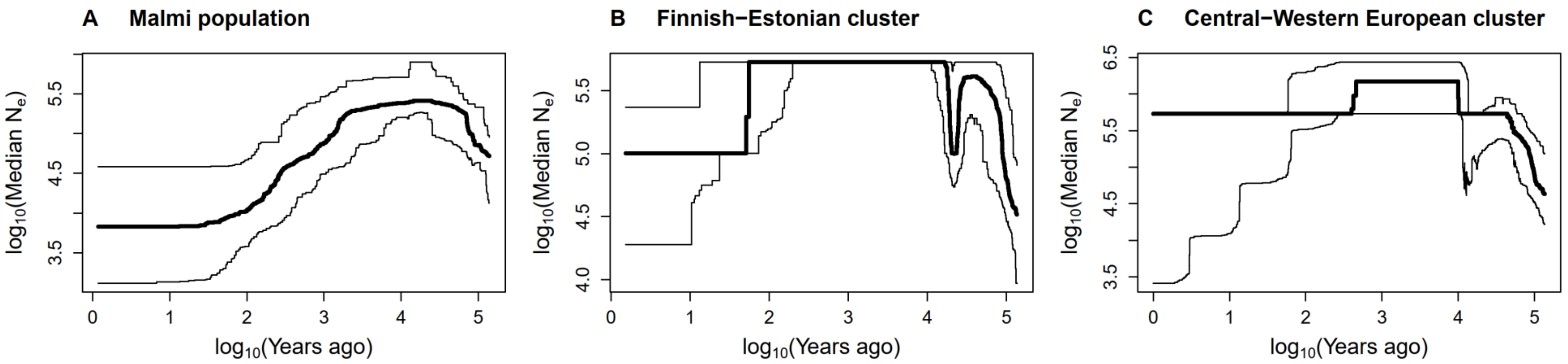


**Figure S6.** Estimated historical effective population size ($N_e$; thick line) and its 95% confidence intervals (thin lines) in the three genetic clusters of *Ch. clathrata* that were suggested by the population admixture analysis (see Fig. 2): (A) the Finnish urban Malmi population, (B) the remaining Finnish and Estonian populations, and (C) all the rest of the (Central and Western European) populations. Note that both axes of the panels are in logarithmic scale. The estimates of effective population sizes were derived with the stairwayplot2 software [3,4]. The numbers of sequences were 32, 224, and 394 for A, B, and C, respectively, and the sequence length being 139579 for each of them. We derived the cluster-specific site frequency spectra from the respective subsets of the .geno file (imputed to remove missing values) by using the R function 'gl.sfs' (package 'dartR.popgen' [5,6]) and set generation length to 0.5 years because almost all of the studied populations are bivoltine. Otherwise, we used the default settings of stairwayplot2 that were included in the template blueprint file for the Java program.

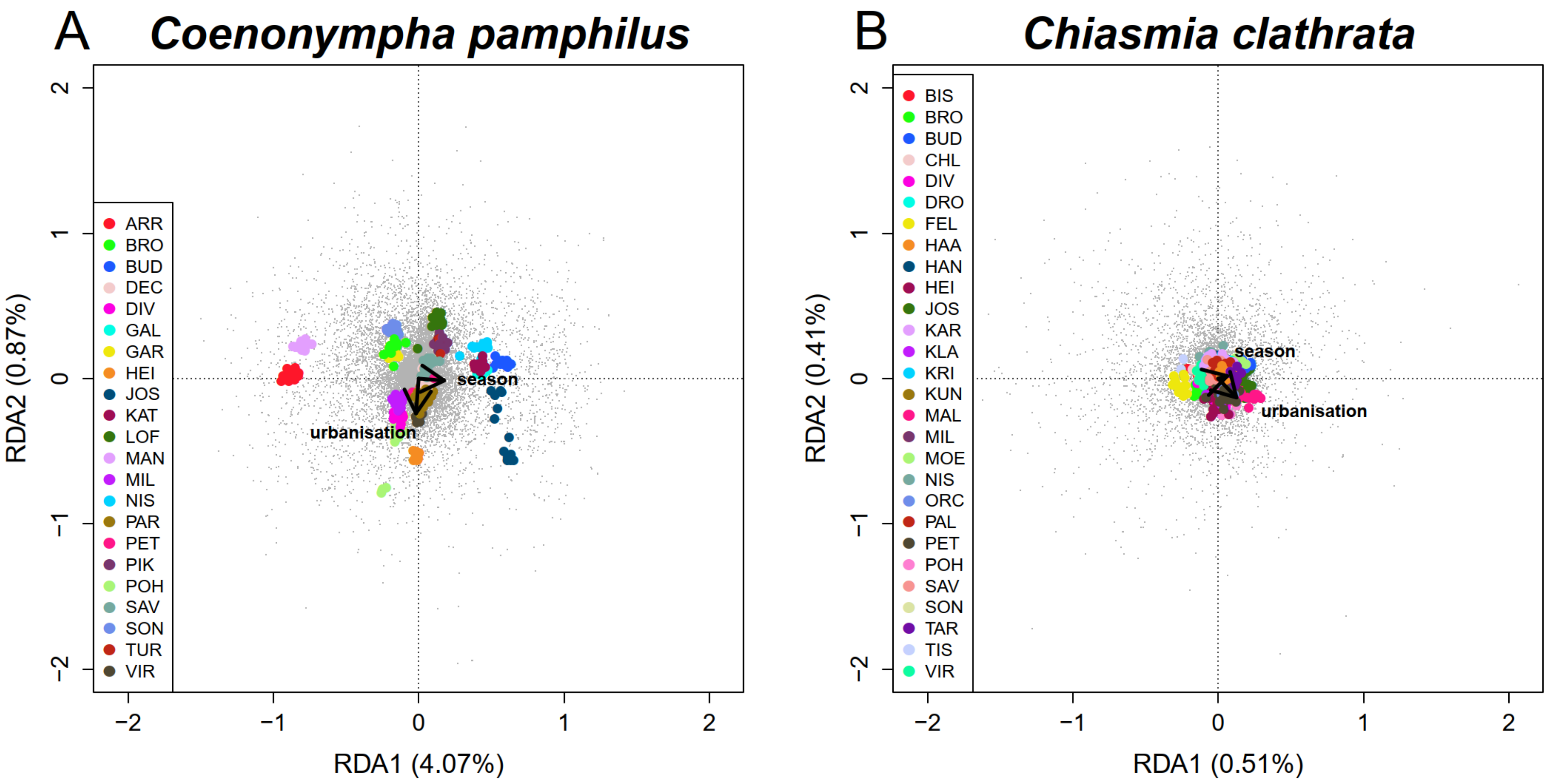


**Figure S7.** Visualization of results of a redundancy analysis (RDA) in *Co. pamphilus* (A) and *Ch. clathrata* (B). Scaling is such that the distances among objects depend on their similarity. The small grey points depict the locus (SNP) scores and the large, coloured points the individual scores (colour indicates the population of origin; see Table S1 for an abbreviation key, and Fig. 1 for geographic locations). Individuals that are close to each other have more similar genotypes than individuals that are further apart, and SNPs that are close to each other are present in more individuals than SNPs that are further apart. The arrows illustrate directions of increasing urbanisation and season length (arrow length is multiplied by three to improve readability). Proportions of variance explained by the RDA axes are given in the axis labels.

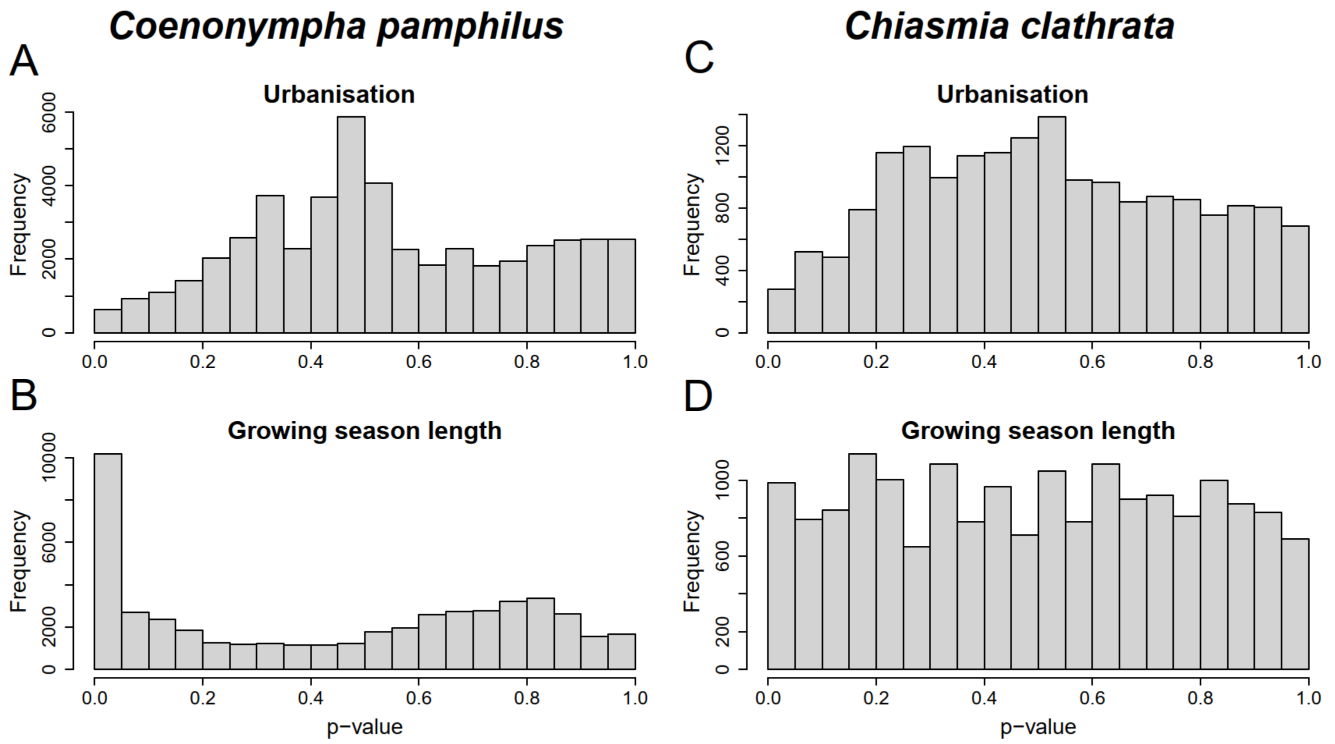


**Figure S8.** Histograms illustrating distributions of *p*-values of SNP-specific tests of association with urbanisation (A, C) and growing season length (B, D) in *Co. pamphilus* (A, B) and *Ch. clathrata* (C, D). Histograms are derived from the models used in inferences (12 latent factors in the model for *Co. pamphilus*; four latent factors in the model for *Ch. clathrata*). The model including 12 latent factors in *Co. pamphilus* resulted in a genomic inflation factor for growing season length that was closest to unity (1.004) and a relatively low genomic inflation factor for urbanisation (2.405; always > 2.1). We investigated models with 7–15 latent factors included for *Co. pamphilus*, the distribution of *p*-values for urbanisation being non-ideal in all investigated models, while that for growing season length being consistently ideal in this species. Differences among the models fitted with different numbers of latent factors were minimal, and the number of latent factors included did not affect the inferences. In *Ch. clathrata*, we compared models including 2–5 latent factors, and the model including four latent factors resulted in genomic inflation factors that were closest to unity (1.626 for urbanisation; 1.371 for growing season length). The distribution of *p*-values for urbanisation was non-ideal in all models also in *Ch. clathrata*, while that of growing season length was consistently uniform. Similarly as for *Co. pamphilus*, the number of latent factors included in the model had a minimal effect on diagnostics and results; inferences concerning *Ch. clathrata* remaining unaffected by the number of latent factors included in the model.

**Table S1.** Sampling sites (considered as populations) for both *Co. pamphilus* and *Ch. clathrata*, and basic site-specific information. Abbreviations for countries are: Austria (AT), Belgium (BE), Czech Republic (CZ), Estonia (EE), Finland (FI), France (FR), Greece (GR), Italy (IT), and Sweden (SE).

| Population (site) | Abbr.[a] | City / munici-pality | Coun-try | Latitude (WGS84) | Longitude (WGS84) | Degree of urbanisa-tion (pro-portion)[b] | Mean growing season length (days)[c] | N[d] | |
|---|---|---|---|---|---|---|---|---|---|
| | | | | | | | | *Co. pamphilus* | *Ch. clathrata* |
| Drösing | DRO | Drösing | AT | 48.519 | 16.908 | 0.018 | 252 | | 15 |
| Feldkirch | FEL | Feldkirch | AT | 47.261 | 9.542 | 0.064 | 224 | | 13 |
| Klagenfurt | KLA | Klagenfurt | AT | 46.649 | 14.333 | 0.191 | 238 | | 2 |
| Buda | BUD | Brussels | BE | 50.907 | 4.409 | 0.420 | 315 | 16 | 8 |
| Josaphat | JOS | Brussels | BE | 50.864 | 4.397 | 0.687 | 313 | 16 | 16 |
| Moeraske | MOE | Brussels | BE | 50.882 | 4.396 | 0.525 | 314 | | 6 |
| Nismes | NIS | Nismes | BE | 50.075 | 4.531 | 0.072 | 282 | 16 | 15 |
| Blšany u Loun | BIS | Bisanský chlum | CZ | 50.347 | 13.84 | 0.215 | 232 | | 11 |
| Chlum | CHL | Chlum | CZ | 50.868 | 14.563 | 0.020 | 216 | | 16 |
| Dívčí Hrady | DIV | Prague | CZ | 50.051 | 14.402 | 0.521 | 241 | 13 | 16 |
| Milovice | MIL | Milovice | CZ | 50.252 | 14.887 | 0.042 | 240 | 16 | 16 |
| Pohořelec | POH | Prague | CZ | 50.087 | 14.392 | 0.676 | 242 | 16 | 5 |
| Haabersti | HAA | Tallinn | EE | 59.424 | 24.638 | 0.243 | 182 | | 16 |
| Karilatsi | KAR | Karilatsi | EE | 58.124 | 26.915 | 0.002 | 185 | | 15 |
| Paldiski | PAL | Paldiski | EE | 59.359 | 24.051 | 0.081 | 182 | | 15 |
| Tartu | TAR | Tartu | EE | 58.362 | 26.681 | 0.313 | 187 | | 15 |
| Häntälä | HAN | Somero | FI | 60.576 | 23.351 | 0.003 | 172 | | 16 |
| Heinäpää | HEI | Oulu | FI | 64.999 | 25.45 | 0.472 | 150 | 16 | 16 |

| | | | | | | | | | |
|---|---|---|---|---|---|---|---|---|---|
| Kuninkaantammi | KUN | Helsinki | FI | 60.26 | 24.9 | 0.158 | 177 | | 5 |
| Lofsdal | LOF | Parainen | FI | 60.239 | 22.254 | 0.001 | 187 | 16 | |
| Malmi | MAL | Helsinki | FI | 60.247 | 25.048 | 0.345 | 177 | | 16 |
| Pikisaari | PIK | Turku | FI | 60.428 | 22.202 | 0.211 | 185 | 16 | |
| Turku | TUR | Turku | FI | 60.428 | 22.245 | 0.296 | 184 | 4 | |
| Virpiniemi | VIR | Oulu | FI | 65.157 | 25.301 | 0.077 | 146 | 16 | 14 |
| Decines-Charpieu | DEC | Lyon | FR | 45.776 | 4.96 | 0.440 | 295 | 16 | |
| Plaine des Orchidées | ORC | Lyon | FR | 45.79 | 4.882 | 0.535 | 299 | | 3 |
| Parc de Parilly | PAR | Lyon | FR | 45.716 | 4.902 | 0.720 | 299 | 16 | |
| Petit Parilly | PET | Lyon | FR | 45.719 | 4.92 | 0.585 | 298 | 6 | 16 |
| Savigny | SAV | Savigny | FR | 45.829 | 4.585 | 0.094 | 296 | 18 | 16 |
| Galaktokomiki | GAL | Ioánnina | GR | 39.634 | 20.882 | 0.185 | 305 | 16 | |
| Katsikas | KAT | Ioánnina | GR | 39.601 | 20.9 | 0.049 | 302 | 11 | |
| Parco dell'Arrivore | ARR | Torino | IT | 45.105 | 7.711 | 0.587 | 267 | 17 | |
| Parco La Mandria | MAN | Torino | IT | 45.145 | 7.602 | 0.071 | 257 | 18 | |
| Bromma | BRO | Stockholm | SE | 59.363 | 17.928 | 0.306 | 192 | 16 | 16 |
| Gärdet | GAR | Stockholm | SE | 59.339 | 18.106 | 0.482 | 193 | 15 | |
| Kristineholm | KRI | Norrtälje | SE | 59.841 | 18.474 | 0.002 | 187 | | 2 |
| Sonö | SON | Norrtälje | SE | 59.899 | 18.608 | 0.000 | 186 | 16 | 2 |
| Tisslinge | TIS | Norrtälje | SE | 59.863 | 18.814 | 0.000 | 187 | | 3 |

[a] Abbreviation of population (site) name.

[b] Proportion of land covered by impervious structures within 2500 m radius

[c] Mean length of growing season, defined as the number of days when mean daily temperature continuously exceeds 5 °C. Beginning and end of the season are defined as the dates after which mean daily temperature remains above and below the 5 °C threshold for seven consecutive days, respectively.
[d] Number of individuals included in the analysis.

**Table S2.** Estimated pairwise $F_{ST}$ (below diagonal) and Nei's genetic distance (above diagonal) between *Co. pamphilus* populations based on SNP data. $F_{ST}$ values that are statistically significant at the risk level of 0.05 are indicated by an asterisk. Nei's distances were calculated using StAMPP [7].

| | | 1 | 2 | 3 | 4 | 5 | 6 | 7 | 8 | 9 | 10 | 11 | 12 | 13 | 14 | 15 | 16 | 17 | 18 | 19 | 20 | 21 | 22 |
|---|---|---|---|---|---|---|---|---|---|---|---|---|---|---|---|---|---|---|---|---|---|---|---|
| 1 | HEI | 0 | 0.010 | 0.016 | 0.020 | 0.024 | 0.035 | 0.036 | 0.033 | 0.027 | 0.030 | 0.032 | 0.036 | 0.030 | 0.032 | 0.032 | 0.047 | 0.045 | 0.050 | 0.051 | 0.041 | 0.042 | 0.041 |
| 2 | VIR | 0.246* | 0 | 0.014 | 0.018 | 0.020 | 0.032 | 0.033 | 0.031 | 0.026 | 0.028 | 0.030 | 0.034 | 0.028 | 0.031 | 0.030 | 0.046 | 0.043 | 0.048 | 0.048 | 0.040 | 0.042 | 0.039 |
| 3 | PIK | 0.323* | 0.241* | 0 | 0.007 | 0.012 | 0.019 | 0.020 | 0.018 | 0.020 | 0.022 | 0.025 | 0.029 | 0.023 | 0.025 | 0.024 | 0.040 | 0.036 | 0.040 | 0.042 | 0.032 | 0.034 | 0.032 |
| 4 | LOF | 0.368* | 0.275* | 0.073* | 0 | 0.016 | 0.017 | 0.017 | 0.016 | 0.021 | 0.024 | 0.027 | 0.029 | 0.024 | 0.025 | 0.025 | 0.040 | 0.036 | 0.039 | 0.040 | 0.031 | 0.032 | 0.031 |
| 5 | TUR | 0.436* | 0.314* | 0.122* | 0.133* | 0 | 0.025 | 0.026 | 0.025 | 0.026 | 0.029 | 0.032 | 0.034 | 0.029 | 0.031 | 0.030 | 0.045 | 0.042 | 0.047 | 0.048 | 0.038 | 0.040 | 0.038 |
| 6 | GAR | 0.445* | 0.389* | 0.264* | 0.177* | 0.286* | 0 | 0.009 | 0.007 | 0.022 | 0.024 | 0.026 | 0.028 | 0.023 | 0.026 | 0.025 | 0.038 | 0.035 | 0.035 | 0.037 | 0.029 | 0.031 | 0.030 |
| 7 | SON | 0.478* | 0.425* | 0.264* | 0.185* | 0.321* | 0.099* | 0 | 0.007 | 0.023 | 0.027 | 0.027 | 0.029 | 0.024 | 0.027 | 0.026 | 0.039 | 0.035 | 0.036 | 0.037 | 0.029 | 0.031 | 0.029 |
| 8 | BRO | 0.459* | 0.382* | 0.277* | 0.195* | 0.288* | 0.081* | 0.114* | 0 | 0.021 | 0.024 | 0.026 | 0.028 | 0.023 | 0.025 | 0.024 | 0.039 | 0.035 | 0.035 | 0.036 | 0.028 | 0.030 | 0.029 |
| 9 | NIS | 0.393* | 0.361* | 0.264* | 0.271* | 0.268* | 0.296* | 0.295* | 0.272* | 0 | 0.009 | 0.012 | 0.017 | 0.012 | 0.015 | 0.013 | 0.036 | 0.033 | 0.038 | 0.039 | 0.029 | 0.030 | 0.029 |
| 10 | BUD | 0.449* | 0.400* | 0.331* | 0.311* | 0.317* | 0.338* | 0.350* | 0.325* | 0.043* | 0 | 0.014 | 0.020 | 0.015 | 0.017 | 0.016 | 0.039 | 0.036 | 0.041 | 0.042 | 0.032 | 0.034 | 0.031 |
| 11 | JOS | 0.497* | 0.478* | 0.401* | 0.385* | 0.407* | 0.406* | 0.396* | 0.394* | 0.159* | 0.185* | 0 | 0.023 | 0.019 | 0.020 | 0.019 | 0.043 | 0.039 | 0.045 | 0.045 | 0.035 | 0.036 | 0.034 |
| 12 | PET | 0.466* | 0.410* | 0.351* | 0.325* | 0.276* | 0.343* | 0.357* | 0.279* | 0.147* | 0.190* | 0.276* | 0 | 0.012 | 0.013 | 0.012 | 0.033 | 0.029 | 0.037 | 0.038 | 0.030 | 0.032 | 0.030 |
| 13 | SAV | 0.404* | 0.368* | 0.311* | 0.292* | 0.270* | 0.267* | 0.302* | 0.260* | 0.120* | 0.147* | 0.224* | 0.006 | 0 | 0.010 | 0.008 | 0.028 | 0.025 | 0.033 | 0.033 | 0.025 | 0.026 | 0.025 |
| 14 | PAR | 0.420* | 0.397* | 0.353* | 0.340* | 0.314* | 0.329* | 0.356* | 0.308* | 0.148* | 0.191* | 0.262* | -0.008 | 0.030* | 0 | 0.009 | 0.029 | 0.027 | 0.034 | 0.035 | 0.027 | 0.028 | 0.027 |
| 15 | DEC | 0.437* | 0.392* | 0.334* | 0.326* | 0.321* | 0.302* | 0.323* | 0.256* | 0.152* | 0.196* | 0.257* | 0.004 | 0.008 | 0.050* | 0 | 0.029 | 0.026 | 0.034 | 0.034 | 0.026 | 0.027 | 0.026 |
| 16 | ARR | 0.522* | 0.464* | 0.398* | 0.367* | 0.394* | 0.357* | 0.393* | 0.324* | 0.321* | 0.359* | 0.440* | 0.231* | 0.210* | 0.246* | 0.215* | 0 | 0.015 | 0.029 | 0.030 | 0.027 | 0.029 | 0.028 |
| 17 | MAN | 0.473* | 0.434* | 0.377* | 0.337* | 0.344* | 0.331* | 0.359* | 0.302* | 0.320* | 0.349* | 0.408* | 0.206* | 0.207* | 0.239* | 0.224* | 0.045* | 0 | 0.025 | 0.026 | 0.024 | 0.024 | 0.024 |
| 18 | GAL | 0.540* | 0.491* | 0.434* | 0.392* | 0.393* | 0.360* | 0.384* | 0.334* | 0.389* | 0.411* | 0.456* | 0.295* | 0.278* | 0.295* | 0.280* | 0.182* | 0.143* | 0 | 0.012 | 0.015 | 0.016 | 0.015 |
| 19 | KAT | 0.546* | 0.495* | 0.446* | 0.402* | 0.403* | 0.352* | 0.385* | 0.325* | 0.401* | 0.427* | 0.460* | 0.302* | 0.290* | 0.304* | 0.279* | 0.175* | 0.153* | 0.021 | 0 | 0.016 | 0.016 | 0.017 |
| 20 | MIL | 0.477* | 0.426* | 0.384* | 0.344* | 0.348* | 0.301* | 0.330* | 0.312* | 0.332* | 0.348* | 0.380* | 0.269* | 0.233* | 0.254* | 0.244* | 0.196* | 0.162* | 0.044* | 0.087* | 0 | 0.011 | 0.011 |
| 21 | POH | 0.520* | 0.470* | 0.417* | 0.379* | 0.406* | 0.357* | 0.366* | 0.342* | 0.369* | 0.392* | 0.419* | 0.328* | 0.287* | 0.315* | 0.292* | 0.235* | 0.223* | 0.108* | 0.120* | 0.061* | 0 | 0.012 |
| 22 | DIV | 0.493* | 0.453* | 0.372* | 0.333* | 0.357* | 0.298* | 0.319* | 0.323* | 0.337* | 0.347* | 0.395* | 0.296* | 0.251* | 0.285* | 0.268* | 0.228* | 0.192* | 0.104* | 0.124* | -0.001 | 0.059* | 0 |

**Table S3.** Estimated pairwise $F_{ST}$ (below diagonal) and Nei's genetic distance (above diagonal) between *Ch. clathrata* populations based on SNP data. $F_{ST}$ values that are statistically significant at the risk level of 0.05 are indicated by an asterisk. Nei's distances were calculated using StAMPP [7].

| | | 1 | 2 | 3 | 4 | 5 | 6 | 7 | 8 | 9 | 10 | 11 | 12 | 13 | 14 | 15 | 16 | 17 | 18 | 19 | 20 | 21 | 22 | 23 | 24 | 25 | 26 | 27 | 28 |
|---|---|---|---|---|---|---|---|---|---|---|---|---|---|---|---|---|---|---|---|---|---|---|---|---|---|---|---|---|---|
| 1 | MAL | 0 | 0.009 | 0.017 | 0.012 | 0.013 | 0.011 | 0.023 | 0.023 | 0.021 | 0.015 | 0.011 | 0.010 | 0.011 | 0.014 | 0.016 | 0.019 | 0.021 | 0.014 | 0.014 | 0.024 | 0.018 | 0.014 | 0.027 | 0.015 | 0.017 | 0.014 | 0.012 | 0.012 |
| 2 | HAN | 0.045* | 0 | 0.013 | 0.009 | 0.010 | 0.008 | 0.021 | 0.020 | 0.020 | 0.011 | 0.009 | 0.007 | 0.008 | 0.012 | 0.013 | 0.017 | 0.016 | 0.011 | 0.012 | 0.022 | 0.015 | 0.010 | 0.026 | 0.013 | 0.014 | 0.010 | 0.010 | 0.010 |
| 3 | KUN | 0.182* | 0.089* | 0 | 0.016 | 0.017 | 0.014 | 0.027 | 0.025 | 0.024 | 0.017 | 0.014 | 0.014 | 0.015 | 0.018 | 0.019 | 0.022 | 0.022 | 0.017 | 0.017 | 0.027 | 0.021 | 0.017 | 0.029 | 0.017 | 0.019 | 0.017 | 0.015 | 0.016 |
| 4 | HEI | 0.125* | 0.112* | 0.214* | 0 | 0.010 | 0.010 | 0.021 | 0.022 | 0.021 | 0.013 | 0.010 | 0.010 | 0.010 | 0.013 | 0.015 | 0.017 | 0.018 | 0.012 | 0.013 | 0.024 | 0.016 | 0.012 | 0.025 | 0.014 | 0.016 | 0.012 | 0.012 | 0.011 |
| 5 | VIR | 0.058* | 0.030 | 0.132* | 0.040* | 0 | 0.009 | 0.021 | 0.021 | 0.021 | 0.013 | 0.009 | 0.011 | 0.011 | 0.012 | 0.014 | 0.018 | 0.019 | 0.013 | 0.013 | 0.026 | 0.016 | 0.012 | 0.027 | 0.014 | 0.015 | 0.012 | 0.010 | 0.011 |
| 6 | BRO | 0.141* | 0.070* | 0.085* | 0.102* | 0.018 | 0 | 0.019 | 0.019 | 0.018 | 0.012 | 0.008 | 0.007 | 0.008 | 0.009 | 0.012 | 0.015 | 0.016 | 0.009 | 0.009 | 0.021 | 0.013 | 0.011 | 0.023 | 0.011 | 0.012 | 0.009 | 0.007 | 0.010 |
| 7 | TIS | 0.038 | 0.007 | 0.073 | -0.034 | -0.077 | -0.243 | 0 | 0.028 | 0.029 | 0.023 | 0.020 | 0.020 | 0.021 | 0.022 | 0.023 | 0.025 | 0.026 | 0.021 | 0.023 | 0.032 | 0.024 | 0.022 | 0.034 | 0.024 | 0.026 | 0.021 | 0.021 | 0.021 |
| 8 | KRI | -0.003 | -0.105 | 0.105 | 0.080 | -0.108 | -0.056 | -0.536 | 0 | 0.030 | 0.022 | 0.019 | 0.019 | 0.020 | 0.022 | 0.024 | 0.026 | 0.026 | 0.022 | 0.022 | 0.032 | 0.027 | 0.021 | 0.033 | 0.023 | 0.023 | 0.021 | 0.020 | 0.020 |
| 9 | SON | 0.225* | 0.157* | 0.172 | 0.150* | 0.035 | -0.006 | 0.042 | 0.070 | 0 | 0.022 | 0.019 | 0.019 | 0.019 | 0.021 | 0.022 | 0.026 | 0.026 | 0.020 | 0.021 | 0.029 | 0.023 | 0.022 | 0.030 | 0.023 | 0.022 | 0.020 | 0.018 | 0.019 |
| 10 | KAR | 0.059* | 0.007 | 0.058 | -0.018 | -0.108 | -0.069 | 0.101 | 0.020 | 0.208 | 0 | 0.011 | 0.012 | 0.013 | 0.014 | 0.016 | 0.019 | 0.020 | 0.014 | 0.014 | 0.026 | 0.017 | 0.014 | 0.028 | 0.017 | 0.018 | 0.014 | 0.012 | 0.013 |
| 11 | TAR | 0.116* | 0.054* | 0.068* | 0.044* | -0.049 | 0.005 | -0.154 | -0.036 | -0.044 | -0.184 | 0 | 0.008 | 0.008 | 0.011 | 0.013 | 0.016 | 0.017 | 0.011 | 0.011 | 0.023 | 0.015 | 0.011 | 0.024 | 0.013 | 0.014 | 0.010 | 0.008 | 0.009 |
| 12 | PAL | 0.090* | 0.060* | 0.072* | 0.072* | 0.026 | 0.025 | -0.083 | 0.017 | 0.092 | -0.160 | -0.019 | 0 | 0.007 | 0.011 | 0.012 | 0.017 | 0.017 | 0.011 | 0.010 | 0.022 | 0.014 | 0.010 | 0.024 | 0.012 | 0.013 | 0.010 | 0.008 | 0.009 |
| 13 | HAA | 0.106* | 0.085* | 0.132* | 0.125* | 0.047* | 0.050* | -0.041 | 0.034 | 0.165* | 0.033 | 0.023 | 0.047* | 0 | 0.011 | 0.013 | 0.017 | 0.018 | 0.012 | 0.012 | 0.022 | 0.015 | 0.011 | 0.025 | 0.013 | 0.014 | 0.011 | 0.009 | 0.010 |
| 14 | NIS | 0.181* | 0.143* | 0.098* | 0.172* | 0.098* | 0.066* | 0.063 | 0.152 | 0.204* | -0.002 | 0.059* | 0.077* | 0.122* | 0 | 0.012 | 0.015 | 0.014 | 0.009 | 0.009 | 0.020 | 0.015 | 0.010 | 0.025 | 0.013 | 0.014 | 0.010 | 0.009 | 0.010 |
| 15 | JOS | 0.145* | 0.133* | 0.148* | 0.133* | 0.078* | 0.075* | 0.026 | 0.209* | 0.280* | -0.040 | 0.048* | 0.028 | 0.114* | 0.033 | 0 | 0.015 | 0.014 | 0.010 | 0.010 | 0.021 | 0.015 | 0.012 | 0.025 | 0.014 | 0.015 | 0.011 | 0.011 | 0.012 |
| 16 | MOE | 0.087* | 0.132* | 0.100 | 0.124* | 0.030 | 0.119* | 0.023 | 0.108 | 0.251 | 0.035 | 0.044 | 0.051 | 0.091* | 0.005 | 0.050 | 0 | 0.019 | 0.014 | 0.014 | 0.024 | 0.020 | 0.016 | 0.029 | 0.018 | 0.020 | 0.016 | 0.015 | 0.015 |
| 17 | BUD | 0.186* | 0.172* | 0.193* | 0.224* | 0.153* | 0.124* | 0.097 | 0.135 | 0.166 | 0.031 | 0.125* | 0.144* | 0.204* | 0.070* | 0.108* | 0.097* | 0 | 0.014 | 0.014 | 0.027 | 0.019 | 0.017 | 0.028 | 0.018 | 0.019 | 0.015 | 0.014 | 0.015 |
| 18 | PET | 0.136* | 0.125* | 0.082* | 0.124* | 0.070* | 0.031* | -0.103 | 0.040 | 0.013 | -0.086 | 0.005 | 0.060* | 0.086* | 0.003 | 0.046* | 0.048 | 0.022 | 0 | 0.007 | 0.017 | 0.014 | 0.009 | 0.023 | 0.012 | 0.014 | 0.010 | 0.008 | 0.009 |
| 19 | SAV | 0.180* | 0.109* | 0.118* | 0.134* | 0.072* | 0.045* | -0.061 | 0.017 | 0.057 | -0.041 | 0.042* | 0.083* | 0.122* | 0.004 | 0.047* | 0.019 | 0.098* | -0.016 | 0 | 0.017 | 0.014 | 0.008 | 0.023 | 0.012 | 0.013 | 0.010 | 0.008 | 0.008 |
| 20 | ORC | 0.109* | 0.099 | 0.108 | 0.210* | 0.017 | -0.030 | 0.045 | 0.009 | 0.234 | -0.005 | 0.012 | 0.007 | 0.140* | -0.011 | 0.117* | 0.093 | 0.017 | -0.019 | 0.005 | 0 | 0.025 | 0.019 | 0.033 | 0.023 | 0.025 | 0.021 | 0.019 | 0.019 |
| 21 | FEL | 0.002 | -0.021 | 0.100* | 0.010 | -0.055 | -0.086 | -0.025 | -0.233 | 0.151* | 0.061* | -0.117 | -0.055 | 0.078* | 0.005 | 0.052* | 0.012 | 0.033 | -0.093 | -0.047 | 0.043 | 0 | 0.015 | 0.026 | 0.017 | 0.017 | 0.014 | 0.012 | 0.013 |
| 22 | DRO | 0.117* | 0.124* | 0.127* | 0.143* | 0.050* | 0.030 | 0.020 | 0.156* | 0.208* | -0.029 | 0.013 | 0.044* | 0.113* | 0.047* | 0.052* | 0.122* | 0.119* | -0.035 | 0.049* | 0.068 | 0.017 | 0 | 0.025 | 0.013 | 0.013 | 0.010 | 0.007 | 0.009 |
| 23 | KLA | 0.356* | 0.342* | 0.397* | 0.367* | 0.300* | 0.231* | 0.239 | 0.160 | 0.480 | 0.452* | 0.274* | 0.330* | 0.347* | 0.290* | 0.364* | 0.223* | 0.347* | 0.204* | 0.215* | 0.320 | 0.240* | 0.340* | 0 | 0.026 | 0.027 | 0.023 | 0.022 | 0.022 |
| 24 | BIS | 0.034 | -0.026 | -0.063 | 0.004 | -0.063 | -0.067 | -0.074 | -0.132 | 0.129 | -0.009 | -0.122 | -0.076 | 0.074* | -0.040 | -0.008 | 0.046 | 0.045 | -0.090 | -0.062 | -0.007 | -0.004 | 0.009 | 0.233* | 0 | 0.016 | 0.012 | 0.010 | 0.011 |
| 25 | POH | 0.152* | 0.135* | -0.009 | 0.054 | 0.010 | 0.015 | -0.131 | 0.068 | -0.091 | 0.071 | 0.021 | 0.078* | 0.127* | 0.124* | 0.135* | 0.097* | 0.170* | -0.034 | 0.035 | 0.153 | -0.013 | 0.017 | 0.319* | -0.037 | 0 | 0.013 | 0.012 | 0.014 |
| 26 | DIV | 0.116* | 0.081* | 0.105* | 0.123* | 0.061* | 0.029 | -0.060 | 0.085 | 0.066 | -0.023 | 0.016 | 0.059* | 0.099* | 0.069* | 0.063* | 0.108* | 0.050 | 0.010 | 0.052* | 0.080 | -0.021 | 0.003 | 0.347* | -0.047 | 0.029 | 0 | 0.007 | 0.009 |
| 27 | MIL | 0.102* | 0.084* | 0.062 | 0.099* | 0.038* | -0.003 | -0.166 | 0.047 | 0.054 | -0.111 | -0.021 | 0.034* | 0.079* | 0.046* | 0.054 | 0.050 | 0.068* | -0.015 | 0.017 | 0.029 | -0.087 | -0.025 | 0.278* | -0.099 | -0.016 | 0.005* | 0 | 0.007 |
| 28 | CHL | 0.146* | 0.072* | 0.086* | 0.124* | 0.026 | 0.033* | -0.037 | 0.077 | 0.134* | -0.024 | 0.028 | 0.048* | 0.112* | 0.032 | 0.071* | 0.073* | 0.103* | 0.024 | 0.040* | 0.062 | -0.035 | 0.013 | 0.266* | -0.075 | 0.009 | 0.026 | -0.030 | 0 |

**Table S4.** Details of DNA sequencing data for each *Co. pamphilus* individual. The sample identities (SampleID) are combinations of a unique code, species abbreviation ("pamp"), country abbreviation (see Table S1 for a key for this and the population abbreviations), and a habitat type indicator ("R" for rural; "U" for urban). Sequencing batch indicates the sequencing platform (1 = HiSeq X PE150; 2 = NovaSeq X Plus PE150). Clusters refer to the number of clusters in initial grouping of similar sequences within an individual. Average depth refers to the average read depth. Hetero estimate refers to the estimated per-site heterozygosity within individual samples. Error estimate is the estimated per-site sequencing error rate, inferred from base call mismatches among reads within a cluster. Reads consensus refers to the number of consensus loci (assembled clusters) successfully built after clustering and filtering. Heterozygosity is the average heterozygosity across loci. Loci in assembly is the total number of RAD loci retained in the final assembly after applying all filters. Missing data is the proportion of missing data.

| No. | SampleID | Population | Sequencing plate | Sequencing batch | Gb sequenced | Total reads | Reads passed filter | Clusters | Average depth | Hetero estimate | Error estimate | Reads consensus | Heterozygosity | Loci in assembly | Missing data |
|---|---|---|---|---|---|---|---|---|---|---|---|---|---|---|---|
| 1 | Z30026_pamp_FI_U | HEI | plate 4 | 1 | 0.63 | 2102479 | 2102471 | 76477 | 15.20 | 0.0033 | 0.0006 | 27449 | 0.0016 | 1191 | 0.45 |
| 2 | Z30027_pamp_FI_U | HEI | plate 4 | 1 | 0.43 | 1439070 | 1439063 | 50421 | 19.29 | 0.0032 | 0.0006 | 25121 | 0.0019 | 1387 | 0.37 |
| 3 | Z30028_pamp_FI_U | HEI | plate 4 | 1 | 0.58 | 1936688 | 1936681 | 60841 | 21.44 | 0.0034 | 0.0006 | 26146 | 0.0018 | 1395 | 0.36 |
| 4 | Z30029_pamp_FI_U | HEI | plate 4 | 1 | 0.72 | 2397756 | 2397745 | 65209 | 24.88 | 0.0036 | 0.0005 | 27595 | 0.0019 | 1472 | 0.32 |
| 5 | Z30030_pamp_FI_U | HEI | plate 4 | 1 | 0.49 | 1633149 | 1633143 | 52136 | 21.22 | 0.0034 | 0.0006 | 24436 | 0.0019 | 1408 | 0.35 |
| 6 | Z30031_pamp_FI_U | HEI | plate 4 | 1 | 0.80 | 2668846 | 2668833 | 78187 | 20.68 | 0.0035 | 0.0006 | 28360 | 0.0018 | 1300 | 0.39 |
| 7 | Z30032_pamp_FI_U | HEI | plate 4 | 1 | 0.69 | 2291173 | 2291164 | 68301 | 21.83 | 0.0034 | 0.0006 | 27442 | 0.0017 | 1366 | 0.38 |
| 8 | Z30033_pamp_FI_U | HEI | plate 4 | 1 | 0.63 | 2103520 | 2103510 | 62677 | 21.67 | 0.0033 | 0.0006 | 26377 | 0.0019 | 1371 | 0.37 |
| 9 | Z30034_pamp_FI_U | HEI | plate 4 | 1 | 0.53 | 1763669 | 1763662 | 57790 | 19.03 | 0.0039 | 0.0006 | 24332 | 0.0022 | 1466 | 0.33 |
| 10 | Z30035_pamp_FI_U | HEI | plate 4 | 1 | 0.59 | 1964021 | 1964006 | 70734 | 17.57 | 0.0033 | 0.0006 | 24872 | 0.0018 | 1366 | 0.37 |
| 11 | Z30042_pamp_FI_U | HEI | plate 4 | 1 | 0.41 | 1381649 | 1381639 | 55355 | 16.13 | 0.0032 | 0.0006 | 26819 | 0.0015 | 1419 | 0.35 |
| 12 | Z30043_pamp_FI_U | HEI | plate 4 | 1 | 0.46 | 1533152 | 1533146 | 45094 | 21.21 | 0.0030 | 0.0006 | 23260 | 0.0018 | 1401 | 0.36 |
| 13 | Z30044_pamp_FI_U | HEI | plate 4 | 1 | 0.57 | 1884798 | 1884786 | 57594 | 21.34 | 0.0033 | 0.0006 | 25468 | 0.0017 | 1340 | 0.38 |
| 14 | Z30045_pamp_FI_U | HEI | plate 4 | 1 | 0.55 | 1844649 | 1844639 | 57848 | 20.22 | 0.0034 | 0.0006 | 26427 | 0.0018 | 1416 | 0.35 |
| 15 | Z30046_pamp_FI_U | HEI | plate 4 | 1 | 0.64 | 2130949 | 2130939 | 61555 | 21.84 | 0.0036 | 0.0005 | 26605 | 0.0020 | 1398 | 0.35 |
| 16 | Z30047_pamp_FI_U | HEI | plate 4 | 1 | 0.49 | 1633922 | 1633913 | 54289 | 19.45 | 0.0035 | 0.0005 | 25708 | 0.0019 | 1411 | 0.35 |
| 17 | Z30083_pamp_FI_R | VIR | plate 5 | 2 | 0.33 | 1085779 | 1085626 | 42948 | 16.78 | 0.0039 | 0.0006 | 20585 | 0.0020 | 1403 | 0.34 |
| 18 | Z30084_pamp_FI_R | VIR | plate 5 | 2 | 0.49 | 1623860 | 1623606 | 66065 | 14.48 | 0.0038 | 0.0005 | 23139 | 0.0018 | 1297 | 0.39 |
| 19 | Z30085_pamp_FI_R | VIR | plate 5 | 2 | 0.72 | 2385365 | 2385032 | 69013 | 18.86 | 0.0040 | 0.0005 | 27581 | 0.0018 | 1439 | 0.33 |
| 20 | Z30086_pamp_FI_R | VIR | plate 5 | 2 | 0.77 | 2574949 | 2574584 | 75340 | 19.62 | 0.0064 | 0.0005 | 28651 | 0.0017 | 1345 | 0.38 |
| 21 | Z30087_pamp_FI_R | VIR | plate 5 | 2 | 0.50 | 1654474 | 1654275 | 56029 | 19.05 | 0.0041 | 0.0006 | 24837 | 0.0020 | 1463 | 0.32 |

| | | | | | | | | | | | | | | | |
|---|---|---|---|---|---|---|---|---|---|---|---|---|---|---|---|
| 22 | Z30088_pamp_FI_R | VIR | plate 5 | 2 | 0.76 | 2525053 | 2524710 | 67141 | 23.72 | 0.0042 | 0.0005 | 27789 | 0.0019 | 1391 | 0.35 |
| 23 | Z30089_pamp_FI_R | VIR | plate 5 | 2 | 0.62 | 2055580 | 2055281 | 64015 | 19.50 | 0.0042 | 0.0005 | 26695 | 0.0019 | 1385 | 0.36 |
| 24 | Z30090_pamp_FI_R | VIR | plate 5 | 2 | 0.64 | 2144656 | 2144330 | 63511 | 21.86 | 0.0042 | 0.0005 | 26769 | 0.0019 | 1412 | 0.34 |
| 25 | Z30092_pamp_FI_R | VIR | plate 4 | 1 | 0.64 | 2124621 | 2124606 | 43279 | 27.67 | 0.0038 | 0.0006 | 21899 | 0.0020 | 1477 | 0.32 |
| 26 | Z36615_pamp_SE_U | GAR | plate 5 | 2 | 0.68 | 2262732 | 2262391 | 69725 | 15.74 | 0.0051 | 0.0006 | 28417 | 0.0023 | 1408 | 0.34 |
| 27 | Z36616_pamp_SE_U | GAR | plate 5 | 2 | 0.50 | 1651210 | 1650978 | 67263 | 14.40 | 0.0047 | 0.0006 | 27746 | 0.0023 | 1440 | 0.33 |
| 28 | Z36617_pamp_SE_U | GAR | plate 5 | 2 | 0.55 | 1843336 | 1843090 | 65133 | 16.67 | 0.0048 | 0.0006 | 27729 | 0.0024 | 1412 | 0.35 |
| 29 | Z36618_pamp_SE_U | GAR | plate 5 | 2 | 0.63 | 2112023 | 2111725 | 69709 | 16.50 | 0.0048 | 0.0006 | 28495 | 0.0023 | 1343 | 0.37 |
| 30 | Z36619_pamp_SE_U | GAR | plate 5 | 2 | 0.65 | 2174470 | 2174177 | 66524 | 19.52 | 0.0047 | 0.0005 | 27098 | 0.0023 | 1437 | 0.33 |
| 31 | Z36620_pamp_SE_U | GAR | plate 5 | 2 | 0.49 | 1634355 | 1634101 | 58465 | 16.24 | 0.0045 | 0.0005 | 25171 | 0.0024 | 1392 | 0.34 |
| 32 | Z36684_pamp_BE_U | JOS | plate 3 | 1 | 0.73 | 2420483 | 2420469 | 76811 | 19.77 | 0.0046 | 0.0006 | 33174 | 0.0022 | 1408 | 0.35 |
| 33 | Z36685_pamp_BE_U | JOS | plate 3 | 1 | 0.39 | 1311237 | 1311224 | 55450 | 14.26 | 0.0047 | 0.0006 | 25805 | 0.0025 | 1404 | 0.35 |
| 34 | Z36699_pamp_FI_U | PIK | plate 4 | 1 | 0.45 | 1512684 | 1512675 | 64210 | 14.85 | 0.0043 | 0.0006 | 25118 | 0.0023 | 1478 | 0.32 |
| 35 | Z36700_pamp_FI_U | PIK | plate 4 | 1 | 0.49 | 1627090 | 1627080 | 58968 | 18.79 | 0.0040 | 0.0006 | 23673 | 0.0023 | 1457 | 0.34 |
| 36 | Z36701_pamp_FI_U | PIK | plate 4 | 1 | 0.63 | 2099247 | 2099235 | 60365 | 19.84 | 0.0043 | 0.0006 | 24197 | 0.0024 | 1488 | 0.32 |
| 37 | Z36702_pamp_FI_U | PIK | plate 4 | 1 | 1.00 | 3318274 | 3318262 | 93867 | 17.90 | 0.0047 | 0.0006 | 28551 | 0.0022 | 1419 | 0.35 |
| 38 | Z36703_pamp_FI_U | PIK | plate 4 | 1 | 0.47 | 1568165 | 1568160 | 58585 | 19.39 | 0.0043 | 0.0006 | 23193 | 0.0025 | 1489 | 0.32 |
| 39 | Z36704_pamp_FI_U | PIK | plate 4 | 1 | 0.37 | 1247289 | 1247281 | 54841 | 15.58 | 0.0044 | 0.0007 | 22472 | 0.0024 | 1508 | 0.32 |
| 40 | Z36705_pamp_FI_U | PIK | plate 4 | 1 | 0.68 | 2253677 | 2253658 | 66606 | 19.55 | 0.0042 | 0.0006 | 27188 | 0.0023 | 1452 | 0.33 |
| 41 | Z36706_pamp_FI_U | PIK | plate 4 | 1 | 0.86 | 2872170 | 2872160 | 66640 | 29.52 | 0.0042 | 0.0006 | 27028 | 0.0022 | 1490 | 0.32 |
| 42 | Z36707_pamp_FI_U | PIK | plate 4 | 1 | 0.30 | 1015418 | 1015413 | 45967 | 14.49 | 0.0047 | 0.0007 | 22155 | 0.0022 | 1463 | 0.34 |
| 43 | Z36708_pamp_FI_U | PIK | plate 4 | 1 | 0.56 | 1873169 | 1873157 | 56467 | 21.16 | 0.0043 | 0.0006 | 23147 | 0.0024 | 1485 | 0.33 |
| 44 | Z36777_pamp_BE_U | BUD | plate 3 | 1 | 0.67 | 2222597 | 2222581 | 71708 | 18.73 | 0.0044 | 0.0006 | 28615 | 0.0023 | 1356 | 0.37 |
| 45 | Z36778_pamp_BE_U | BUD | plate 3 | 1 | 0.51 | 1707602 | 1707590 | 56554 | 16.65 | 0.0044 | 0.0006 | 24555 | 0.0022 | 1375 | 0.37 |
| 46 | Z36779_pamp_BE_U | BUD | plate 3 | 1 | 1.87 | 6238577 | 6238545 | 75289 | 17.85 | 0.0061 | 0.0007 | 28772 | 0.0023 | 1398 | 0.36 |
| 47 | Z36780_pamp_BE_U | BUD | plate 3 | 1 | 1.16 | 3859701 | 3859682 | 66395 | 25.44 | 0.0040 | 0.0006 | 26283 | 0.0019 | 1388 | 0.36 |
| 48 | Z36781_pamp_BE_U | BUD | plate 3 | 1 | 0.97 | 3226183 | 3226164 | 78968 | 20.86 | 0.0051 | 0.0006 | 30234 | 0.0023 | 1418 | 0.35 |
| 49 | Z36782_pamp_BE_U | BUD | plate 3 | 1 | 0.55 | 1840001 | 1839995 | 76741 | 14.60 | 0.0044 | 0.0006 | 28842 | 0.0022 | 1416 | 0.35 |
| 50 | Z36799_pamp_BE_U | JOS | plate 3 | 1 | 0.85 | 2849627 | 2849607 | 81911 | 22.49 | 0.0048 | 0.0006 | 34236 | 0.0023 | 1404 | 0.35 |
| 51 | Z36800_pamp_BE_U | JOS | plate 3 | 1 | 0.92 | 3076507 | 3076496 | 97432 | 19.30 | 0.0046 | 0.0006 | 36391 | 0.0021 | 1323 | 0.38 |
| 52 | Z36801_pamp_BE_U | JOS | plate 3 | 1 | 0.84 | 2788536 | 2788519 | 79600 | 19.96 | 0.0044 | 0.0006 | 32156 | 0.0022 | 1311 | 0.39 |
| 53 | Z36802_pamp_BE_U | JOS | plate 3 | 1 | 0.93 | 3083722 | 3083700 | 79903 | 18.96 | 0.0049 | 0.0006 | 31794 | 0.0023 | 1241 | 0.41 |

| | | | | | | | | | | | | | | | |
|---|---|---|---|---|---|---|---|---|---|---|---|---|---|---|---|
| 54 | Z36803_pamp_BE_U | JOS | plate 3 | 1 | 0.61 | 2030605 | 2030590 | 68651 | 18.29 | 0.0044 | 0.0006 | 30370 | 0.0024 | 1353 | 0.37 |
| 55 | Z36804_pamp_BE_U | JOS | plate 3 | 1 | 0.52 | 1736944 | 1736934 | 61790 | 18.58 | 0.0042 | 0.0006 | 28965 | 0.0023 | 1419 | 0.34 |
| 56 | Z36805_pamp_BE_U | JOS | plate 3 | 1 | 0.44 | 1465887 | 1465872 | 52236 | 18.90 | 0.0047 | 0.0006 | 26721 | 0.0027 | 1450 | 0.33 |
| 57 | Z36806_pamp_BE_U | JOS | plate 3 | 1 | 0.78 | 2597221 | 2597204 | 61350 | 30.09 | 0.0049 | 0.0005 | 29025 | 0.0025 | 1448 | 0.33 |
| 58 | Z36807_pamp_BE_U | JOS | plate 3 | 1 | 0.58 | 1932366 | 1932357 | 61163 | 17.26 | 0.0046 | 0.0006 | 28822 | 0.0025 | 1435 | 0.34 |
| 59 | Z36868_pamp_FI_R | VIR | plate 5 | 2 | 0.74 | 2465270 | 2464972 | 67138 | 22.01 | 0.0039 | 0.0005 | 24994 | 0.0019 | 1325 | 0.38 |
| 60 | Z36881_pamp_FI_R | LOF | plate 4 | 1 | 0.64 | 2122673 | 2122659 | 80700 | 14.37 | 0.0044 | 0.0006 | 25774 | 0.0023 | 1387 | 0.35 |
| 61 | Z36882_pamp_FI_R | VIR | plate 5 | 2 | 0.60 | 2006109 | 2005774 | 69010 | 19.00 | 0.0044 | 0.0005 | 25350 | 0.0020 | 1437 | 0.33 |
| 62 | Z36883_pamp_FI_R | LOF | plate 4 | 1 | 0.57 | 1889535 | 1889521 | 73495 | 13.42 | 0.0040 | 0.0007 | 25426 | 0.0024 | 1382 | 0.36 |
| 63 | Z36884_pamp_FI_R | LOF | plate 4 | 1 | 0.66 | 2208387 | 2208377 | 71335 | 16.94 | 0.0044 | 0.0006 | 25057 | 0.0024 | 1428 | 0.35 |
| 64 | Z36885_pamp_FI_R | LOF | plate 4 | 1 | 0.59 | 1959117 | 1959107 | 63199 | 18.11 | 0.0044 | 0.0006 | 24114 | 0.0025 | 1480 | 0.32 |
| 65 | Z36886_pamp_FI_R | LOF | plate 4 | 1 | 0.78 | 2615709 | 2615699 | 110720 | 13.72 | 0.0042 | 0.0006 | 29747 | 0.0021 | 1289 | 0.41 |
| 66 | Z36887_pamp_FI_R | LOF | plate 4 | 1 | 0.62 | 2074850 | 2074837 | 62551 | 21.45 | 0.0042 | 0.0006 | 24329 | 0.0025 | 1438 | 0.34 |
| 67 | Z36888_pamp_FI_R | LOF | plate 4 | 1 | 0.59 | 1960786 | 1960779 | 85332 | 13.19 | 0.0045 | 0.0006 | 27512 | 0.0022 | 1437 | 0.34 |
| 68 | Z36889_pamp_FI_R | LOF | plate 4 | 1 | 0.52 | 1741595 | 1741584 | 65861 | 15.88 | 0.0043 | 0.0006 | 24742 | 0.0024 | 1450 | 0.34 |
| 69 | Z36898_pamp_FI_U | TUR | plate 4 | 1 | 0.66 | 2215997 | 2215984 | 72066 | 18.74 | 0.0041 | 0.0006 | 28490 | 0.0021 | 1482 | 0.33 |
| 70 | Z36899_pamp_FI_U | PIK | plate 4 | 1 | 0.27 | 898550 | 898545 | 32087 | 20.62 | 0.0049 | 0.0008 | 17340 | 0.0024 | 1277 | 0.41 |
| 71 | Z36901_pamp_FI_U | PIK | plate 4 | 1 | 0.59 | 1950817 | 1950802 | 64385 | 19.65 | 0.0040 | 0.0006 | 24680 | 0.0023 | 1471 | 0.33 |
| 72 | Z36902_pamp_FI_R | LOF | plate 4 | 1 | 0.61 | 2047421 | 2047408 | 73562 | 17.57 | 0.0041 | 0.0006 | 26538 | 0.0023 | 1477 | 0.32 |
| 73 | Z36906_pamp_FI_R | VIR | plate 5 | 2 | 0.51 | 1708198 | 1707939 | 51367 | 18.54 | 0.0040 | 0.0005 | 22512 | 0.0019 | 1417 | 0.34 |
| 74 | Z36907_pamp_FI_R | VIR | plate 5 | 2 | 0.68 | 2272107 | 2271818 | 63346 | 19.66 | 0.0043 | 0.0005 | 24161 | 0.0021 | 1409 | 0.34 |
| 75 | Z36908_pamp_FI_R | VIR | plate 5 | 2 | 0.87 | 2909591 | 2909197 | 80135 | 20.90 | 0.0064 | 0.0005 | 27076 | 0.0019 | 1348 | 0.37 |
| 76 | Z36909_pamp_FI_R | VIR | plate 5 | 2 | 0.91 | 3028049 | 3027646 | 60672 | 32.56 | 0.0039 | 0.0005 | 23600 | 0.0018 | 1422 | 0.34 |
| 77 | Z36910_pamp_FI_U | PIK | plate 4 | 1 | 0.71 | 2366015 | 2366004 | 68423 | 21.66 | 0.0041 | 0.0006 | 24921 | 0.0022 | 1483 | 0.32 |
| 78 | Z36911_pamp_FI_U | TUR | plate 4 | 1 | 0.65 | 2156025 | 2156006 | 69990 | 18.48 | 0.0040 | 0.0006 | 27895 | 0.0021 | 1430 | 0.34 |
| 79 | Z36916_pamp_FI_R | VIR | plate 5 | 2 | 0.71 | 2350707 | 2350361 | 64882 | 20.03 | 0.0037 | 0.0006 | 24634 | 0.0019 | 1382 | 0.36 |
| 80 | Z36923_pamp_FI_U | PIK | plate 4 | 1 | 0.56 | 1881790 | 1881774 | 59880 | 19.63 | 0.0041 | 0.0006 | 23381 | 0.0023 | 1388 | 0.37 |
| 81 | Z36924_pamp_FI_U | TUR | plate 4 | 1 | 0.46 | 1545642 | 1545631 | 54726 | 18.31 | 0.0042 | 0.0006 | 24906 | 0.0022 | 1469 | 0.33 |
| 82 | Z36925_pamp_FI_U | PIK | plate 4 | 1 | 0.48 | 1613803 | 1613793 | 62679 | 16.57 | 0.0041 | 0.0006 | 23778 | 0.0023 | 1444 | 0.34 |
| 83 | Z36926_pamp_FI_U | TUR | plate 4 | 1 | 0.65 | 2165296 | 2165279 | 69110 | 20.69 | 0.0042 | 0.0006 | 27550 | 0.0022 | 1423 | 0.34 |
| 84 | Z36927_pamp_FI_U | PIK | plate 4 | 1 | 0.36 | 1198684 | 1198679 | 44408 | 17.98 | 0.0045 | 0.0007 | 22777 | 0.0024 | 1510 | 0.31 |
| 85 | Z36928_pamp_FI_R | LOF | plate 4 | 1 | 1.22 | 4067108 | 4067086 | 75469 | 40.08 | 0.0046 | 0.0006 | 26718 | 0.0024 | 1424 | 0.33 |

| | | | | | | | | | | | | | | | |
|---|---|---|---|---|---|---|---|---|---|---|---|---|---|---|---|
| 86 | Z36929_pamp_FI_R | LOF | plate 4 | 1 | 0.96 | 3203802 | 3203783 | 78750 | 22.95 | 0.0038 | 0.0006 | 28469 | 0.0022 | 1440 | 0.35 |
| 87 | Z36930_pamp_FI_R | LOF | plate 4 | 1 | 0.45 | 1509173 | 1509163 | 58176 | 16.48 | 0.0043 | 0.0006 | 24929 | 0.0025 | 1494 | 0.32 |
| 88 | Z36931_pamp_FI_R | LOF | plate 4 | 1 | 0.70 | 2328165 | 2328152 | 71428 | 21.43 | 0.0034 | 0.0006 | 31784 | 0.0019 | 1429 | 0.35 |
| 89 | Z36932_pamp_FI_R | LOF | plate 4 | 1 | 0.47 | 1557956 | 1557952 | 22424 | 50.78 | 0.0041 | 0.0006 | 7739 | 0.0010 | 619 | 0.73 |
| 90 | Z36933_pamp_FI_R | LOF | plate 4 | 1 | 0.40 | 1341739 | 1341734 | 51662 | 14.17 | 0.0044 | 0.0007 | 23700 | 0.0026 | 1475 | 0.33 |
| 91 | Z36934_pamp_FI_R | LOF | plate 4 | 1 | 0.39 | 1310675 | 1310668 | 53709 | 15.45 | 0.0046 | 0.0006 | 21933 | 0.0025 | 1469 | 0.33 |
| 92 | Z36940_pamp_CZ_R | MIL | plate 4 | 1 | 0.41 | 1374174 | 1374170 | 57803 | 15.86 | 0.0054 | 0.0006 | 24334 | 0.0030 | 1075 | 0.49 |
| 93 | Z36941_pamp_CZ_R | MIL | plate 4 | 1 | 0.53 | 1750744 | 1750726 | 57875 | 20.26 | 0.0052 | 0.0006 | 26078 | 0.0030 | 1145 | 0.45 |
| 94 | Z36942_pamp_CZ_R | MIL | plate 4 | 1 | 0.68 | 2255323 | 2255310 | 77307 | 19.63 | 0.0051 | 0.0006 | 28482 | 0.0028 | 1072 | 0.48 |
| 95 | Z36943_pamp_CZ_R | MIL | plate 4 | 1 | 0.73 | 2446631 | 2446620 | 71773 | 16.59 | 0.0053 | 0.0006 | 27203 | 0.0028 | 1114 | 0.46 |
| 96 | Z36944_pamp_CZ_R | MIL | plate 4 | 1 | 0.80 | 2658419 | 2658398 | 68078 | 19.17 | 0.0055 | 0.0006 | 27360 | 0.0028 | 1137 | 0.45 |
| 97 | Z36945_pamp_CZ_R | MIL | plate 4 | 1 | 0.71 | 2356789 | 2356778 | 73054 | 20.72 | 0.0051 | 0.0006 | 28154 | 0.0028 | 1102 | 0.47 |
| 98 | Z36946_pamp_CZ_R | MIL | plate 4 | 1 | 0.59 | 1964824 | 1964811 | 62343 | 20.37 | 0.0052 | 0.0005 | 26953 | 0.0030 | 1136 | 0.45 |
| 99 | Z36947_pamp_CZ_R | MIL | plate 4 | 1 | 0.60 | 2004565 | 2004551 | 62166 | 19.67 | 0.0054 | 0.0005 | 26291 | 0.0030 | 1054 | 0.46 |
| 100 | Z36948_pamp_CZ_U | POH | plate 4 | 1 | 0.54 | 1808032 | 1808027 | 59025 | 18.49 | 0.0053 | 0.0006 | 25993 | 0.0029 | 1162 | 0.43 |
| 101 | Z36949_pamp_CZ_U | POH | plate 4 | 1 | 0.91 | 3045291 | 3045269 | 74616 | 18.15 | 0.0056 | 0.0006 | 28191 | 0.0028 | 1145 | 0.44 |
| 102 | Z36950_pamp_CZ_U | POH | plate 4 | 1 | 0.74 | 2454364 | 2454348 | 83576 | 18.42 | 0.0050 | 0.0006 | 29147 | 0.0027 | 1072 | 0.45 |
| 103 | Z36951_pamp_CZ_U | POH | plate 4 | 1 | 0.34 | 1136738 | 1136729 | 44617 | 16.99 | 0.0053 | 0.0006 | 25173 | 0.0029 | 1150 | 0.43 |
| 104 | Z36952_pamp_CZ_U | POH | plate 4 | 1 | 0.63 | 2104036 | 2104024 | 62130 | 20.10 | 0.0049 | 0.0006 | 29391 | 0.0027 | 1144 | 0.45 |
| 105 | Z36953_pamp_CZ_U | POH | plate 4 | 1 | 0.87 | 2910007 | 2909989 | 69542 | 22.49 | 0.0054 | 0.0006 | 31093 | 0.0027 | 1179 | 0.42 |
| 106 | Z36954_pamp_CZ_U | POH | plate 4 | 1 | 0.85 | 2830537 | 2830517 | 76953 | 21.15 | 0.0058 | 0.0006 | 32644 | 0.0026 | 1168 | 0.44 |
| 107 | Z36955_pamp_CZ_U | POH | plate 4 | 1 | 0.84 | 2793585 | 2793564 | 66648 | 20.18 | 0.0052 | 0.0006 | 30711 | 0.0026 | 1230 | 0.42 |
| 108 | Z36963_pamp_CZ_U | POH | plate 4 | 1 | 0.61 | 2040491 | 2040486 | 61060 | 18.35 | 0.0053 | 0.0006 | 25819 | 0.0029 | 1133 | 0.45 |
| 109 | Z36964_pamp_CZ_U | POH | plate 4 | 1 | 0.43 | 1417212 | 1417206 | 50361 | 18.58 | 0.0050 | 0.0006 | 24377 | 0.0030 | 1181 | 0.42 |
| 110 | Z36965_pamp_CZ_U | POH | plate 4 | 1 | 0.70 | 2342180 | 2342162 | 62810 | 15.05 | 0.0055 | 0.0006 | 25879 | 0.0028 | 1040 | 0.47 |
| 111 | Z36966_pamp_CZ_U | POH | plate 4 | 1 | 0.47 | 1551036 | 1551029 | 59774 | 17.57 | 0.0050 | 0.0006 | 26162 | 0.0028 | 1170 | 0.44 |
| 112 | Z36967_pamp_CZ_U | POH | plate 4 | 1 | 0.67 | 2217972 | 2217964 | 60958 | 16.98 | 0.0054 | 0.0006 | 26375 | 0.0029 | 1177 | 0.43 |
| 113 | Z36968_pamp_CZ_U | POH | plate 4 | 1 | 0.88 | 2931872 | 2931859 | 57205 | 18.77 | 0.0053 | 0.0006 | 25788 | 0.0029 | 1173 | 0.44 |
| 114 | Z36969_pamp_CZ_U | POH | plate 4 | 1 | 0.55 | 1827486 | 1827473 | 66382 | 17.13 | 0.0051 | 0.0006 | 27043 | 0.0027 | 1127 | 0.45 |
| 115 | Z36970_pamp_CZ_U | POH | plate 4 | 1 | 0.50 | 1655701 | 1655691 | 52952 | 18.63 | 0.0054 | 0.0006 | 24935 | 0.0031 | 1104 | 0.44 |
| 116 | Z36972_pamp_CZ_U | DIV | plate 3 | 1 | 0.99 | 3298217 | 3298191 | 60355 | 26.15 | 0.0054 | 0.0006 | 32292 | 0.0024 | 1181 | 0.42 |
| 117 | Z36973_pamp_CZ_U | DIV | plate 3 | 1 | 0.76 | 2525092 | 2525082 | 48474 | 23.76 | 0.0057 | 0.0007 | 27772 | 0.0026 | 1237 | 0.39 |

| | | | | | | | | | | | | | | | |
|---|---|---|---|---|---|---|---|---|---|---|---|---|---|---|---|
| 118 | Z36974_pamp_CZ_U | DIV | plate 4 | 1 | 0.54 | 1784885 | 1784880 | 57863 | 20.65 | 0.0055 | 0.0006 | 29008 | 0.0029 | 1091 | 0.49 |
| 119 | Z36975_pamp_CZ_U | DIV | plate 4 | 1 | 0.44 | 1462548 | 1462538 | 48414 | 16.93 | 0.0058 | 0.0006 | 26440 | 0.0030 | 1096 | 0.48 |
| 120 | Z36976_pamp_CZ_U | DIV | plate 4 | 1 | 0.43 | 1424356 | 1424351 | 56936 | 16.90 | 0.0055 | 0.0005 | 28273 | 0.0028 | 1066 | 0.49 |
| 121 | Z36977_pamp_CZ_U | DIV | plate 4 | 1 | 0.81 | 2684184 | 2684172 | 53478 | 18.18 | 0.0065 | 0.0006 | 27517 | 0.0029 | 1101 | 0.49 |
| 122 | Z36978_pamp_CZ_U | DIV | plate 4 | 1 | 0.87 | 2908785 | 2908777 | 67141 | 18.15 | 0.0062 | 0.0006 | 30512 | 0.0028 | 1098 | 0.48 |
| 123 | Z36979_pamp_CZ_U | DIV | plate 4 | 1 | 0.83 | 2770959 | 2770950 | 69305 | 21.11 | 0.0055 | 0.0005 | 31541 | 0.0028 | 1101 | 0.48 |
| 124 | Z36980_pamp_CZ_U | DIV | plate 4 | 1 | 0.60 | 1985344 | 1985333 | 57430 | 17.33 | 0.0059 | 0.0006 | 28597 | 0.0030 | 1103 | 0.48 |
| 125 | Z36981_pamp_CZ_U | DIV | plate 4 | 1 | 1.03 | 3427769 | 3427757 | 74147 | 20.93 | 0.0061 | 0.0006 | 32020 | 0.0028 | 1114 | 0.47 |
| 126 | Z36982_pamp_CZ_U | DIV | plate 4 | 1 | 0.87 | 2906189 | 2906179 | 73612 | 20.03 | 0.0058 | 0.0006 | 31985 | 0.0028 | 1047 | 0.49 |
| 127 | Z36983_pamp_CZ_U | DIV | plate 4 | 1 | 0.79 | 2627883 | 2627876 | 73418 | 19.25 | 0.0059 | 0.0005 | 29435 | 0.0028 | 1116 | 0.47 |
| 128 | Z36984_pamp_CZ_U | DIV | plate 4 | 1 | 0.96 | 3201304 | 3201291 | 69869 | 20.81 | 0.0061 | 0.0006 | 29013 | 0.0030 | 1107 | 0.47 |
| 129 | Z36985_pamp_CZ_R | MIL | plate 4 | 1 | 0.65 | 2156845 | 2156831 | 71471 | 15.94 | 0.0052 | 0.0006 | 27524 | 0.0028 | 1098 | 0.48 |
| 130 | Z37005_pamp_CZ_R | MIL | plate 4 | 1 | 0.82 | 2747368 | 2747358 | 61622 | 17.39 | 0.0063 | 0.0006 | 27613 | 0.0030 | 1077 | 0.49 |
| 131 | Z37006_pamp_CZ_R | MIL | plate 4 | 1 | 0.65 | 2172112 | 2172102 | 70283 | 15.27 | 0.0056 | 0.0006 | 29512 | 0.0027 | 1104 | 0.48 |
| 132 | Z37007_pamp_CZ_R | MIL | plate 4 | 1 | 0.67 | 2218255 | 2218247 | 72897 | 17.32 | 0.0052 | 0.0006 | 29297 | 0.0027 | 1073 | 0.48 |
| 133 | Z37008_pamp_CZ_R | MIL | plate 4 | 1 | 0.76 | 2521728 | 2521717 | 65029 | 26.17 | 0.0054 | 0.0006 | 28064 | 0.0029 | 1080 | 0.49 |
| 134 | Z37009_pamp_CZ_R | MIL | plate 4 | 1 | 0.44 | 1478681 | 1478678 | 49278 | 19.96 | 0.0055 | 0.0006 | 25646 | 0.0031 | 1006 | 0.49 |
| 135 | Z37010_pamp_CZ_R | MIL | plate 4 | 1 | 0.51 | 1698623 | 1698618 | 60562 | 16.83 | 0.0054 | 0.0006 | 27323 | 0.0030 | 1043 | 0.50 |
| 136 | Z37011_pamp_CZ_R | MIL | plate 4 | 1 | 0.71 | 2375035 | 2375026 | 65375 | 19.61 | 0.0056 | 0.0006 | 28358 | 0.0029 | 1095 | 0.48 |
| 137 | Z37053_pamp_SE_R | SON | plate 5 | 2 | 1.13 | 3779219 | 3778677 | 71237 | 31.28 | 0.0046 | 0.0005 | 35794 | 0.0019 | 1403 | 0.35 |
| 138 | Z37054_pamp_SE_R | SON | plate 5 | 2 | 0.90 | 2989445 | 2989024 | 67492 | 28.76 | 0.0047 | 0.0005 | 34917 | 0.0020 | 1443 | 0.33 |
| 139 | Z37055_pamp_SE_U | GAR | plate 5 | 2 | 0.56 | 1873066 | 1872813 | 81896 | 14.34 | 0.0048 | 0.0005 | 28759 | 0.0022 | 1343 | 0.37 |
| 140 | Z37056_pamp_SE_U | GAR | plate 5 | 2 | 0.67 | 2218218 | 2217893 | 64715 | 20.01 | 0.0046 | 0.0005 | 26745 | 0.0023 | 1475 | 0.32 |
| 141 | Z37057_pamp_SE_U | GAR | plate 5 | 2 | 0.86 | 2862509 | 2862051 | 83497 | 20.22 | 0.0047 | 0.0006 | 29691 | 0.0022 | 1338 | 0.38 |
| 142 | Z37058_pamp_SE_U | GAR | plate 5 | 2 | 0.41 | 1376640 | 1376422 | 56046 | 15.37 | 0.0051 | 0.0006 | 24607 | 0.0024 | 1466 | 0.31 |
| 143 | Z37059_pamp_SE_U | GAR | plate 5 | 2 | 0.55 | 1826240 | 1825981 | 64496 | 16.55 | 0.0051 | 0.0006 | 26634 | 0.0023 | 1465 | 0.32 |
| 144 | Z37060_pamp_SE_U | GAR | plate 5 | 2 | 0.74 | 2472065 | 2471709 | 74998 | 17.02 | 0.0045 | 0.0005 | 28170 | 0.0022 | 1374 | 0.37 |
| 145 | Z37061_pamp_SE_U | GAR | plate 5 | 2 | 0.72 | 2415684 | 2415366 | 66972 | 19.82 | 0.0049 | 0.0005 | 27118 | 0.0023 | 1370 | 0.36 |
| 146 | Z37063_pamp_SE_U | BRO | plate 5 | 2 | 0.93 | 3105834 | 3105387 | 69041 | 27.44 | 0.0044 | 0.0006 | 26684 | 0.0022 | 1373 | 0.36 |
| 147 | Z37064_pamp_SE_U | BRO | plate 5 | 2 | 0.25 | 823955 | 823818 | 50133 | 11.64 | 0.0049 | 0.0006 | 21872 | 0.0023 | 1381 | 0.35 |
| 148 | Z37065_pamp_SE_U | BRO | plate 5 | 2 | 0.94 | 3140867 | 3140446 | 83971 | 23.95 | 0.0044 | 0.0005 | 29438 | 0.0021 | 1295 | 0.39 |
| 149 | Z37066_pamp_SE_U | BRO | plate 5 | 2 | 0.27 | 888059 | 887958 | 39989 | 14.44 | 0.0050 | 0.0006 | 20494 | 0.0024 | 1463 | 0.32 |

| | | | | | | | | | | | | | | | |
|---|---|---|---|---|---|---|---|---|---|---|---|---|---|---|---|
| 150 | Z37067_pamp_SE_U | BRO | plate 5 | 2 | 0.68 | 2266540 | 2266238 | 68148 | 20.23 | 0.0046 | 0.0005 | 26350 | 0.0022 | 1291 | 0.39 |
| 151 | Z37068_pamp_SE_U | BRO | plate 5 | 2 | 0.64 | 2141883 | 2141633 | 53743 | 29.78 | 0.0045 | 0.0005 | 23823 | 0.0023 | 1457 | 0.31 |
| 152 | Z37069_pamp_SE_U | BRO | plate 5 | 2 | 0.71 | 2375742 | 2375434 | 89978 | 16.68 | 0.0064 | 0.0005 | 31710 | 0.0020 | 1368 | 0.36 |
| 153 | Z37070_pamp_SE_U | BRO | plate 5 | 2 | 0.55 | 1841349 | 1841084 | 72955 | 15.19 | 0.0044 | 0.0006 | 27070 | 0.0021 | 1360 | 0.36 |
| 154 | Z37071_pamp_SE_U | BRO | plate 5 | 2 | 0.73 | 2431378 | 2431054 | 73102 | 18.36 | 0.0046 | 0.0005 | 30003 | 0.0023 | 1326 | 0.38 |
| 155 | Z37072_pamp_SE_U | BRO | plate 5 | 2 | 0.47 | 1551516 | 1551306 | 62955 | 13.64 | 0.0043 | 0.0006 | 26029 | 0.0020 | 1316 | 0.39 |
| 156 | Z37079_pamp_SE_U | BRO | plate 5 | 2 | 0.61 | 2031927 | 2031662 | 77813 | 15.00 | 0.0048 | 0.0005 | 29048 | 0.0023 | 1347 | 0.37 |
| 157 | Z37082_pamp_SE_R | SON | plate 5 | 2 | 1.01 | 3372222 | 3371681 | 80214 | 25.41 | 0.0044 | 0.0005 | 38384 | 0.0019 | 1392 | 0.35 |
| 158 | Z37083_pamp_SE_R | SON | plate 5 | 2 | 0.68 | 2283149 | 2282794 | 62617 | 22.40 | 0.0045 | 0.0005 | 32178 | 0.0019 | 1429 | 0.34 |
| 159 | Z37084_pamp_SE_R | SON | plate 5 | 2 | 0.67 | 2218414 | 2218076 | 56145 | 25.37 | 0.0049 | 0.0005 | 30184 | 0.0020 | 1447 | 0.33 |
| 160 | Z37085_pamp_SE_R | SON | plate 5 | 2 | 0.44 | 1466603 | 1466360 | 44685 | 21.57 | 0.0042 | 0.0005 | 24593 | 0.0020 | 1449 | 0.33 |
| 161 | Z37086_pamp_SE_R | SON | plate 5 | 2 | 0.78 | 2599601 | 2599226 | 68184 | 22.65 | 0.0064 | 0.0005 | 30395 | 0.0021 | 1417 | 0.34 |
| 162 | Z37087_pamp_SE_R | SON | plate 5 | 2 | 0.92 | 3051541 | 3051201 | 118112 | 15.87 | 0.0046 | 0.0005 | 43211 | 0.0016 | 1445 | 0.32 |
| 163 | Z37088_pamp_SE_R | SON | plate 5 | 2 | 0.38 | 1255455 | 1255253 | 32113 | 9.86 | 0.0055 | 0.0007 | 18051 | 0.0021 | 1284 | 0.41 |
| 164 | Z37089_pamp_SE_R | SON | plate 5 | 2 | 1.01 | 3352824 | 3352345 | 76107 | 21.69 | 0.0045 | 0.0005 | 31858 | 0.0019 | 1347 | 0.37 |
| 165 | Z37090_pamp_SE_R | SON | plate 5 | 2 | 0.86 | 2861332 | 2860937 | 71601 | 19.42 | 0.0044 | 0.0005 | 30267 | 0.0020 | 1343 | 0.38 |
| 166 | Z37102_pamp_SE_U | GAR | plate 5 | 2 | 0.35 | 1164066 | 1163898 | 43430 | 17.29 | 0.0056 | 0.0007 | 19663 | 0.0020 | 1211 | 0.42 |
| 167 | Z37103_pamp_SE_U | GAR | plate 5 | 2 | 0.41 | 1377292 | 1377109 | 47782 | 15.45 | 0.0050 | 0.0006 | 23968 | 0.0024 | 1449 | 0.32 |
| 168 | Z37105_pamp_SE_R | SON | plate 5 | 2 | 0.23 | 769134 | 769019 | 40883 | 12.88 | 0.0046 | 0.0006 | 20978 | 0.0023 | 1454 | 0.32 |
| 169 | Z37106_pamp_SE_R | SON | plate 5 | 2 | 0.39 | 1288684 | 1288523 | 47682 | 16.88 | 0.0049 | 0.0006 | 27258 | 0.0021 | 1436 | 0.33 |
| 170 | Z37107_pamp_SE_R | SON | plate 5 | 2 | 0.58 | 1917313 | 1916949 | 65950 | 19.35 | 0.0049 | 0.0005 | 32943 | 0.0020 | 1467 | 0.32 |
| 171 | Z37108_pamp_SE_R | SON | plate 5 | 2 | 0.80 | 2653311 | 2652946 | 63875 | 23.84 | 0.0045 | 0.0004 | 33171 | 0.0020 | 1414 | 0.34 |
| 172 | Z37109_pamp_SE_R | SON | plate 5 | 2 | 0.76 | 2536995 | 2536671 | 65239 | 21.81 | 0.0046 | 0.0005 | 33145 | 0.0019 | 1433 | 0.34 |
| 173 | Z37110_pamp_SE_U | BRO | plate 5 | 2 | 0.46 | 1529882 | 1529668 | 63564 | 13.65 | 0.0045 | 0.0006 | 25608 | 0.0021 | 1353 | 0.36 |
| 174 | Z37111_pamp_SE_U | BRO | plate 5 | 2 | 0.69 | 2299930 | 2299616 | 68857 | 15.43 | 0.0045 | 0.0005 | 26470 | 0.0022 | 1237 | 0.42 |
| 175 | Z37112_pamp_SE_U | BRO | plate 5 | 2 | 0.52 | 1734163 | 1733942 | 19348 | 31.28 | 0.0072 | 0.0005 | 6556 | 0.0012 | 192 | 0.90 |
| 176 | Z37113_pamp_SE_U | BRO | plate 5 | 2 | 0.56 | 1855948 | 1855702 | 65291 | 16.87 | 0.0045 | 0.0006 | 24859 | 0.0022 | 1297 | 0.38 |
| 177 | Z37114_pamp_SE_U | BRO | plate 5 | 2 | 0.71 | 2359141 | 2358836 | 66724 | 21.83 | 0.0044 | 0.0005 | 26945 | 0.0022 | 1360 | 0.37 |
| 178 | Z37115_pamp_BE_R | NIS | plate 3 | 1 | 1.09 | 3628349 | 3628326 | 78208 | 26.83 | 0.0045 | 0.0006 | 36763 | 0.0019 | 1331 | 0.39 |
| 179 | Z37116_pamp_BE_R | NIS | plate 3 | 1 | 0.46 | 1527309 | 1527299 | 46366 | 22.36 | 0.0044 | 0.0006 | 26871 | 0.0024 | 1444 | 0.32 |
| 180 | Z37117_pamp_BE_R | NIS | plate 3 | 1 | 0.75 | 2491384 | 2491364 | 43427 | 23.77 | 0.0052 | 0.0007 | 26123 | 0.0026 | 1511 | 0.30 |
| 181 | Z37118_pamp_BE_R | NIS | plate 3 | 1 | 0.54 | 1800599 | 1800586 | 52080 | 24.17 | 0.0047 | 0.0006 | 28741 | 0.0024 | 1404 | 0.34 |

| | | | | | | | | | | | | | | | |
|---|---|---|---|---|---|---|---|---|---|---|---|---|---|---|---|
| 182 | Z37119_pamp_BE_R | NIS | plate 3 | 1 | 0.62 | 2054628 | 2054616 | 48263 | 20.35 | 0.0048 | 0.0007 | 26759 | 0.0024 | 1431 | 0.33 |
| 183 | Z37120_pamp_BE_R | NIS | plate 3 | 1 | 0.65 | 2175850 | 2175838 | 57810 | 24.81 | 0.0044 | 0.0006 | 30548 | 0.0023 | 1443 | 0.33 |
| 184 | Z37121_pamp_BE_R | NIS | plate 3 | 1 | 0.44 | 1479754 | 1479745 | 45435 | 21.98 | 0.0048 | 0.0006 | 26151 | 0.0025 | 1371 | 0.34 |
| 185 | Z37122_pamp_BE_R | NIS | plate 3 | 1 | 0.69 | 2315904 | 2315892 | 68396 | 21.79 | 0.0044 | 0.0006 | 33566 | 0.0021 | 1324 | 0.38 |
| 186 | Z37139_pamp_BE_R | NIS | plate 3 | 1 | 0.65 | 2157450 | 2157436 | 61844 | 22.30 | 0.0042 | 0.0007 | 30881 | 0.0021 | 1400 | 0.36 |
| 187 | Z37140_pamp_BE_R | NIS | plate 3 | 1 | 0.42 | 1415711 | 1415698 | 53235 | 16.64 | 0.0042 | 0.0006 | 28419 | 0.0022 | 1433 | 0.34 |
| 188 | Z37141_pamp_BE_R | NIS | plate 3 | 1 | 0.19 | 620469 | 620467 | 29203 | 16.22 | 0.0045 | 0.0007 | 15286 | 0.0016 | 839 | 0.61 |
| 189 | Z37142_pamp_BE_R | NIS | plate 3 | 1 | 0.81 | 2693257 | 2693240 | 76308 | 21.36 | 0.0044 | 0.0006 | 34613 | 0.0020 | 1334 | 0.38 |
| 190 | Z37143_pamp_BE_R | NIS | plate 3 | 1 | 0.89 | 2981186 | 2981171 | 88826 | 17.42 | 0.0042 | 0.0006 | 35413 | 0.0018 | 1127 | 0.46 |
| 191 | Z37144_pamp_BE_R | NIS | plate 3 | 1 | 0.71 | 2371734 | 2371719 | 70386 | 19.89 | 0.0045 | 0.0006 | 32690 | 0.0021 | 1343 | 0.37 |
| 192 | Z37145_pamp_BE_R | NIS | plate 3 | 1 | 0.82 | 2723231 | 2723223 | 75127 | 22.86 | 0.0042 | 0.0006 | 33516 | 0.0020 | 1230 | 0.41 |
| 193 | Z37146_pamp_BE_R | NIS | plate 3 | 1 | 0.56 | 1863545 | 1863532 | 58886 | 20.95 | 0.0046 | 0.0007 | 28860 | 0.0022 | 1304 | 0.36 |
| 194 | Z38283_pamp_BE_U | BUD | plate 3 | 1 | 0.74 | 2457337 | 2457322 | 73709 | 21.32 | 0.0042 | 0.0006 | 28649 | 0.0024 | 1360 | 0.37 |
| 195 | Z38284_pamp_BE_U | BUD | plate 3 | 1 | 0.71 | 2370583 | 2370561 | 87454 | 15.69 | 0.0045 | 0.0006 | 29521 | 0.0021 | 1210 | 0.44 |
| 196 | Z38285_pamp_BE_U | BUD | plate 3 | 1 | 0.42 | 1414727 | 1414719 | 50742 | 17.96 | 0.0046 | 0.0006 | 24493 | 0.0026 | 1430 | 0.34 |
| 197 | Z38286_pamp_BE_U | BUD | plate 3 | 1 | 0.67 | 2236381 | 2236374 | 80663 | 16.30 | 0.0046 | 0.0006 | 33002 | 0.0021 | 1355 | 0.37 |
| 198 | Z38287_pamp_BE_U | BUD | plate 3 | 1 | 0.91 | 3049684 | 3049663 | 88579 | 19.23 | 0.0046 | 0.0006 | 35531 | 0.0020 | 1241 | 0.41 |
| 199 | Z38288_pamp_BE_U | BUD | plate 3 | 1 | 0.67 | 2246174 | 2246158 | 78200 | 18.28 | 0.0041 | 0.0006 | 33043 | 0.0021 | 1395 | 0.36 |
| 200 | Z38289_pamp_BE_U | BUD | plate 3 | 1 | 0.90 | 3001379 | 3001370 | 72387 | 21.52 | 0.0045 | 0.0006 | 31030 | 0.0022 | 1292 | 0.37 |
| 201 | Z38290_pamp_BE_U | BUD | plate 3 | 1 | 0.80 | 2673647 | 2673632 | 82633 | 20.48 | 0.0046 | 0.0006 | 34281 | 0.0022 | 1376 | 0.37 |
| 202 | Z38294_pamp_BE_U | BUD | plate 3 | 1 | 0.34 | 1133377 | 1133372 | 52886 | 13.96 | 0.0039 | 0.0007 | 23599 | 0.0022 | 1336 | 0.37 |
| 203 | Z38295_pamp_BE_U | BUD | plate 3 | 1 | 0.55 | 1817059 | 1817051 | 56584 | 15.51 | 0.0043 | 0.0006 | 24457 | 0.0024 | 1325 | 0.36 |
| 204 | Z38296_pamp_BE_U | JOS | plate 3 | 1 | 1.06 | 3549432 | 3549409 | 66632 | 21.66 | 0.0047 | 0.0006 | 31033 | 0.0019 | 1381 | 0.37 |
| 205 | Z38297_pamp_BE_U | JOS | plate 3 | 1 | 1.02 | 3402305 | 3402289 | 86819 | 21.33 | 0.0044 | 0.0006 | 35362 | 0.0020 | 1335 | 0.38 |
| 206 | Z38298_pamp_BE_U | JOS | plate 3 | 1 | 0.44 | 1483313 | 1483306 | 72865 | 13.88 | 0.0046 | 0.0006 | 31877 | 0.0022 | 1440 | 0.33 |
| 207 | Z38299_pamp_BE_U | JOS | plate 3 | 1 | 0.80 | 2673872 | 2673860 | 69534 | 22.06 | 0.0045 | 0.0006 | 31760 | 0.0023 | 1418 | 0.35 |
| 208 | Z38307_pamp_BE_U | JOS | plate 3 | 1 | 0.95 | 3157734 | 3157717 | 74798 | 27.29 | 0.0043 | 0.0006 | 33457 | 0.0020 | 1200 | 0.43 |
| 209 | Z67792_pamp_GR_U | GAL | plate 7 | 2 | 0.85 | 2817599 | 2817158 | 49233 | 48.30 | 0.0066 | 0.0007 | 19847 | 0.0031 | 1057 | 0.46 |
| 210 | Z67793_pamp_GR_U | GAL | plate 7 | 2 | 0.49 | 1642188 | 1641949 | 51154 | 22.76 | 0.0069 | 0.0006 | 19356 | 0.0029 | 1023 | 0.48 |
| 211 | Z67794_pamp_GR_U | GAL | plate 7 | 2 | 0.31 | 1038508 | 1038369 | 44645 | 15.41 | 0.0068 | 0.0007 | 17825 | 0.0028 | 998 | 0.50 |
| 212 | Z67795_pamp_GR_U | GAL | plate 7 | 2 | 0.47 | 1563513 | 1563289 | 57717 | 17.88 | 0.0059 | 0.0006 | 21210 | 0.0030 | 1025 | 0.49 |
| 213 | Z67796_pamp_FR_U | PET | plate 7 | 2 | 0.50 | 1672764 | 1672539 | 70314 | 13.08 | 0.0056 | 0.0005 | 24308 | 0.0028 | 1319 | 0.37 |

| | | | | | | | | | | | | | | | |
|---|---|---|---|---|---|---|---|---|---|---|---|---|---|---|---|
| 214 | Z67797_pamp_FR_U | PET | plate 7 | 2 | 0.48 | 1587617 | 1587397 | 57420 | 14.97 | 0.0061 | 0.0006 | 21742 | 0.0028 | 1303 | 0.38 |
| 215 | Z67798_pamp_FR_U | PET | plate 7 | 2 | 0.66 | 2209930 | 2209626 | 69407 | 14.44 | 0.0053 | 0.0006 | 23096 | 0.0026 | 1153 | 0.43 |
| 216 | Z67799_pamp_FR_U | PET | plate 7 | 2 | 0.66 | 2209731 | 2209420 | 55449 | 26.28 | 0.0063 | 0.0006 | 21110 | 0.0029 | 1251 | 0.40 |
| 217 | Z67800_pamp_FR_U | PET | plate 7 | 2 | 0.76 | 2541653 | 2541333 | 50064 | 40.13 | 0.0054 | 0.0005 | 22164 | 0.0024 | 1226 | 0.40 |
| 218 | Z67801_pamp_IT_U | ARR | plate 7 | 2 | 0.49 | 1638381 | 1638149 | 72753 | 13.88 | 0.0053 | 0.0005 | 23795 | 0.0029 | 1001 | 0.50 |
| 219 | Z67802_pamp_IT_R | MAN | plate 7 | 2 | 0.75 | 2511201 | 2510830 | 109976 | 13.68 | 0.0052 | 0.0005 | 27905 | 0.0025 | 874 | 0.55 |
| 220 | Z67803_pamp_IT_R | MAN | plate 7 | 2 | 0.43 | 1442890 | 1442701 | 65126 | 14.36 | 0.0059 | 0.0006 | 24004 | 0.0030 | 1070 | 0.46 |
| 221 | Z67829_pamp_GR_U | GAL | plate 7 | 2 | 1.12 | 3732174 | 3731639 | 48389 | 59.16 | 0.0066 | 0.0006 | 19344 | 0.0029 | 994 | 0.50 |
| 222 | Z67830_pamp_GR_U | GAL | plate 7 | 2 | 0.36 | 1205030 | 1204838 | 47188 | 16.94 | 0.0061 | 0.0007 | 19879 | 0.0029 | 1006 | 0.49 |
| 223 | Z67831_pamp_GR_R | KAT | plate 7 | 2 | 1.78 | 5917140 | 5916385 | 46864 | 108.78 | 0.0059 | 0.0006 | 17273 | 0.0024 | 829 | 0.57 |
| 224 | Z67832_pamp_IT_R | MAN | plate 7 | 2 | 1.24 | 4129847 | 4129234 | 72458 | 40.76 | 0.0056 | 0.0006 | 24715 | 0.0029 | 1066 | 0.47 |
| 225 | Z67833_pamp_IT_R | MAN | plate 7 | 2 | 0.73 | 2434562 | 2434203 | 74384 | 13.80 | 0.0054 | 0.0006 | 24032 | 0.0028 | 964 | 0.51 |
| 226 | Z67834_pamp_IT_R | MAN | plate 7 | 2 | 0.50 | 1667496 | 1667244 | 65975 | 14.65 | 0.0064 | 0.0005 | 24122 | 0.0030 | 1030 | 0.48 |
| 227 | Z67835_pamp_IT_R | MAN | plate 7 | 2 | 0.45 | 1506316 | 1506131 | 60815 | 16.57 | 0.0056 | 0.0005 | 23719 | 0.0030 | 1004 | 0.49 |
| 228 | Z67836_pamp_IT_U | ARR | plate 7 | 2 | 0.73 | 2417141 | 2416774 | 71661 | 17.02 | 0.0055 | 0.0006 | 23648 | 0.0029 | 1025 | 0.49 |
| 229 | Z67837_pamp_IT_U | ARR | plate 7 | 2 | 0.96 | 3197025 | 3196652 | 43125 | 64.57 | 0.0068 | 0.0007 | 19125 | 0.0031 | 945 | 0.51 |
| 230 | Z67838_pamp_IT_U | ARR | plate 7 | 2 | 0.79 | 2649055 | 2648673 | 70742 | 21.57 | 0.0055 | 0.0006 | 23615 | 0.0028 | 1028 | 0.49 |
| 231 | Z67839_pamp_IT_U | ARR | plate 7 | 2 | 0.25 | 827310 | 827189 | 54724 | 9.75 | 0.0062 | 0.0007 | 21355 | 0.0027 | 966 | 0.51 |
| 232 | Z67840_pamp_FR_R | SAV | plate 7 | 2 | 0.37 | 1228471 | 1228298 | 55478 | 10.47 | 0.0064 | 0.0006 | 20112 | 0.0029 | 1235 | 0.42 |
| 233 | Z67841_pamp_FR_R | SAV | plate 7 | 2 | 0.55 | 1842838 | 1842577 | 53980 | 15.58 | 0.0058 | 0.0006 | 20563 | 0.0028 | 1270 | 0.39 |
| 234 | Z67842_pamp_FR_R | SAV | plate 7 | 2 | 0.73 | 2431531 | 2431159 | 71154 | 12.37 | 0.0058 | 0.0006 | 24577 | 0.0024 | 1321 | 0.38 |
| 235 | Z67843_pamp_FR_R | SAV | plate 7 | 2 | 0.62 | 2066068 | 2065802 | 53081 | 30.59 | 0.0063 | 0.0007 | 17949 | 0.0027 | 1185 | 0.43 |
| 236 | Z67844_pamp_FR_R | SAV | plate 7 | 2 | 0.71 | 2363771 | 2363413 | 85790 | 12.83 | 0.0047 | 0.0006 | 23948 | 0.0025 | 1158 | 0.45 |
| 237 | Z67845_pamp_FR_R | SAV | plate 7 | 2 | 0.65 | 2182152 | 2181873 | 76385 | 18.78 | 0.0057 | 0.0005 | 22698 | 0.0027 | 1258 | 0.41 |
| 238 | Z67846_pamp_FR_U | PAR | plate 7 | 2 | 0.22 | 718583 | 718476 | 41523 | 11.79 | 0.0059 | 0.0006 | 19136 | 0.0028 | 1298 | 0.37 |
| 239 | Z67847_pamp_FR_U | PAR | plate 7 | 2 | 0.68 | 2261756 | 2261439 | 72355 | 14.39 | 0.0051 | 0.0005 | 23415 | 0.0027 | 1199 | 0.43 |
| 240 | Z67848_pamp_FR_U | PAR | plate 7 | 2 | 0.60 | 1992109 | 1991817 | 48447 | 20.15 | 0.0054 | 0.0006 | 20847 | 0.0027 | 1304 | 0.38 |
| 241 | Z67849_pamp_FR_U | PAR | plate 7 | 2 | 0.15 | 496028 | 495968 | 31952 | 11.34 | 0.0063 | 0.0007 | 15932 | 0.0028 | 1201 | 0.44 |
| 242 | Z67850_pamp_FR_U | PAR | plate 7 | 2 | 1.01 | 3358945 | 3358409 | 87915 | 22.17 | 0.0049 | 0.0005 | 25139 | 0.0026 | 1236 | 0.42 |
| 243 | Z67851_pamp_FR_U | PAR | plate 7 | 2 | 0.56 | 1852593 | 1852337 | 69287 | 12.28 | 0.0060 | 0.0006 | 22693 | 0.0026 | 1233 | 0.40 |
| 244 | Z67852_pamp_FR_U | DEC | plate 7 | 2 | 0.62 | 2078620 | 2078309 | 81089 | 14.35 | 0.0052 | 0.0005 | 23991 | 0.0027 | 1186 | 0.42 |
| 245 | Z67853_pamp_FR_U | DEC | plate 7 | 2 | 0.55 | 1843127 | 1842866 | 59155 | 19.72 | 0.0055 | 0.0006 | 21140 | 0.0029 | 1282 | 0.39 |

| | | | | | | | | | | | | | | | |
|---|---|---|---|---|---|---|---|---|---|---|---|---|---|---|---|
| 246 | Z67854_pamp_FR_U | DEC | plate 7 | 2 | 0.61 | 2028231 | 2027904 | 85781 | 12.61 | 0.0052 | 0.0005 | 23597 | 0.0027 | 1168 | 0.45 |
| 247 | Z67855_pamp_FR_U | DEC | plate 7 | 2 | 0.65 | 2159643 | 2159330 | 56355 | 10.85 | 0.0060 | 0.0007 | 20491 | 0.0029 | 1213 | 0.43 |
| 248 | Z67856_pamp_FR_U | DEC | plate 7 | 2 | 0.40 | 1332320 | 1332130 | 53920 | 15.09 | 0.0061 | 0.0006 | 21443 | 0.0029 | 1290 | 0.39 |
| 249 | Z67857_pamp_FR_U | DEC | plate 7 | 2 | 0.74 | 2481074 | 2480758 | 68242 | 20.14 | 0.0055 | 0.0005 | 23231 | 0.0028 | 1296 | 0.39 |
| 250 | Z67858_pamp_FR_U | DEC | plate 7 | 2 | 0.60 | 1999218 | 1998946 | 35598 | 9.85 | 0.0062 | 0.0006 | 14768 | 0.0026 | 1019 | 0.49 |
| 251 | Z67859_pamp_FR_U | DEC | plate 7 | 2 | 0.22 | 729271 | 729172 | 45465 | 10.95 | 0.0062 | 0.0007 | 18848 | 0.0028 | 1237 | 0.40 |
| 252 | Z67860_pamp_FR_U | DEC | plate 7 | 2 | 0.79 | 2644694 | 2644300 | 116129 | 8.90 | 0.0052 | 0.0006 | 25431 | 0.0023 | 962 | 0.54 |
| 253 | Z67861_pamp_FR_U | DEC | plate 7 | 2 | 0.39 | 1305382 | 1305194 | 54723 | 15.63 | 0.0059 | 0.0007 | 21382 | 0.0028 | 1300 | 0.36 |
| 254 | Z67862_pamp_FR_U | DEC | plate 7 | 2 | 0.54 | 1802717 | 1802457 | 54656 | 20.17 | 0.0059 | 0.0007 | 21158 | 0.0030 | 1282 | 0.39 |
| 255 | Z67863_pamp_FR_U | DEC | plate 7 | 2 | 0.61 | 2035961 | 2035588 | 98232 | 11.80 | 0.0057 | 0.0006 | 26249 | 0.0027 | 1283 | 0.40 |
| 256 | Z67864_pamp_FR_U | DEC | plate 7 | 2 | 0.34 | 1116995 | 1116846 | 60256 | 11.80 | 0.0058 | 0.0006 | 21978 | 0.0029 | 1300 | 0.39 |
| 257 | Z67865_pamp_FR_U | DEC | plate 7 | 2 | 0.51 | 1714607 | 1714362 | 69608 | 14.26 | 0.0052 | 0.0005 | 23314 | 0.0027 | 1253 | 0.42 |
| 258 | Z67866_pamp_FR_U | DEC | plate 7 | 2 | 0.54 | 1785143 | 1784874 | 57617 | 18.88 | 0.0067 | 0.0006 | 21262 | 0.0028 | 1285 | 0.39 |
| 259 | Z67867_pamp_FR_U | DEC | plate 7 | 2 | 0.44 | 1475100 | 1474929 | 40990 | 9.83 | 0.0066 | 0.0007 | 17451 | 0.0027 | 1200 | 0.42 |
| 260 | Z67875_pamp_FR_R | SAV | plate 5 | 2 | 1.09 | 3648359 | 3647805 | 47595 | 47.12 | 0.0064 | 0.0006 | 27226 | 0.0023 | 1313 | 0.39 |
| 261 | Z67876_pamp_FR_U | PET | plate 7 | 2 | 0.73 | 2443595 | 2443283 | 59383 | 21.59 | 0.0052 | 0.0006 | 22563 | 0.0028 | 1319 | 0.38 |
| 262 | Z67877_pamp_FR_U | PET | plate 7 | 2 | 0.42 | 1397296 | 1397104 | 54816 | 17.11 | 0.0050 | 0.0006 | 23709 | 0.0027 | 1313 | 0.38 |
| 263 | Z67878_pamp_FR_R | SAV | plate 5 | 2 | 0.93 | 3110274 | 3109811 | 56417 | 39.94 | 0.0062 | 0.0005 | 28963 | 0.0025 | 1333 | 0.38 |
| 264 | Z67879_pamp_FR_R | SAV | plate 5 | 2 | 0.82 | 2729536 | 2729167 | 36108 | 48.57 | 0.0078 | 0.0007 | 19690 | 0.0024 | 1173 | 0.45 |
| 265 | Z67880_pamp_FR_R | SAV | plate 5 | 2 | 0.90 | 3007422 | 3007028 | 61849 | 23.71 | 0.0064 | 0.0005 | 33748 | 0.0021 | 1355 | 0.36 |
| 266 | Z67881_pamp_FR_R | SAV | plate 5 | 2 | 0.61 | 2046683 | 2046445 | 51977 | 23.07 | 0.0056 | 0.0005 | 27564 | 0.0024 | 1287 | 0.38 |
| 267 | Z67882_pamp_FR_R | SAV | plate 5 | 2 | 0.84 | 2802436 | 2802054 | 56247 | 36.91 | 0.0057 | 0.0005 | 31709 | 0.0022 | 1360 | 0.35 |
| 268 | Z67883_pamp_FR_R | SAV | plate 5 | 2 | 0.59 | 1980588 | 1980303 | 59054 | 24.37 | 0.0059 | 0.0006 | 29623 | 0.0024 | 1327 | 0.39 |
| 269 | Z67884_pamp_FR_R | SAV | plate 5 | 2 | 0.89 | 2962226 | 2961795 | 71196 | 28.97 | 0.0065 | 0.0005 | 33014 | 0.0021 | 1294 | 0.40 |
| 270 | Z67885_pamp_FR_R | SAV | plate 5 | 2 | 1.02 | 3403205 | 3402714 | 62500 | 23.12 | 0.0054 | 0.0005 | 30631 | 0.0023 | 1190 | 0.44 |
| 271 | Z67886_pamp_FR_R | SAV | plate 5 | 2 | 0.10 | 328576 | 328521 | 23334 | 9.40 | 0.0052 | 0.0007 | 13438 | 0.0022 | 892 | 0.57 |
| 272 | Z67887_pamp_FR_U | PET | plate 7 | 2 | 0.75 | 2505764 | 2505403 | 72263 | 15.09 | 0.0055 | 0.0005 | 23663 | 0.0027 | 1274 | 0.39 |
| 273 | Z67917_pamp_FR_U | PAR | plate 5 | 2 | 1.07 | 3580533 | 3580056 | 63965 | 23.97 | 0.0095 | 0.0007 | 29509 | 0.0026 | 1367 | 0.36 |
| 274 | Z67918_pamp_FR_U | PAR | plate 5 | 2 | 0.51 | 1714346 | 1714099 | 45533 | 30.00 | 0.0061 | 0.0007 | 24105 | 0.0025 | 1269 | 0.40 |
| 275 | Z67919_pamp_FR_U | PAR | plate 5 | 2 | 1.09 | 3639475 | 3638921 | 33034 | 74.44 | 0.0066 | 0.0006 | 16101 | 0.0024 | 997 | 0.52 |
| 276 | Z67920_pamp_FR_U | PAR | plate 5 | 2 | 0.60 | 2007400 | 2007082 | 59633 | 21.58 | 0.0054 | 0.0006 | 31078 | 0.0022 | 1327 | 0.38 |
| 277 | Z67921_pamp_FR_U | PAR | plate 5 | 2 | 0.71 | 2352479 | 2352131 | 37137 | 20.28 | 0.0067 | 0.0007 | 21771 | 0.0024 | 1256 | 0.40 |

| | | | | | | | | | | | | | | | |
|---|---|---|---|---|---|---|---|---|---|---|---|---|---|---|---|
| 278 | Z67922_pamp_FR_U | PAR | plate 5 | 2 | 0.57 | 1915711 | 1915440 | 55043 | 21.93 | 0.0054 | 0.0006 | 31131 | 0.0023 | 1361 | 0.35 |
| 279 | Z67923_pamp_FR_U | PAR | plate 5 | 2 | 0.25 | 838657 | 838555 | 31571 | 19.23 | 0.0062 | 0.0007 | 19115 | 0.0024 | 1172 | 0.45 |
| 280 | Z67924_pamp_FR_U | PAR | plate 5 | 2 | 0.66 | 2203960 | 2203653 | 58428 | 24.22 | 0.0054 | 0.0006 | 32374 | 0.0021 | 1312 | 0.38 |
| 281 | Z67925_pamp_FR_U | PAR | plate 5 | 2 | 0.53 | 1769781 | 1769532 | 48865 | 21.26 | 0.0056 | 0.0005 | 28259 | 0.0022 | 1319 | 0.38 |
| 282 | Z67926_pamp_FR_U | PAR | plate 5 | 2 | 0.89 | 2963107 | 2962705 | 53978 | 30.92 | 0.0063 | 0.0006 | 29781 | 0.0023 | 1342 | 0.36 |
| 283 | Z67927_pamp_IT_U | ARR | plate 6 | 2 | 0.98 | 3280529 | 3280044 | 100690 | 19.52 | 0.0051 | 0.0005 | 32378 | 0.0023 | 951 | 0.52 |
| 284 | Z67928_pamp_IT_U | ARR | plate 6 | 2 | 0.47 | 1560184 | 1559909 | 83181 | 12.30 | 0.0059 | 0.0005 | 29055 | 0.0025 | 1029 | 0.48 |
| 285 | Z67929_pamp_IT_U | ARR | plate 6 | 2 | 0.59 | 1973092 | 1972851 | 78246 | 16.77 | 0.0064 | 0.0005 | 28329 | 0.0025 | 976 | 0.51 |
| 286 | Z67930_pamp_IT_U | ARR | plate 6 | 2 | 0.69 | 2311425 | 2311082 | 111177 | 13.01 | 0.0053 | 0.0005 | 33484 | 0.0022 | 933 | 0.53 |
| 287 | Z67931_pamp_IT_U | ARR | plate 6 | 2 | 0.39 | 1292813 | 1292638 | 72350 | 12.54 | 0.0053 | 0.0005 | 26868 | 0.0026 | 980 | 0.50 |
| 288 | Z67932_pamp_IT_U | ARR | plate 6 | 2 | 0.40 | 1342431 | 1342244 | 76228 | 9.97 | 0.0059 | 0.0005 | 28191 | 0.0026 | 957 | 0.52 |
| 289 | Z67933_pamp_IT_U | ARR | plate 6 | 2 | 0.71 | 2359516 | 2359165 | 104307 | 11.32 | 0.0058 | 0.0005 | 32835 | 0.0024 | 967 | 0.52 |
| 290 | Z67934_pamp_IT_U | ARR | plate 6 | 2 | 0.57 | 1913535 | 1913267 | 89594 | 13.08 | 0.0060 | 0.0005 | 29480 | 0.0024 | 934 | 0.53 |
| 291 | Z67935_pamp_IT_U | ARR | plate 6 | 2 | 0.25 | 847578 | 847468 | 51666 | 12.05 | 0.0061 | 0.0006 | 23153 | 0.0030 | 1018 | 0.48 |
| 292 | Z67936_pamp_IT_U | ARR | plate 6 | 2 | 0.20 | 676926 | 676822 | 54404 | 9.21 | 0.0062 | 0.0006 | 22644 | 0.0028 | 1019 | 0.48 |
| 293 | Z67937_pamp_IT_U | ARR | plate 6 | 2 | 0.35 | 1166779 | 1166628 | 71247 | 11.56 | 0.0057 | 0.0005 | 26230 | 0.0026 | 1014 | 0.48 |
| 294 | Z67938_pamp_IT_U | ARR | plate 6 | 2 | 0.60 | 1989416 | 1989148 | 83412 | 14.44 | 0.0047 | 0.0006 | 26006 | 0.0024 | 936 | 0.52 |
| 295 | Z67939_pamp_IT_R | MAN | plate 6 | 2 | 0.31 | 1035433 | 1035284 | 73703 | 8.67 | 0.0056 | 0.0006 | 24668 | 0.0026 | 935 | 0.53 |
| 296 | Z67940_pamp_IT_R | MAN | plate 6 | 2 | 0.68 | 2280277 | 2279950 | 107263 | 13.78 | 0.0052 | 0.0005 | 29938 | 0.0025 | 949 | 0.51 |
| 297 | Z67941_pamp_IT_R | MAN | plate 6 | 2 | 0.34 | 1146220 | 1146025 | 65716 | 11.98 | 0.0061 | 0.0006 | 24225 | 0.0029 | 1030 | 0.48 |
| 298 | Z67942_pamp_IT_R | MAN | plate 6 | 2 | 0.21 | 692757 | 692664 | 42774 | 11.96 | 0.0060 | 0.0006 | 20710 | 0.0031 | 1021 | 0.48 |
| 299 | Z67943_pamp_IT_R | MAN | plate 6 | 2 | 0.45 | 1487951 | 1487764 | 89517 | 11.08 | 0.0060 | 0.0005 | 27988 | 0.0027 | 1056 | 0.47 |
| 300 | Z67944_pamp_IT_R | MAN | plate 6 | 2 | 0.77 | 2562526 | 2562175 | 99160 | 15.53 | 0.0054 | 0.0004 | 29210 | 0.0024 | 945 | 0.53 |
| 301 | Z67945_pamp_IT_R | MAN | plate 6 | 2 | 0.39 | 1310369 | 1310200 | 82237 | 10.68 | 0.0055 | 0.0005 | 26446 | 0.0027 | 963 | 0.51 |
| 302 | Z67946_pamp_IT_R | MAN | plate 6 | 2 | 0.41 | 1361418 | 1361235 | 87008 | 10.57 | 0.0054 | 0.0005 | 27109 | 0.0026 | 974 | 0.51 |
| 303 | Z67947_pamp_IT_R | MAN | plate 6 | 2 | 0.65 | 2152955 | 2152647 | 104776 | 12.34 | 0.0050 | 0.0005 | 28996 | 0.0024 | 899 | 0.55 |
| 304 | Z67948_pamp_IT_R | MAN | plate 6 | 2 | 0.46 | 1549750 | 1549543 | 91114 | 11.14 | 0.0053 | 0.0005 | 26590 | 0.0027 | 969 | 0.52 |
| 305 | Z67949_pamp_IT_R | MAN | plate 6 | 2 | 0.62 | 2077294 | 2077027 | 99912 | 13.41 | 0.0055 | 0.0005 | 27597 | 0.0027 | 948 | 0.52 |
| 306 | Z67950_pamp_IT_R | MAN | plate 6 | 2 | 0.47 | 1563948 | 1563732 | 65546 | 16.25 | 0.0055 | 0.0005 | 24679 | 0.0030 | 1073 | 0.46 |
| 307 | Z67979_pamp_GR_U | GAL | plate 5 | 2 | 0.39 | 1309541 | 1309353 | 41589 | 15.05 | 0.0059 | 0.0006 | 23256 | 0.0024 | 1028 | 0.50 |
| 308 | Z67980_pamp_GR_U | GAL | plate 5 | 2 | 0.56 | 1865575 | 1865297 | 54377 | 24.30 | 0.0057 | 0.0006 | 28026 | 0.0022 | 1057 | 0.48 |
| 309 | Z67981_pamp_GR_U | GAL | plate 5 | 2 | 0.63 | 2107960 | 2107655 | 65107 | 19.96 | 0.0056 | 0.0005 | 31673 | 0.0023 | 1000 | 0.50 |

| | | | | | | | | | | | | | | | |
|---|---|---|---|---|---|---|---|---|---|---|---|---|---|---|---|
| 310 | Z67982_pamp_GR_U | GAL | plate 5 | 2 | 0.39 | 1292454 | 1292268 | 43054 | 19.46 | 0.0063 | 0.0006 | 22560 | 0.0026 | 1020 | 0.49 |
| 311 | Z67983_pamp_GR_U | GAL | plate 5 | 2 | 0.16 | 535749 | 535675 | 37570 | 8.13 | 0.0070 | 0.0008 | 18333 | 0.0025 | 975 | 0.52 |
| 312 | Z67984_pamp_GR_U | GAL | plate 5 | 2 | 0.50 | 1682438 | 1682197 | 59575 | 15.93 | 0.0054 | 0.0006 | 28301 | 0.0024 | 1059 | 0.47 |
| 313 | Z67985_pamp_GR_U | GAL | plate 5 | 2 | 0.15 | 484388 | 484310 | 33751 | 9.30 | 0.0062 | 0.0007 | 18108 | 0.0025 | 907 | 0.54 |
| 314 | Z67986_pamp_GR_U | GAL | plate 5 | 2 | 0.23 | 755427 | 755311 | 41092 | 10.73 | 0.0065 | 0.0007 | 21391 | 0.0023 | 968 | 0.51 |
| 315 | Z67987_pamp_GR_U | GAL | plate 5 | 2 | 3.76 | 12532722 | 12530898 | 37304 | 13.32 | 0.0064 | 0.0008 | 18632 | 0.0025 | 942 | 0.53 |
| 316 | Z67988_pamp_GR_U | GAL | plate 5 | 2 | 0.20 | 656015 | 655911 | 45082 | 9.54 | 0.0064 | 0.0008 | 22991 | 0.0024 | 1004 | 0.50 |
| 317 | Z67989_pamp_GR_R | KAT | plate 5 | 2 | 0.50 | 1681382 | 1681135 | 47086 | 10.33 | 0.0072 | 0.0007 | 22061 | 0.0025 | 966 | 0.52 |
| 318 | Z67990_pamp_GR_R | KAT | plate 5 | 2 | 0.34 | 1147767 | 1147601 | 44253 | 12.55 | 0.0066 | 0.0006 | 22569 | 0.0025 | 1026 | 0.49 |
| 319 | Z67991_pamp_GR_R | KAT | plate 5 | 2 | 0.47 | 1553452 | 1553218 | 30160 | 14.35 | 0.0049 | 0.0007 | 13610 | 0.0016 | 610 | 0.69 |
| 320 | Z67992_pamp_GR_R | KAT | plate 6 | 2 | 0.69 | 2310237 | 2309958 | 128277 | 10.91 | 0.0068 | 0.0006 | 44271 | 0.0020 | 1002 | 0.50 |
| 321 | Z67993_pamp_GR_R | KAT | plate 6 | 2 | 0.35 | 1178022 | 1177823 | 71446 | 7.70 | 0.0060 | 0.0006 | 25587 | 0.0025 | 972 | 0.50 |
| 322 | Z67994_pamp_GR_R | KAT | plate 6 | 2 | 0.51 | 1701046 | 1700821 | 77745 | 10.46 | 0.0060 | 0.0005 | 28290 | 0.0026 | 1018 | 0.50 |
| 323 | Z67995_pamp_GR_R | KAT | plate 6 | 2 | 0.48 | 1612567 | 1612333 | 85514 | 8.86 | 0.0059 | 0.0005 | 28868 | 0.0025 | 989 | 0.51 |
| 324 | Z67996_pamp_GR_R | KAT | plate 6 | 2 | 0.48 | 1598347 | 1598134 | 101211 | 8.06 | 0.0061 | 0.0006 | 31677 | 0.0024 | 984 | 0.51 |
| 325 | Z67997_pamp_GR_R | KAT | plate 6 | 2 | 0.22 | 748616 | 748504 | 51724 | 5.62 | 0.0064 | 0.0006 | 19542 | 0.0025 | 919 | 0.54 |
| 326 | Z67998_pamp_GR_R | KAT | plate 6 | 2 | 0.27 | 901382 | 901232 | 43006 | 6.78 | 0.0059 | 0.0007 | 18305 | 0.0024 | 892 | 0.54 |

**Table S5.** Details of DNA sequencing data for each *Ch. clathrata* individual. The sample identities (SampleID) are combinations of a unique code, species abbreviation ("clat"), country abbreviation (see Table S1 for a key for this and the population abbreviations), and a habitat type indicator ("R" for rural; "U" for urban). Sequencing batch indicates the sequencing platform (1 = HiSeq X PE150; 2 = NovaSeq X Plus PE150). Clusters refer to the number of clusters in initial grouping of similar sequences within an individual. Average depth refers to the average read depth. Hetero estimate refers to the estimated per-site heterozygosity within individual samples. Error estimate is the estimated per-site sequencing error rate, inferred from base call mismatches among reads within a cluster. Reads consensus refers to the number of consensus loci (assembled clusters) successfully built after clustering and filtering. Heterozygosity is the average heterozygosity across loci. Loci in assembly is the total number of RAD loci retained in the final assembly after applying all filters. Missing data is the proportion of missing data.

| No. | SampleID | Population | Sequencing plate | Sequencing batch | Gb sequenced | Total reads | Reads passed filter | Clusters | Average depth | Hetero estimate | Error estimate | Reads consens | Heterozygosity | Loci in assembly | Missing data |
|---|---|---|---|---|---|---|---|---|---|---|---|---|---|---|---|
| 1 | Z30529_clat_CZ | BIS | plate 3 | 1 | 0.08 | 266663 | 266662 | 15103 | 10.30 | 0.0056 | 0.0007 | 5497 | 0.0014 | 208 | 0.73 |
| 2 | Z30531_clat_CZ | BIS | plate 3 | 1 | 0.99 | 3288367 | 3288348 | 29453 | 87.37 | 0.0054 | 0.0007 | 12973 | 0.0021 | 379 | 0.51 |
| 3 | Z30532_clat_CZ | BIS | plate 3 | 1 | 1.76 | 5876742 | 5876717 | 39583 | 41.80 | 0.0057 | 0.0007 | 17237 | 0.0025 | 459 | 0.41 |
| 4 | Z30533_clat_CZ | BIS | plate 3 | 1 | 0.41 | 1356926 | 1356920 | 13139 | 34.81 | 0.0055 | 0.0004 | 4070 | 0.0011 | 149 | 0.81 |
| 5 | Z30534_clat_CZ | BIS | plate 3 | 1 | 0.34 | 1145574 | 1145566 | 27854 | 22.49 | 0.0061 | 0.0006 | 12195 | 0.0025 | 374 | 0.51 |
| 6 | Z30535_clat_CZ | BIS | plate 3 | 1 | 0.70 | 2347745 | 2347731 | 13357 | 155.75 | 0.0060 | 0.0006 | 3476 | 0.0013 | 154 | 0.81 |
| 7 | Z30537_clat_CZ | BIS | plate 3 | 1 | 0.56 | 1854337 | 1854327 | 47428 | 26.43 | 0.0066 | 0.0007 | 20146 | 0.0023 | 454 | 0.43 |
| 8 | Z30538_clat_CZ | BIS | plate 3 | 1 | 0.66 | 2197011 | 2196999 | 48097 | 27.13 | 0.0078 | 0.0009 | 21344 | 0.0028 | 518 | 0.39 |
| 9 | Z30539_clat_CZ | BIS | plate 3 | 1 | 0.68 | 2277485 | 2277473 | 66117 | 22.66 | 0.0050 | 0.0006 | 24866 | 0.0025 | 506 | 0.34 |
| 10 | Z30540_clat_CZ | BIS | plate 3 | 1 | 0.56 | 1868880 | 1868871 | 43479 | 23.58 | 0.0054 | 0.0006 | 18804 | 0.0025 | 479 | 0.36 |
| 11 | Z30541_clat_CZ | BIS | plate 3 | 1 | 0.13 | 425574 | 425571 | 15156 | 12.44 | 0.0052 | 0.0007 | 5344 | 0.0013 | 205 | 0.73 |
| 12 | Z30632_clat_FI_U | HEI | plate 2 | 1 | 0.50 | 1681585 | 1681580 | 37155 | 14.03 | 0.0052 | 0.0007 | 17469 | 0.0032 | 616 | 0.26 |
| 13 | Z36558_clat_CZ_U | POH | plate 2 | 1 | 0.97 | 3248273 | 3248265 | 46574 | 25.27 | 0.0049 | 0.0007 | 20659 | 0.0030 | 585 | 0.28 |
| 14 | Z36640_clat_SE_U | BRO | plate 2 | 1 | 0.51 | 1688423 | 1688410 | 43930 | 18.25 | 0.0054 | 0.0006 | 19875 | 0.0030 | 588 | 0.28 |
| 15 | Z36641_clat_SE_U | BRO | plate 2 | 1 | 0.76 | 2518009 | 2518003 | 51368 | 18.86 | 0.0048 | 0.0006 | 21306 | 0.0028 | 566 | 0.30 |
| 16 | Z36642_clat_SE_U | BRO | plate 2 | 1 | 0.53 | 1754815 | 1754806 | 49636 | 20.77 | 0.0048 | 0.0007 | 20786 | 0.0029 | 596 | 0.27 |
| 17 | Z36643_clat_SE_U | BRO | plate 2 | 1 | 0.63 | 2091970 | 2091962 | 54546 | 19.95 | 0.0046 | 0.0006 | 22838 | 0.0026 | 572 | 0.29 |
| 18 | Z36644_clat_SE_U | BRO | plate 2 | 1 | 0.49 | 1645344 | 1645339 | 41100 | 16.82 | 0.0050 | 0.0006 | 19826 | 0.0030 | 582 | 0.28 |
| 19 | Z36645_clat_SE_R | TIS | plate 2 | 1 | 0.64 | 2119197 | 2119189 | 58109 | 23.02 | 0.0048 | 0.0006 | 23912 | 0.0026 | 569 | 0.31 |
| 20 | Z36646_clat_SE_R | TIS | plate 3 | 1 | 0.61 | 2046139 | 2046135 | 60616 | 24.21 | 0.0081 | 0.0005 | 31834 | 0.0031 | 298 | 0.68 |
| 21 | Z36647_clat_SE_R | TIS | plate 3 | 1 | 0.66 | 2186547 | 2186545 | 49521 | 34.87 | 0.0073 | 0.0005 | 27694 | 0.0034 | 262 | 0.71 |

| | | | | | | | | | | | | | | | |
|---|---|---|---|---|---|---|---|---|---|---|---|---|---|---|---|
| 22 | Z36648_clat_SE_R | SON | plate 2 | 1 | 0.55 | 1823661 | 1823656 | 49423 | 18.62 | 0.0050 | 0.0006 | 21696 | 0.0028 | 576 | 0.30 |
| 23 | Z36649_clat_SE_R | KRI | plate 2 | 1 | 0.66 | 2190568 | 2190563 | 52553 | 16.88 | 0.0052 | 0.0006 | 22255 | 0.0027 | 556 | 0.32 |
| 24 | Z36650_clat_SE_R | KRI | plate 2 | 1 | 0.66 | 2196134 | 2196123 | 57314 | 18.24 | 0.0050 | 0.0006 | 23268 | 0.0026 | 563 | 0.29 |
| 25 | Z36741_clat_FI_U | MAL | plate 2 | 1 | 0.77 | 2579335 | 2579331 | 62984 | 23.49 | 0.0046 | 0.0006 | 22645 | 0.0026 | 538 | 0.32 |
| 26 | Z36742_clat_FI_U | MAL | plate 2 | 1 | 0.52 | 1743335 | 1743329 | 48865 | 22.73 | 0.0049 | 0.0006 | 21073 | 0.0029 | 577 | 0.29 |
| 27 | Z36743_clat_FI_U | MAL | plate 2 | 1 | 1.19 | 3952268 | 3952244 | 40220 | 33.25 | 0.0050 | 0.0007 | 19444 | 0.0029 | 603 | 0.24 |
| 28 | Z36744_clat_FI_U | MAL | plate 2 | 1 | 0.43 | 1422571 | 1422567 | 45043 | 22.37 | 0.0049 | 0.0006 | 20245 | 0.0029 | 570 | 0.27 |
| 29 | Z36745_clat_FI_U | MAL | plate 2 | 1 | 0.40 | 1337698 | 1337693 | 38939 | 18.81 | 0.0046 | 0.0006 | 18918 | 0.0029 | 594 | 0.26 |
| 30 | Z36746_clat_FI_U | MAL | plate 2 | 1 | 0.91 | 3045168 | 3045158 | 52036 | 40.43 | 0.0045 | 0.0006 | 22457 | 0.0026 | 555 | 0.30 |
| 31 | Z36747_clat_FI_U | MAL | plate 2 | 1 | 0.53 | 1776156 | 1776146 | 39988 | 29.42 | 0.0046 | 0.0007 | 19986 | 0.0027 | 591 | 0.26 |
| 32 | Z36748_clat_FI_U | MAL | plate 2 | 1 | 1.07 | 3551855 | 3551840 | 57213 | 30.73 | 0.0046 | 0.0006 | 23213 | 0.0027 | 556 | 0.28 |
| 33 | Z36817_clat_CZ_U | POH | plate 2 | 1 | 0.53 | 1777734 | 1777727 | 57957 | 21.29 | 0.0045 | 0.0007 | 22121 | 0.0027 | 528 | 0.35 |
| 34 | Z36822_clat_CZ_U | DIV | plate 1 | 1 | 0.41 | 1370756 | 1370748 | 33281 | 23.62 | 0.0050 | 0.0007 | 16305 | 0.0031 | 525 | 0.29 |
| 35 | Z36823_clat_CZ_U | DIV | plate 1 | 1 | 0.64 | 2142599 | 2142589 | 69631 | 16.78 | 0.0045 | 0.0006 | 21834 | 0.0024 | 517 | 0.37 |
| 36 | Z36824_clat_CZ_U | DIV | plate 1 | 1 | 0.57 | 1883482 | 1883477 | 62783 | 17.09 | 0.0049 | 0.0006 | 20730 | 0.0025 | 534 | 0.36 |
| 37 | Z36825_clat_CZ_U | DIV | plate 1 | 1 | 0.75 | 2495499 | 2495489 | 76300 | 18.24 | 0.0045 | 0.0006 | 22384 | 0.0024 | 454 | 0.42 |
| 38 | Z36826_clat_CZ_U | DIV | plate 1 | 1 | 0.43 | 1428885 | 1428865 | 55712 | 14.43 | 0.0046 | 0.0006 | 19558 | 0.0028 | 506 | 0.37 |
| 39 | Z36827_clat_CZ_U | DIV | plate 1 | 1 | 0.56 | 1865464 | 1865452 | 59175 | 14.83 | 0.0050 | 0.0006 | 20563 | 0.0027 | 548 | 0.30 |
| 40 | Z36828_clat_CZ_U | DIV | plate 1 | 1 | 1.45 | 4829285 | 4829264 | 63171 | 63.66 | 0.0048 | 0.0007 | 22285 | 0.0026 | 524 | 0.33 |
| 41 | Z36829_clat_CZ_U | DIV | plate 1 | 1 | 1.12 | 3747432 | 3747419 | 90145 | 26.82 | 0.0045 | 0.0006 | 28811 | 0.0019 | 475 | 0.40 |
| 42 | Z36830_clat_CZ_U | DIV | plate 1 | 1 | 0.99 | 3288943 | 3288927 | 100800 | 17.15 | 0.0043 | 0.0006 | 29528 | 0.0018 | 447 | 0.44 |
| 43 | Z36831_clat_CZ_U | DIV | plate 1 | 1 | 0.75 | 2502692 | 2502684 | 77301 | 21.57 | 0.0044 | 0.0006 | 26663 | 0.0021 | 477 | 0.39 |
| 44 | Z36832_clat_CZ_U | DIV | plate 1 | 1 | 1.31 | 4375717 | 4375684 | 84018 | 34.14 | 0.0042 | 0.0006 | 25990 | 0.0019 | 452 | 0.41 |
| 45 | Z36836_clat_CZ_R | MIL | plate 2 | 1 | 0.48 | 1605769 | 1605763 | 53403 | 23.64 | 0.0051 | 0.0007 | 20937 | 0.0030 | 562 | 0.31 |
| 46 | Z36837_clat_CZ_R | MIL | plate 2 | 1 | 0.47 | 1559709 | 1559702 | 52281 | 24.90 | 0.0052 | 0.0006 | 21282 | 0.0032 | 565 | 0.30 |
| 47 | Z36838_clat_CZ_R | MIL | plate 2 | 1 | 0.37 | 1228515 | 1228504 | 47194 | 21.52 | 0.0053 | 0.0006 | 20329 | 0.0032 | 578 | 0.29 |
| 48 | Z36839_clat_CZ_R | MIL | plate 2 | 1 | 0.43 | 1426217 | 1426207 | 40747 | 29.02 | 0.0052 | 0.0007 | 19363 | 0.0033 | 549 | 0.29 |
| 49 | Z36840_clat_CZ_R | MIL | plate 2 | 1 | 0.65 | 2166095 | 2166084 | 46808 | 34.35 | 0.0050 | 0.0006 | 20368 | 0.0032 | 537 | 0.33 |
| 50 | Z37527_clat_CZ | CHL | plate 1 | 1 | 1.27 | 4223455 | 4223431 | 75928 | 31.93 | 0.0050 | 0.0007 | 22015 | 0.0025 | 481 | 0.37 |
| 51 | Z37528_clat_CZ | CHL | plate 1 | 1 | 0.84 | 2815755 | 2815735 | 129956 | 12.11 | 0.0052 | 0.0006 | 34657 | 0.0018 | 423 | 0.44 |
| 52 | Z37529_clat_CZ | CHL | plate 1 | 1 | 0.87 | 2892201 | 2892184 | 46590 | 14.81 | 0.0054 | 0.0007 | 18269 | 0.0028 | 512 | 0.31 |
| 53 | Z37530_clat_CZ | CHL | plate 1 | 1 | 0.68 | 2256494 | 2256483 | 66795 | 19.04 | 0.0052 | 0.0007 | 22155 | 0.0026 | 518 | 0.32 |

| | | | | | | | | | | | | | | | |
|---|---|---|---|---|---|---|---|---|---|---|---|---|---|---|---|
| 54 | Z37531_clat_CZ | CHL | plate 1 | 1 | 0.94 | 3135174 | 3135155 | 37420 | 58.39 | 0.0058 | 0.0008 | 14755 | 0.0028 | 467 | 0.38 |
| 55 | Z37532_clat_CZ | CHL | plate 1 | 1 | 0.78 | 2616298 | 2616291 | 65092 | 20.70 | 0.0052 | 0.0006 | 21832 | 0.0026 | 508 | 0.35 |
| 56 | Z37533_clat_CZ | CHL | plate 1 | 1 | 0.55 | 1844524 | 1844514 | 42752 | 33.54 | 0.0056 | 0.0007 | 16312 | 0.0026 | 473 | 0.38 |
| 57 | Z37534_clat_CZ | CHL | plate 1 | 1 | 1.49 | 4979428 | 4979407 | 24361 | 127.62 | 0.0083 | 0.0008 | 8817 | 0.0023 | 290 | 0.60 |
| 58 | Z37535_clat_CZ | CHL | plate 1 | 1 | 0.67 | 2241073 | 2241059 | 51701 | 24.78 | 0.0033 | 0.0006 | 24613 | 0.0019 | 508 | 0.33 |
| 59 | Z37538_clat_CZ | CHL | plate 1 | 1 | 0.94 | 3122796 | 3122787 | 29513 | 72.96 | 0.0055 | 0.0008 | 13848 | 0.0029 | 459 | 0.38 |
| 60 | Z37539_clat_CZ | CHL | plate 1 | 1 | 0.31 | 1046306 | 1046299 | 45476 | 13.34 | 0.0052 | 0.0007 | 17905 | 0.0023 | 437 | 0.41 |
| 61 | Z37540_clat_CZ | CHL | plate 1 | 1 | 0.44 | 1463568 | 1463520 | 43083 | 19.31 | 0.0048 | 0.0008 | 17225 | 0.0021 | 455 | 0.40 |
| 62 | Z37541_clat_CZ | CHL | plate 1 | 1 | 1.15 | 3816847 | 3816820 | 113692 | 11.59 | 0.0059 | 0.0007 | 31492 | 0.0021 | 448 | 0.40 |
| 63 | Z37542_clat_CZ | CHL | plate 1 | 1 | 0.56 | 1857213 | 1857200 | 28532 | 24.07 | 0.0066 | 0.0008 | 12228 | 0.0026 | 390 | 0.48 |
| 64 | Z37543_clat_CZ | CHL | plate 1 | 1 | 0.25 | 822249 | 822245 | 53718 | 11.47 | 0.0055 | 0.0007 | 20160 | 0.0026 | 506 | 0.32 |
| 65 | Z37544_clat_CZ | CHL | plate 1 | 1 | 0.45 | 1485759 | 1485747 | 62709 | 13.96 | 0.0049 | 0.0007 | 22819 | 0.0023 | 479 | 0.36 |
| 66 | Z38322_clat_BE_R | NIS | plate 1 | 1 | 0.47 | 1564799 | 1564792 | 59886 | 14.80 | 0.0044 | 0.0007 | 19660 | 0.0027 | 506 | 0.36 |
| 67 | Z38323_clat_BE_R | NIS | plate 1 | 1 | 0.53 | 1779652 | 1779647 | 68224 | 18.16 | 0.0048 | 0.0006 | 20994 | 0.0026 | 516 | 0.36 |
| 68 | Z38324_clat_BE_R | NIS | plate 1 | 1 | 0.64 | 2131495 | 2131486 | 82572 | 16.06 | 0.0047 | 0.0006 | 22043 | 0.0024 | 474 | 0.41 |
| 69 | Z38325_clat_BE_R | NIS | plate 1 | 1 | 0.67 | 2222365 | 2222360 | 76914 | 15.07 | 0.0046 | 0.0006 | 21571 | 0.0025 | 500 | 0.37 |
| 70 | Z38326_clat_BE_R | NIS | plate 1 | 1 | 1.16 | 3859266 | 3859250 | 177770 | 8.24 | 0.0055 | 0.0006 | 29270 | 0.0017 | 369 | 0.51 |
| 71 | Z38327_clat_BE_R | NIS | plate 1 | 1 | 0.92 | 3051611 | 3051596 | 82715 | 24.04 | 0.0043 | 0.0006 | 21963 | 0.0024 | 489 | 0.42 |
| 72 | Z38328_clat_BE_R | NIS | plate 1 | 1 | 0.83 | 2773015 | 2773001 | 74300 | 17.08 | 0.0047 | 0.0006 | 21970 | 0.0024 | 537 | 0.33 |
| 73 | Z38329_clat_BE_R | NIS | plate 1 | 1 | 0.34 | 1122553 | 1122547 | 51683 | 12.28 | 0.0051 | 0.0006 | 18941 | 0.0028 | 528 | 0.33 |
| 74 | Z38332_clat_BE_R | NIS | plate 1 | 1 | 0.84 | 2808173 | 2808161 | 80295 | 15.43 | 0.0043 | 0.0007 | 20178 | 0.0024 | 464 | 0.43 |
| 75 | Z38334_clat_BE_R | NIS | plate 1 | 1 | 0.09 | 305242 | 305239 | 23676 | 7.96 | 0.0052 | 0.0008 | 12356 | 0.0031 | 467 | 0.37 |
| 76 | Z38335_clat_BE_R | NIS | plate 1 | 1 | 0.59 | 1972776 | 1972767 | 83770 | 16.11 | 0.0054 | 0.0007 | 24096 | 0.0022 | 449 | 0.43 |
| 77 | Z38336_clat_BE_R | NIS | plate 1 | 1 | 0.41 | 1369261 | 1369257 | 65056 | 13.03 | 0.0044 | 0.0007 | 18654 | 0.0026 | 509 | 0.36 |
| 78 | Z38337_clat_BE_R | NIS | plate 1 | 1 | 1.37 | 4568334 | 4568305 | 113393 | 26.23 | 0.0043 | 0.0007 | 23651 | 0.0021 | 411 | 0.47 |
| 79 | Z38338_clat_BE_R | NIS | plate 1 | 1 | 0.75 | 2486174 | 2486159 | 72060 | 17.72 | 0.0044 | 0.0007 | 19512 | 0.0024 | 470 | 0.41 |
| 80 | Z38339_clat_BE_R | NIS | plate 1 | 1 | 0.63 | 2107970 | 2107954 | 90620 | 13.57 | 0.0048 | 0.0007 | 20888 | 0.0023 | 435 | 0.47 |
| 81 | Z38346_clat_BE_U | JOS | plate 1 | 1 | 0.75 | 2515776 | 2515765 | 72448 | 14.45 | 0.0047 | 0.0006 | 18982 | 0.0027 | 483 | 0.37 |
| 82 | Z38347_clat_BE_U | JOS | plate 1 | 1 | 0.62 | 2070471 | 2070454 | 70957 | 15.39 | 0.0043 | 0.0006 | 19451 | 0.0025 | 480 | 0.39 |
| 83 | Z38348_clat_BE_U | JOS | plate 1 | 1 | 0.69 | 2301123 | 2301109 | 95107 | 12.04 | 0.0048 | 0.0007 | 19134 | 0.0025 | 440 | 0.47 |
| 84 | Z38349_clat_BE_U | JOS | plate 1 | 1 | 0.79 | 2625955 | 2625935 | 96285 | 15.35 | 0.0046 | 0.0006 | 20185 | 0.0024 | 413 | 0.47 |
| 85 | Z38350_clat_BE_U | JOS | plate 1 | 1 | 0.96 | 3204297 | 3204270 | 120706 | 12.15 | 0.0045 | 0.0007 | 20326 | 0.0023 | 395 | 0.50 |

| | | | | | | | | | | | | | | | |
|---|---|---|---|---|---|---|---|---|---|---|---|---|---|---|---|
| 86 | Z38351_clat_BE_U | JOS | plate 1 | 1 | 0.47 | 1556583 | 1556576 | 59736 | 17.37 | 0.0046 | 0.0007 | 18728 | 0.0028 | 487 | 0.35 |
| 87 | Z38354_clat_BE_U | MOE | plate 3 | 1 | 0.60 | 2003007 | 2002998 | 54692 | 15.14 | 0.0049 | 0.0006 | 21432 | 0.0027 | 525 | 0.36 |
| 88 | Z38355_clat_BE_U | JOS | plate 1 | 1 | 0.55 | 1822097 | 1822090 | 86605 | 13.40 | 0.0047 | 0.0006 | 20428 | 0.0025 | 462 | 0.41 |
| 89 | Z38356_clat_BE_U | JOS | plate 1 | 1 | 0.59 | 1950974 | 1950967 | 74458 | 16.00 | 0.0043 | 0.0006 | 20095 | 0.0025 | 473 | 0.41 |
| 90 | Z38357_clat_BE_U | JOS | plate 1 | 1 | 0.74 | 2471280 | 2471268 | 84940 | 12.77 | 0.0048 | 0.0006 | 20920 | 0.0025 | 473 | 0.43 |
| 91 | Z38358_clat_BE_U | JOS | plate 1 | 1 | 0.79 | 2628098 | 2628092 | 134365 | 9.02 | 0.0050 | 0.0007 | 23574 | 0.0021 | 393 | 0.47 |
| 92 | Z38359_clat_BE_U | JOS | plate 1 | 1 | 0.83 | 2777193 | 2777175 | 82518 | 12.56 | 0.0048 | 0.0007 | 22306 | 0.0023 | 517 | 0.32 |
| 93 | Z38360_clat_BE_U | JOS | plate 1 | 1 | 0.61 | 2025841 | 2025831 | 74850 | 16.77 | 0.0040 | 0.0007 | 17272 | 0.0023 | 447 | 0.44 |
| 94 | Z38361_clat_BE_U | JOS | plate 1 | 1 | 0.50 | 1680617 | 1680609 | 60094 | 19.28 | 0.0063 | 0.0006 | 18567 | 0.0029 | 480 | 0.38 |
| 95 | Z38362_clat_BE_U | JOS | plate 1 | 1 | 0.50 | 1655854 | 1655838 | 76911 | 12.23 | 0.0044 | 0.0007 | 18800 | 0.0025 | 497 | 0.39 |
| 96 | Z38381_clat_BE_U | JOS | plate 1 | 1 | 0.78 | 2600677 | 2600663 | 90533 | 18.87 | 0.0063 | 0.0007 | 20994 | 0.0023 | 435 | 0.46 |
| 97 | Z38382_clat_BE_U | JOS | plate 1 | 1 | 0.78 | 2598166 | 2598152 | 75352 | 16.92 | 0.0045 | 0.0007 | 20860 | 0.0025 | 493 | 0.38 |
| 98 | Z38416_clat_BE_U | BUD | plate 1 | 1 | 1.03 | 3435176 | 3435156 | 135200 | 11.78 | 0.0043 | 0.0006 | 23407 | 0.0020 | 390 | 0.52 |
| 99 | Z38417_clat_BE_U | BUD | plate 1 | 1 | 0.88 | 2939151 | 2939139 | 53024 | 23.87 | 0.0046 | 0.0007 | 18554 | 0.0028 | 564 | 0.29 |
| 100 | Z38418_clat_BE_U | BUD | plate 1 | 1 | 0.61 | 2026832 | 2026820 | 55450 | 14.33 | 0.0047 | 0.0007 | 18155 | 0.0029 | 559 | 0.33 |
| 101 | Z38419_clat_BE_U | BUD | plate 1 | 1 | 0.69 | 2300946 | 2300929 | 68244 | 20.15 | 0.0042 | 0.0007 | 20410 | 0.0024 | 489 | 0.39 |
| 102 | Z38420_clat_BE_U | BUD | plate 1 | 1 | 0.54 | 1808036 | 1808025 | 61330 | 9.69 | 0.0058 | 0.0007 | 17358 | 0.0028 | 516 | 0.34 |
| 103 | Z38421_clat_BE_U | BUD | plate 1 | 1 | 0.71 | 2366362 | 2366344 | 74961 | 14.77 | 0.0044 | 0.0007 | 20397 | 0.0025 | 471 | 0.41 |
| 104 | Z38422_clat_BE_U | BUD | plate 1 | 1 | 0.71 | 2362374 | 2362368 | 78695 | 12.96 | 0.0048 | 0.0007 | 21442 | 0.0023 | 456 | 0.40 |
| 105 | Z38423_clat_BE_U | BUD | plate 1 | 1 | 0.15 | 494597 | 494596 | 36812 | 5.07 | 0.0056 | 0.0007 | 12603 | 0.0028 | 489 | 0.38 |
| 106 | Z38424_clat_BE_U | MOE | plate 3 | 1 | 0.53 | 1779064 | 1779057 | 37466 | 15.61 | 0.0050 | 0.0006 | 18204 | 0.0031 | 547 | 0.34 |
| 107 | Z38425_clat_BE_U | MOE | plate 3 | 1 | 0.65 | 2162309 | 2162301 | 58412 | 18.34 | 0.0048 | 0.0006 | 21830 | 0.0028 | 538 | 0.37 |
| 108 | Z38426_clat_BE_U | MOE | plate 3 | 1 | 0.51 | 1689559 | 1689549 | 46410 | 16.45 | 0.0048 | 0.0006 | 20005 | 0.0028 | 536 | 0.35 |
| 109 | Z38427_clat_BE_U | MOE | plate 3 | 1 | 0.52 | 1733029 | 1733028 | 45026 | 32.00 | 0.0047 | 0.0006 | 19523 | 0.0029 | 562 | 0.32 |
| 110 | Z38428_clat_BE_U | MOE | plate 3 | 1 | 0.58 | 1943796 | 1943789 | 53869 | 25.52 | 0.0050 | 0.0006 | 20884 | 0.0028 | 546 | 0.34 |
| 111 | Z38434_clat_SE_U | BRO | plate 2 | 1 | 0.67 | 2232704 | 2232701 | 56766 | 17.34 | 0.0048 | 0.0006 | 22663 | 0.0025 | 531 | 0.33 |
| 112 | Z38435_clat_SE_U | BRO | plate 2 | 1 | 0.47 | 1582880 | 1582873 | 39515 | 20.42 | 0.0051 | 0.0006 | 19518 | 0.0030 | 605 | 0.24 |
| 113 | Z38436_clat_SE_U | BRO | plate 2 | 1 | 0.77 | 2552629 | 2552627 | 54022 | 21.41 | 0.0050 | 0.0006 | 23286 | 0.0027 | 604 | 0.28 |
| 114 | Z38437_clat_SE_U | BRO | plate 2 | 1 | 0.55 | 1836406 | 1836398 | 45445 | 20.99 | 0.0047 | 0.0006 | 20406 | 0.0028 | 589 | 0.29 |
| 115 | Z38438_clat_SE_U | BRO | plate 2 | 1 | 1.41 | 4690038 | 4690004 | 51530 | 76.89 | 0.0047 | 0.0006 | 20378 | 0.0028 | 569 | 0.31 |
| 116 | Z38439_clat_SE_U | BRO | plate 2 | 1 | 0.61 | 2048782 | 2048779 | 55032 | 15.86 | 0.0048 | 0.0006 | 20898 | 0.0027 | 542 | 0.31 |
| 117 | Z38440_clat_SE_U | BRO | plate 2 | 1 | 0.53 | 1758677 | 1758668 | 58990 | 15.97 | 0.0049 | 0.0006 | 22489 | 0.0026 | 572 | 0.29 |

| | | | | | | | | | | | | | | | |
|---|---|---|---|---|---|---|---|---|---|---|---|---|---|---|---|
| 118 | Z38441_clat_SE_U | BRO | plate 2 | 1 | 0.64 | 2142613 | 2142607 | 48689 | 21.37 | 0.0049 | 0.0006 | 21087 | 0.0028 | 575 | 0.30 |
| 119 | Z38442_clat_SE_U | BRO | plate 2 | 1 | 0.28 | 937497 | 937491 | 37504 | 11.66 | 0.0051 | 0.0007 | 17480 | 0.0030 | 611 | 0.26 |
| 120 | Z38443_clat_SE_U | BRO | plate 2 | 1 | 0.58 | 1935500 | 1935491 | 48335 | 18.12 | 0.0045 | 0.0006 | 19976 | 0.0028 | 564 | 0.30 |
| 121 | Z38444_clat_SE_U | BRO | plate 2 | 1 | 0.37 | 1247992 | 1247985 | 37793 | 15.99 | 0.0051 | 0.0006 | 18062 | 0.0030 | 601 | 0.27 |
| 122 | Z38450_clat_SE_R | SON | plate 2 | 1 | 0.56 | 1870805 | 1870800 | 46260 | 22.74 | 0.0048 | 0.0006 | 21562 | 0.0028 | 589 | 0.28 |
| 123 | Z38451_clat_CZ_U | DIV | plate 1 | 1 | 1.00 | 3336630 | 3336616 | 89790 | 14.79 | 0.0063 | 0.0006 | 23234 | 0.0023 | 447 | 0.43 |
| 124 | Z38452_clat_CZ_U | DIV | plate 1 | 1 | 0.97 | 3244966 | 3244948 | 68145 | 26.46 | 0.0045 | 0.0006 | 20840 | 0.0026 | 485 | 0.38 |
| 125 | Z38453_clat_CZ_U | DIV | plate 1 | 1 | 1.02 | 3411870 | 3411859 | 78964 | 32.75 | 0.0045 | 0.0006 | 23171 | 0.0025 | 536 | 0.34 |
| 126 | Z38454_clat_CZ_R | MIL | plate 1 | 1 | 0.58 | 1932102 | 1932097 | 74806 | 14.09 | 0.0044 | 0.0006 | 21485 | 0.0023 | 492 | 0.38 |
| 127 | Z38455_clat_CZ_R | MIL | plate 1 | 1 | 0.58 | 1938624 | 1938614 | 56706 | 26.31 | 0.0043 | 0.0006 | 19416 | 0.0026 | 539 | 0.35 |
| 128 | Z38456_clat_CZ_R | MIL | plate 1 | 1 | 0.69 | 2298092 | 2298082 | 89785 | 16.03 | 0.0047 | 0.0006 | 22497 | 0.0022 | 451 | 0.42 |
| 129 | Z38457_clat_CZ_R | MIL | plate 1 | 1 | 0.82 | 2725405 | 2725388 | 67980 | 21.50 | 0.0043 | 0.0006 | 20797 | 0.0024 | 502 | 0.35 |
| 130 | Z38458_clat_CZ_R | MIL | plate 1 | 1 | 0.76 | 2544908 | 2544894 | 80172 | 23.92 | 0.0042 | 0.0006 | 22102 | 0.0023 | 498 | 0.36 |
| 131 | Z38459_clat_CZ_R | MIL | plate 1 | 1 | 0.56 | 1876884 | 1876876 | 67988 | 15.25 | 0.0047 | 0.0006 | 22107 | 0.0024 | 516 | 0.36 |
| 132 | Z38460_clat_CZ_R | MIL | plate 1 | 1 | 0.50 | 1658861 | 1658857 | 55506 | 21.55 | 0.0047 | 0.0006 | 19622 | 0.0027 | 526 | 0.32 |
| 133 | Z38461_clat_CZ_R | MIL | plate 1 | 1 | 0.76 | 2546726 | 2546710 | 96410 | 15.55 | 0.0048 | 0.0006 | 23713 | 0.0021 | 427 | 0.46 |
| 134 | Z38476_clat_CZ_U | POH | plate 2 | 1 | 0.56 | 1870135 | 1870135 | 31318 | 10.95 | 0.0052 | 0.0008 | 15933 | 0.0026 | 484 | 0.43 |
| 135 | Z38477_clat_CZ_U | POH | plate 2 | 1 | 0.65 | 2175473 | 2175465 | 47176 | 29.25 | 0.0049 | 0.0006 | 20943 | 0.0029 | 565 | 0.31 |
| 136 | Z38478_clat_CZ_U | POH | plate 2 | 1 | 0.34 | 1146006 | 1145999 | 36583 | 15.22 | 0.0050 | 0.0007 | 17831 | 0.0030 | 579 | 0.28 |
| 137 | Z38479_clat_CZ_U | DIV | plate 1 | 1 | 1.32 | 4406497 | 4406462 | 105280 | 29.60 | 0.0044 | 0.0006 | 23586 | 0.0021 | 425 | 0.46 |
| 138 | Z38480_clat_CZ_U | DIV | plate 1 | 1 | 0.31 | 1034348 | 1034341 | 21844 | 28.74 | 0.0048 | 0.0005 | 7335 | 0.0013 | 198 | 0.71 |
| 139 | Z38481_clat_CZ_R | MIL | plate 2 | 1 | 1.21 | 4027856 | 4027839 | 35967 | 23.20 | 0.0052 | 0.0007 | 16997 | 0.0032 | 522 | 0.33 |
| 140 | Z38482_clat_CZ_R | MIL | plate 2 | 1 | 0.68 | 2279423 | 2279409 | 38299 | 37.58 | 0.0047 | 0.0007 | 19064 | 0.0030 | 548 | 0.29 |
| 141 | Z38483_clat_CZ_R | MIL | plate 2 | 1 | 0.66 | 2206354 | 2206346 | 38639 | 19.74 | 0.0052 | 0.0007 | 18049 | 0.0032 | 520 | 0.35 |
| 142 | Z38506_clat_AT | FEL | plate 3 | 1 | 1.48 | 4928122 | 4928119 | 50425 | 23.54 | 0.0075 | 0.0006 | 25986 | 0.0027 | 192 | 0.79 |
| 143 | Z38507_clat_AT | FEL | plate 3 | 1 | 0.51 | 1713807 | 1713806 | 43252 | 18.50 | 0.0077 | 0.0006 | 23956 | 0.0032 | 197 | 0.79 |
| 144 | Z38508_clat_AT | FEL | plate 3 | 1 | 0.74 | 2472952 | 2472950 | 81019 | 21.23 | 0.0056 | 0.0005 | 44043 | 0.0021 | 230 | 0.75 |
| 145 | Z38509_clat_AT | FEL | plate 3 | 1 | 1.98 | 6613324 | 6613320 | 46415 | 28.67 | 0.0074 | 0.0006 | 25030 | 0.0025 | 185 | 0.80 |
| 146 | Z38510_clat_AT | FEL | plate 3 | 1 | 1.44 | 4787062 | 4787057 | 63152 | 23.45 | 0.0070 | 0.0005 | 32035 | 0.0029 | 269 | 0.70 |
| 147 | Z38511_clat_AT | FEL | plate 3 | 1 | 0.84 | 2809781 | 2809780 | 56428 | 30.74 | 0.0066 | 0.0006 | 30901 | 0.0027 | 222 | 0.75 |
| 148 | Z38512_clat_AT | FEL | plate 3 | 1 | 1.34 | 4480352 | 4480351 | 94785 | 18.37 | 0.0056 | 0.0005 | 49826 | 0.0018 | 243 | 0.73 |
| 149 | Z38513_clat_AT | FEL | plate 3 | 1 | 0.81 | 2686872 | 2686859 | 73929 | 27.61 | 0.0034 | 0.0006 | 43059 | 0.0015 | 532 | 0.31 |

| | | | | | | | | | | | | | | | |
|---|---|---|---|---|---|---|---|---|---|---|---|---|---|---|---|
| 150 | Z38514_clat_AT | FEL | plate 3 | 1 | 0.58 | 1943643 | 1943630 | 28186 | 24.70 | 0.0049 | 0.0006 | 16722 | 0.0026 | 489 | 0.37 |
| 151 | Z38515_clat_AT | FEL | plate 3 | 1 | 0.57 | 1912284 | 1912271 | 38883 | 35.02 | 0.0048 | 0.0006 | 22869 | 0.0024 | 518 | 0.32 |
| 152 | Z38604_clat_FI_U | MAL | plate 2 | 1 | 0.72 | 2409864 | 2409853 | 60460 | 21.72 | 0.0048 | 0.0006 | 22558 | 0.0028 | 570 | 0.31 |
| 153 | Z38605_clat_FI_U | MAL | plate 2 | 1 | 0.70 | 2336972 | 2336966 | 55626 | 22.11 | 0.0049 | 0.0006 | 21632 | 0.0028 | 587 | 0.29 |
| 154 | Z38606_clat_FI_U | MAL | plate 2 | 1 | 0.53 | 1774530 | 1774528 | 53110 | 15.21 | 0.0050 | 0.0007 | 20574 | 0.0028 | 572 | 0.30 |
| 155 | Z38607_clat_FI_U | MAL | plate 2 | 1 | 0.51 | 1714208 | 1714205 | 49219 | 19.55 | 0.0048 | 0.0006 | 20445 | 0.0028 | 577 | 0.31 |
| 156 | Z38608_clat_FI_U | MAL | plate 2 | 1 | 0.71 | 2356692 | 2356682 | 65242 | 22.15 | 0.0050 | 0.0006 | 23517 | 0.0025 | 576 | 0.30 |
| 157 | Z38609_clat_FI_U | MAL | plate 2 | 1 | 0.56 | 1861547 | 1861537 | 57709 | 17.11 | 0.0046 | 0.0006 | 23302 | 0.0025 | 574 | 0.30 |
| 158 | Z38610_clat_FI_U | MAL | plate 2 | 1 | 0.54 | 1801325 | 1801318 | 57699 | 20.01 | 0.0044 | 0.0006 | 21985 | 0.0024 | 566 | 0.30 |
| 159 | Z38611_clat_FI_U | MAL | plate 2 | 1 | 0.79 | 2649283 | 2649273 | 77080 | 16.16 | 0.0046 | 0.0006 | 24057 | 0.0024 | 529 | 0.36 |
| 160 | Z38612_clat_FI_R | HAN | plate 2 | 1 | 0.48 | 1594690 | 1594684 | 51626 | 23.45 | 0.0050 | 0.0007 | 19843 | 0.0028 | 558 | 0.33 |
| 161 | Z38613_clat_FI_R | HAN | plate 2 | 1 | 0.10 | 325108 | 325106 | 27992 | 10.06 | 0.0054 | 0.0008 | 14654 | 0.0032 | 516 | 0.34 |
| 162 | Z38614_clat_FI_R | HAN | plate 2 | 1 | 0.59 | 1959546 | 1959539 | 52970 | 24.29 | 0.0045 | 0.0007 | 20819 | 0.0029 | 561 | 0.30 |
| 163 | Z38615_clat_FI_R | HAN | plate 2 | 1 | 0.87 | 2883473 | 2883457 | 63857 | 28.29 | 0.0045 | 0.0006 | 22187 | 0.0025 | 541 | 0.34 |
| 164 | Z38616_clat_FI_R | HAN | plate 2 | 1 | 0.53 | 1770583 | 1770573 | 52736 | 20.29 | 0.0048 | 0.0007 | 20709 | 0.0028 | 589 | 0.29 |
| 165 | Z38617_clat_FI_R | HAN | plate 2 | 1 | 0.47 | 1558955 | 1558947 | 46778 | 24.30 | 0.0049 | 0.0006 | 20275 | 0.0031 | 569 | 0.29 |
| 166 | Z38618_clat_FI_R | HAN | plate 2 | 1 | 0.57 | 1909664 | 1909658 | 49449 | 26.67 | 0.0049 | 0.0006 | 20480 | 0.0029 | 544 | 0.32 |
| 167 | Z38619_clat_FI_R | HAN | plate 2 | 1 | 0.58 | 1940543 | 1940531 | 68416 | 19.58 | 0.0050 | 0.0007 | 23252 | 0.0026 | 575 | 0.30 |
| 168 | Z38630_clat_FI_R | HAN | plate 2 | 1 | 0.47 | 1575852 | 1575850 | 48883 | 21.35 | 0.0044 | 0.0007 | 19449 | 0.0029 | 596 | 0.28 |
| 169 | Z38631_clat_FI_R | HAN | plate 2 | 1 | 0.55 | 1843278 | 1843275 | 45736 | 30.40 | 0.0046 | 0.0007 | 19219 | 0.0030 | 587 | 0.26 |
| 170 | Z38632_clat_FI_R | HAN | plate 2 | 1 | 0.46 | 1516677 | 1516672 | 53987 | 15.15 | 0.0048 | 0.0007 | 19776 | 0.0028 | 571 | 0.29 |
| 171 | Z38633_clat_FI_R | HAN | plate 2 | 1 | 1.04 | 3465587 | 3465570 | 69122 | 41.27 | 0.0043 | 0.0006 | 20539 | 0.0026 | 538 | 0.34 |
| 172 | Z38634_clat_FI_R | HAN | plate 2 | 1 | 0.31 | 1046465 | 1046455 | 41933 | 15.31 | 0.0048 | 0.0007 | 17530 | 0.0031 | 573 | 0.31 |
| 173 | Z38635_clat_FI_R | HAN | plate 2 | 1 | 0.51 | 1691456 | 1691452 | 53843 | 18.78 | 0.0046 | 0.0007 | 19762 | 0.0027 | 585 | 0.29 |
| 174 | Z38636_clat_FI_R | HAN | plate 2 | 1 | 0.44 | 1472220 | 1472218 | 50099 | 16.97 | 0.0047 | 0.0006 | 18998 | 0.0029 | 563 | 0.33 |
| 175 | Z38637_clat_FI_R | HAN | plate 2 | 1 | 0.81 | 2688174 | 2688166 | 83380 | 20.22 | 0.0054 | 0.0007 | 24099 | 0.0025 | 582 | 0.30 |
| 176 | Z38641_clat_FI_U | KUN | plate 3 | 1 | 0.73 | 2427925 | 2427915 | 48543 | 25.02 | 0.0040 | 0.0006 | 20927 | 0.0023 | 575 | 0.29 |
| 177 | Z38642_clat_FI_U | KUN | plate 3 | 1 | 0.89 | 2972370 | 2972360 | 68473 | 18.33 | 0.0043 | 0.0006 | 25090 | 0.0022 | 551 | 0.33 |
| 178 | Z38643_clat_FI_U | KUN | plate 3 | 1 | 0.64 | 2134815 | 2134806 | 56528 | 22.21 | 0.0047 | 0.0006 | 23759 | 0.0026 | 590 | 0.28 |
| 179 | Z38644_clat_FI_U | KUN | plate 3 | 1 | 0.47 | 1552230 | 1552219 | 50267 | 22.11 | 0.0047 | 0.0006 | 21748 | 0.0026 | 577 | 0.28 |
| 180 | Z38645_clat_FI_U | KUN | plate 3 | 1 | 0.68 | 2276747 | 2276736 | 79936 | 14.93 | 0.0052 | 0.0006 | 27862 | 0.0021 | 556 | 0.32 |
| 181 | Z38648_clat_FI_R | VIR | plate 2 | 1 | 0.93 | 3094280 | 3094267 | 56845 | 22.90 | 0.0046 | 0.0006 | 22674 | 0.0025 | 536 | 0.32 |

| | | | | | | | | | | | | | | | |
|---|---|---|---|---|---|---|---|---|---|---|---|---|---|---|---|
| 182 | Z41211_clat_Fl_U | HEI | plate 2 | 1 | 1.56 | 5188924 | 5188902 | 30197 | 36.16 | 0.0054 | 0.0009 | 14860 | 0.0032 | 527 | 0.32 |
| 183 | Z41212_clat_Fl_U | HEI | plate 2 | 1 | 0.24 | 791683 | 791679 | 33581 | 18.51 | 0.0051 | 0.0007 | 16845 | 0.0032 | 581 | 0.25 |
| 184 | Z41213_clat_Fl_U | HEI | plate 2 | 1 | 0.52 | 1721012 | 1721006 | 46562 | 30.15 | 0.0048 | 0.0006 | 18797 | 0.0031 | 595 | 0.25 |
| 185 | Z41214_clat_Fl_U | HEI | plate 2 | 1 | 0.38 | 1250814 | 1250812 | 44533 | 17.36 | 0.0048 | 0.0006 | 18639 | 0.0032 | 601 | 0.28 |
| 186 | Z41215_clat_Fl_U | HEI | plate 2 | 1 | 0.51 | 1685078 | 1685073 | 58621 | 16.07 | 0.0052 | 0.0006 | 20613 | 0.0029 | 588 | 0.28 |
| 187 | Z41216_clat_Fl_U | HEI | plate 2 | 1 | 0.55 | 1820481 | 1820479 | 56691 | 24.11 | 0.0049 | 0.0006 | 20253 | 0.0029 | 579 | 0.28 |
| 188 | Z41217_clat_Fl_U | HEI | plate 2 | 1 | 0.37 | 1247011 | 1247007 | 43050 | 12.05 | 0.0052 | 0.0007 | 17891 | 0.0030 | 587 | 0.27 |
| 189 | Z41219_clat_Fl_U | HEI | plate 2 | 1 | 0.43 | 1445393 | 1445391 | 36346 | 17.34 | 0.0049 | 0.0007 | 17745 | 0.0031 | 596 | 0.26 |
| 190 | Z41224_clat_Fl_U | HEI | plate 2 | 1 | 0.63 | 2115102 | 2115096 | 45627 | 31.52 | 0.0044 | 0.0006 | 20599 | 0.0028 | 576 | 0.28 |
| 191 | Z41225_clat_Fl_U | HEI | plate 2 | 1 | 0.51 | 1689741 | 1689738 | 42471 | 17.73 | 0.0045 | 0.0007 | 20602 | 0.0027 | 600 | 0.28 |
| 192 | Z41226_clat_Fl_U | HEI | plate 2 | 1 | 0.77 | 2578053 | 2578046 | 64958 | 17.34 | 0.0046 | 0.0007 | 23047 | 0.0026 | 560 | 0.31 |
| 193 | Z41227_clat_Fl_U | HEI | plate 2 | 1 | 0.58 | 1948940 | 1948934 | 49327 | 21.01 | 0.0045 | 0.0007 | 20659 | 0.0028 | 564 | 0.32 |
| 194 | Z41228_clat_Fl_U | HEI | plate 2 | 1 | 0.42 | 1383978 | 1383975 | 45523 | 13.21 | 0.0049 | 0.0006 | 18653 | 0.0028 | 585 | 0.28 |
| 195 | Z41229_clat_Fl_U | HEI | plate 2 | 1 | 0.43 | 1447589 | 1447585 | 43908 | 15.73 | 0.0046 | 0.0007 | 18551 | 0.0030 | 596 | 0.27 |
| 196 | Z41230_clat_Fl_U | HEI | plate 2 | 1 | 0.66 | 2205916 | 2205910 | 59092 | 24.00 | 0.0044 | 0.0007 | 20860 | 0.0026 | 535 | 0.35 |
| 197 | Z41232_clat_Fl_R | VIR | plate 2 | 1 | 0.66 | 2210607 | 2210600 | 49397 | 23.62 | 0.0043 | 0.0006 | 20324 | 0.0025 | 526 | 0.30 |
| 198 | Z41233_clat_Fl_R | VIR | plate 2 | 1 | 0.38 | 1277899 | 1277891 | 41481 | 16.37 | 0.0047 | 0.0007 | 19225 | 0.0028 | 581 | 0.29 |
| 199 | Z41234_clat_Fl_R | VIR | plate 2 | 1 | 0.55 | 1836398 | 1836386 | 49008 | 24.93 | 0.0046 | 0.0006 | 21492 | 0.0027 | 567 | 0.31 |
| 200 | Z41235_clat_Fl_R | VIR | plate 2 | 1 | 0.61 | 2032121 | 2032108 | 50723 | 19.10 | 0.0044 | 0.0007 | 21594 | 0.0025 | 580 | 0.29 |
| 201 | Z41236_clat_Fl_R | VIR | plate 2 | 1 | 0.62 | 2076962 | 2076956 | 49476 | 23.53 | 0.0046 | 0.0006 | 21289 | 0.0027 | 581 | 0.29 |
| 202 | Z41237_clat_Fl_R | VIR | plate 2 | 1 | 0.30 | 1012551 | 1012547 | 32201 | 18.32 | 0.0048 | 0.0006 | 17367 | 0.0031 | 616 | 0.27 |
| 203 | Z41238_clat_Fl_R | VIR | plate 2 | 1 | 0.56 | 1879042 | 1879037 | 49334 | 20.83 | 0.0047 | 0.0006 | 20767 | 0.0028 | 557 | 0.30 |
| 204 | Z41239_clat_Fl_R | VIR | plate 2 | 1 | 0.58 | 1925853 | 1925845 | 48164 | 20.00 | 0.0048 | 0.0007 | 20682 | 0.0027 | 571 | 0.31 |
| 205 | Z41241_clat_Fl_R | VIR | plate 2 | 1 | 0.83 | 2769256 | 2769246 | 42499 | 18.42 | 0.0045 | 0.0007 | 19361 | 0.0027 | 576 | 0.29 |
| 206 | Z41242_clat_Fl_R | VIR | plate 2 | 1 | 0.69 | 2306481 | 2306468 | 58825 | 21.93 | 0.0047 | 0.0007 | 22456 | 0.0025 | 582 | 0.28 |
| 207 | Z41243_clat_Fl_R | VIR | plate 2 | 1 | 0.30 | 983990 | 983986 | 32139 | 16.32 | 0.0048 | 0.0007 | 16379 | 0.0028 | 586 | 0.28 |
| 208 | Z41244_clat_Fl_R | VIR | plate 2 | 1 | 0.91 | 3026861 | 3026846 | 90825 | 16.92 | 0.0045 | 0.0006 | 26246 | 0.0020 | 513 | 0.35 |
| 209 | Z41245_clat_Fl_R | VIR | plate 2 | 1 | 0.55 | 1834672 | 1834666 | 45869 | 16.43 | 0.0047 | 0.0007 | 19963 | 0.0026 | 589 | 0.28 |
| 210 | Z41270_clat_AT | KLA | plate 3 | 1 | 0.71 | 2353526 | 2353514 | 28556 | 52.78 | 0.0056 | 0.0007 | 15991 | 0.0024 | 435 | 0.43 |
| 211 | Z41271_clat_AT | KLA | plate 3 | 1 | 0.48 | 1592763 | 1592757 | 43515 | 22.23 | 0.0049 | 0.0007 | 24420 | 0.0023 | 503 | 0.34 |
| 212 | Z41319_clat_AT | DRO | plate 1 | 1 | 0.56 | 1875015 | 1875007 | 25825 | 27.40 | 0.0069 | 0.0008 | 9773 | 0.0022 | 341 | 0.53 |
| 213 | Z41320_clat_AT | DRO | plate 1 | 1 | 0.44 | 1452194 | 1452181 | 22963 | 17.30 | 0.0079 | 0.0007 | 8890 | 0.0023 | 319 | 0.56 |

| | | | | | | | | | | | | | | | |
|---|---|---|---|---|---|---|---|---|---|---|---|---|---|---|---|
| 214 | Z41321_clat_AT | DRO | plate 1 | 1 | 0.42 | 1403323 | 1403318 | 19153 | 37.47 | 0.0071 | 0.0006 | 7092 | 0.0020 | 273 | 0.62 |
| 215 | Z41322_clat_AT | DRO | plate 1 | 1 | 0.28 | 928707 | 928705 | 34265 | 11.77 | 0.0056 | 0.0007 | 11705 | 0.0022 | 335 | 0.54 |
| 216 | Z41323_clat_AT | DRO | plate 1 | 1 | 0.93 | 3102361 | 3102351 | 26139 | 73.41 | 0.0057 | 0.0006 | 11678 | 0.0026 | 422 | 0.43 |
| 217 | Z41324_clat_AT | DRO | plate 1 | 1 | 0.56 | 1857129 | 1857113 | 28581 | 34.42 | 0.0058 | 0.0006 | 12476 | 0.0025 | 404 | 0.47 |
| 218 | Z41325_clat_AT | DRO | plate 1 | 1 | 0.24 | 792037 | 792033 | 13594 | 29.58 | 0.0074 | 0.0007 | 3979 | 0.0017 | 153 | 0.81 |
| 219 | Z41326_clat_AT | DRO | plate 1 | 1 | 0.35 | 1151985 | 1151977 | 24147 | 25.52 | 0.0067 | 0.0007 | 8729 | 0.0024 | 315 | 0.56 |
| 220 | Z41328_clat_AT | DRO | plate 1 | 1 | 0.73 | 2449356 | 2449303 | 80482 | 18.85 | 0.0046 | 0.0007 | 22065 | 0.0024 | 455 | 0.39 |
| 221 | Z41329_clat_AT | DRO | plate 1 | 1 | 0.86 | 2874792 | 2874772 | 63905 | 23.07 | 0.0049 | 0.0006 | 19353 | 0.0026 | 458 | 0.38 |
| 222 | Z41330_clat_AT | DRO | plate 1 | 1 | 1.23 | 4111341 | 4111323 | 54712 | 48.67 | 0.0050 | 0.0007 | 18499 | 0.0027 | 460 | 0.38 |
| 223 | Z41331_clat_AT | DRO | plate 1 | 1 | 1.11 | 3698031 | 3698009 | 68487 | 17.53 | 0.0056 | 0.0007 | 19975 | 0.0026 | 466 | 0.37 |
| 224 | Z41333_clat_AT | DRO | plate 1 | 1 | 1.29 | 4307408 | 4307388 | 83677 | 20.68 | 0.0050 | 0.0006 | 25665 | 0.0023 | 506 | 0.34 |
| 225 | Z41335_clat_AT | DRO | plate 1 | 1 | 1.06 | 3526037 | 3526020 | 74318 | 23.17 | 0.0046 | 0.0006 | 24249 | 0.0023 | 469 | 0.39 |
| 226 | Z41337_clat_AT | DRO | plate 1 | 1 | 1.11 | 3711609 | 3711596 | 39641 | 66.11 | 0.0051 | 0.0006 | 16745 | 0.0027 | 499 | 0.33 |
| 227 | Z41626_clat_AT | FEL | plate 3 | 1 | 1.07 | 3550056 | 3550033 | 207107 | 15.96 | 0.0017 | 0.0007 | 134888 | 0.0005 | 354 | 0.54 |
| 228 | Z41628_clat_AT | FEL | plate 3 | 1 | 0.25 | 831737 | 831734 | 17285 | 26.78 | 0.0052 | 0.0007 | 9154 | 0.0018 | 313 | 0.59 |
| 229 | Z41629_clat_AT | FEL | plate 3 | 1 | 0.57 | 1889507 | 1889492 | 53074 | 21.66 | 0.0029 | 0.0006 | 28934 | 0.0010 | 349 | 0.54 |
| 230 | Z67784_clat_EE_R | KAR | plate 7 | 2 | 0.48 | 1603973 | 1603721 | 26770 | 44.22 | 0.0077 | 0.0009 | 9565 | 0.0025 | 381 | 0.50 |
| 231 | Z67785_clat_EE_R | KAR | plate 7 | 2 | 0.03 | 115465 | 115452 | 14481 | 3.69 | 0.0088 | 0.0006 | 2891 | 0.0017 | 182 | 0.75 |
| 232 | Z67786_clat_EE_R | KAR | plate 7 | 2 | 0.82 | 2732228 | 2731821 | 71402 | 23.90 | 0.0028 | 0.0004 | 23898 | 0.0007 | 209 | 0.73 |
| 233 | Z67787_clat_EE_R | KAR | plate 7 | 2 | 0.17 | 574991 | 574917 | 21507 | 21.60 | 0.0080 | 0.0007 | 6679 | 0.0021 | 289 | 0.62 |
| 234 | Z67788_clat_EE_R | KAR | plate 7 | 2 | 1.16 | 3881130 | 3880471 | 37140 | 82.31 | 0.0057 | 0.0006 | 14295 | 0.0028 | 500 | 0.35 |
| 235 | Z67789_clat_EE_R | KAR | plate 7 | 2 | 1.43 | 4779111 | 4778533 | 20328 | 12.18 | 0.0041 | 0.0005 | 4798 | 0.0008 | 114 | 0.86 |
| 236 | Z67790_clat_EE_R | KAR | plate 7 | 2 | 0.01 | 47396 | 47392 | 10217 | 3.54 | 0.0095 | 0.0011 | 2098 | 0.0015 | 138 | 0.82 |
| 237 | Z67791_clat_EE_R | KAR | plate 7 | 2 | 0.05 | 161562 | 161548 | 16994 | 7.59 | 0.0087 | 0.0007 | 5739 | 0.0024 | 266 | 0.65 |
| 238 | Z67804_clat_EE_U | TAR | plate 7 | 2 | 0.66 | 2208045 | 2207771 | 50651 | 16.72 | 0.0054 | 0.0005 | 18419 | 0.0028 | 530 | 0.32 |
| 239 | Z67805_clat_EE_U | TAR | plate 7 | 2 | 0.38 | 1272877 | 1272675 | 51663 | 13.06 | 0.0053 | 0.0005 | 17571 | 0.0029 | 503 | 0.34 |
| 240 | Z67806_clat_EE_U | TAR | plate 7 | 2 | 0.52 | 1734675 | 1734441 | 62560 | 15.64 | 0.0050 | 0.0006 | 19510 | 0.0028 | 515 | 0.33 |
| 241 | Z67807_clat_EE_U | TAR | plate 7 | 2 | 0.58 | 1939567 | 1939309 | 54967 | 19.35 | 0.0052 | 0.0005 | 18478 | 0.0029 | 541 | 0.33 |
| 242 | Z67808_clat_EE_U | TAR | plate 7 | 2 | 0.55 | 1841406 | 1841164 | 66833 | 18.69 | 0.0051 | 0.0005 | 18979 | 0.0030 | 523 | 0.36 |
| 243 | Z67809_clat_EE_U | TAR | plate 7 | 2 | 0.59 | 1963415 | 1963191 | 49218 | 21.93 | 0.0064 | 0.0005 | 18087 | 0.0031 | 558 | 0.27 |
| 244 | Z67810_clat_EE_U | TAR | plate 7 | 2 | 0.92 | 3066172 | 3065722 | 34982 | 36.59 | 0.0053 | 0.0005 | 12797 | 0.0023 | 441 | 0.41 |
| 245 | Z67811_clat_EE_U | TAR | plate 7 | 2 | 0.47 | 1562311 | 1562107 | 45642 | 14.95 | 0.0055 | 0.0006 | 18010 | 0.0027 | 538 | 0.30 |

| | | | | | | | | | | | | | | | |
|---|---|---|---|---|---|---|---|---|---|---|---|---|---|---|---|
| 246 | Z67812_clat_EE_R | PAL | plate 7 | 2 | 0.88 | 2920525 | 2920097 | 69047 | 14.16 | 0.0053 | 0.0005 | 21530 | 0.0027 | 549 | 0.33 |
| 247 | Z67813_clat_EE_R | PAL | plate 7 | 2 | 0.86 | 2882072 | 2881672 | 70201 | 18.27 | 0.0053 | 0.0006 | 20807 | 0.0028 | 544 | 0.36 |
| 248 | Z67814_clat_EE_R | PAL | plate 7 | 2 | 0.88 | 2932684 | 2932277 | 72128 | 20.32 | 0.0047 | 0.0005 | 20469 | 0.0027 | 558 | 0.33 |
| 249 | Z67815_clat_EE_R | PAL | plate 7 | 2 | 0.52 | 1724573 | 1724310 | 88829 | 8.28 | 0.0059 | 0.0006 | 21539 | 0.0026 | 562 | 0.31 |
| 250 | Z67816_clat_EE_R | PAL | plate 7 | 2 | 0.34 | 1125180 | 1124996 | 51983 | 12.21 | 0.0050 | 0.0006 | 17362 | 0.0030 | 565 | 0.31 |
| 251 | Z67817_clat_EE_U | HAA | plate 7 | 2 | 0.57 | 1893117 | 1892853 | 38343 | 30.10 | 0.0058 | 0.0006 | 17616 | 0.0031 | 569 | 0.29 |
| 252 | Z67818_clat_EE_U | HAA | plate 7 | 2 | 0.58 | 1948612 | 1948283 | 60454 | 12.83 | 0.0056 | 0.0006 | 17689 | 0.0029 | 485 | 0.37 |
| 253 | Z67819_clat_EE_U | HAA | plate 7 | 2 | 0.68 | 2281736 | 2281381 | 56893 | 19.62 | 0.0051 | 0.0005 | 20141 | 0.0029 | 565 | 0.30 |
| 254 | Z67820_clat_EE_U | HAA | plate 7 | 2 | 0.97 | 3242304 | 3241859 | 50148 | 27.77 | 0.0054 | 0.0005 | 19899 | 0.0031 | 592 | 0.27 |
| 255 | Z67821_clat_EE_U | HAA | plate 7 | 2 | 0.58 | 1918521 | 1918284 | 56798 | 17.76 | 0.0053 | 0.0005 | 19716 | 0.0030 | 579 | 0.29 |
| 256 | Z67822_clat_EE_U | HAA | plate 7 | 2 | 0.94 | 3141693 | 3141288 | 57690 | 44.18 | 0.0052 | 0.0005 | 18902 | 0.0029 | 569 | 0.30 |
| 257 | Z67823_clat_FR_U | PET | plate 7 | 2 | 0.69 | 2291289 | 2290949 | 65019 | 21.24 | 0.0052 | 0.0005 | 19316 | 0.0030 | 539 | 0.31 |
| 258 | Z67824_clat_FR_U | PET | plate 7 | 2 | 0.44 | 1459656 | 1459464 | 50465 | 17.12 | 0.0053 | 0.0006 | 18484 | 0.0031 | 541 | 0.31 |
| 259 | Z67825_clat_FR_U | PET | plate 7 | 2 | 0.47 | 1568025 | 1567794 | 57525 | 15.96 | 0.0049 | 0.0006 | 18745 | 0.0029 | 517 | 0.35 |
| 260 | Z67826_clat_FR_U | PET | plate 7 | 2 | 0.56 | 1861497 | 1861248 | 60180 | 18.72 | 0.0052 | 0.0006 | 18960 | 0.0030 | 519 | 0.34 |
| 261 | Z67827_clat_FR_R | SAV | plate 7 | 2 | 0.67 | 2241482 | 2241165 | 45181 | 39.05 | 0.0057 | 0.0005 | 17499 | 0.0031 | 544 | 0.30 |
| 262 | Z67828_clat_FR_R | SAV | plate 7 | 2 | 0.80 | 2650617 | 2650229 | 39343 | 18.10 | 0.0056 | 0.0005 | 16375 | 0.0029 | 546 | 0.29 |
| 263 | Z67868_clat_EE_R | KAR | plate 6 | 2 | 0.21 | 683596 | 683504 | 47637 | 10.53 | 0.0057 | 0.0006 | 23709 | 0.0018 | 463 | 0.40 |
| 264 | Z67869_clat_EE_R | KAR | plate 6 | 2 | 0.06 | 216512 | 216474 | 12019 | 9.19 | 0.0033 | 0.0005 | 5206 | 0.0004 | 36 | 0.95 |
| 265 | Z67870_clat_EE_R | KAR | plate 6 | 2 | 0.06 | 213644 | 213617 | 25921 | 5.80 | 0.0028 | 0.0006 | 11384 | 0.0010 | 81 | 0.91 |
| 266 | Z67871_clat_EE_R | KAR | plate 6 | 2 | 0.53 | 1782648 | 1782390 | 51148 | 15.99 | 0.0061 | 0.0006 | 24957 | 0.0015 | 447 | 0.42 |
| 267 | Z67872_clat_EE_R | KAR | plate 6 | 2 | 0.17 | 572649 | 572563 | 39100 | 8.45 | 0.0042 | 0.0006 | 20152 | 0.0016 | 397 | 0.49 |
| 268 | Z67873_clat_EE_R | KAR | plate 6 | 2 | 0.19 | 617871 | 617788 | 45164 | 9.72 | 0.0061 | 0.0006 | 22325 | 0.0023 | 532 | 0.32 |
| 269 | Z67874_clat_EE_R | KAR | plate 6 | 2 | 0.14 | 473060 | 473006 | 28576 | 7.40 | 0.0044 | 0.0006 | 14011 | 0.0016 | 355 | 0.53 |
| 270 | Z67888_clat_FR_R | SAV | plate 6 | 2 | 0.30 | 1005274 | 1005126 | 47219 | 11.06 | 0.0067 | 0.0006 | 18209 | 0.0027 | 520 | 0.33 |
| 271 | Z67889_clat_FR_R | SAV | plate 6 | 2 | 0.75 | 2491306 | 2490973 | 52031 | 40.31 | 0.0060 | 0.0005 | 19024 | 0.0029 | 509 | 0.32 |
| 272 | Z67890_clat_FR_R | SAV | plate 6 | 2 | 0.73 | 2444205 | 2443765 | 41218 | 17.20 | 0.0065 | 0.0006 | 19036 | 0.0025 | 509 | 0.33 |
| 273 | Z67891_clat_FR_R | SAV | plate 6 | 2 | 0.15 | 497571 | 497503 | 35258 | 9.91 | 0.0061 | 0.0006 | 16311 | 0.0029 | 526 | 0.31 |
| 274 | Z67892_clat_FR_R | SAV | plate 6 | 2 | 0.41 | 1354529 | 1354325 | 58355 | 11.94 | 0.0057 | 0.0006 | 22992 | 0.0025 | 516 | 0.34 |
| 275 | Z67893_clat_FR_R | SAV | plate 6 | 2 | 0.51 | 1703916 | 1703662 | 65114 | 13.43 | 0.0053 | 0.0005 | 20847 | 0.0025 | 515 | 0.32 |
| 276 | Z67894_clat_FR_R | SAV | plate 6 | 2 | 0.37 | 1226672 | 1226490 | 51683 | 16.26 | 0.0055 | 0.0005 | 22225 | 0.0026 | 546 | 0.29 |
| 277 | Z67895_clat_FR_R | SAV | plate 6 | 2 | 0.31 | 1049984 | 1049833 | 46145 | 11.72 | 0.0057 | 0.0006 | 20036 | 0.0025 | 529 | 0.30 |

| | | | | | | | | | | | | | | | |
|---|---|---|---|---|---|---|---|---|---|---|---|---|---|---|---|
| 278 | Z67896_clat_FR_R | SAV | plate 6 | 2 | 0.28 | 919910 | 919776 | 39714 | 16.19 | 0.0057 | 0.0006 | 19431 | 0.0029 | 527 | 0.29 |
| 279 | Z67897_clat_FR_R | SAV | plate 6 | 2 | 0.47 | 1581020 | 1580774 | 51897 | 18.78 | 0.0056 | 0.0005 | 23027 | 0.0026 | 555 | 0.33 |
| 280 | Z67898_clat_FR_R | SAV | plate 6 | 2 | 0.61 | 2035613 | 2035313 | 67454 | 18.24 | 0.0052 | 0.0004 | 25413 | 0.0023 | 533 | 0.33 |
| 281 | Z67899_clat_FR_R | SAV | plate 6 | 2 | 0.50 | 1651817 | 1651597 | 51825 | 16.37 | 0.0062 | 0.0006 | 21198 | 0.0025 | 493 | 0.33 |
| 282 | Z67900_clat_FR_R | SAV | plate 6 | 2 | 0.52 | 1725792 | 1725568 | 53234 | 21.55 | 0.0053 | 0.0005 | 22875 | 0.0025 | 522 | 0.33 |
| 283 | Z67901_clat_FR_R | SAV | plate 6 | 2 | 0.51 | 1693619 | 1693369 | 54923 | 19.55 | 0.0054 | 0.0005 | 22775 | 0.0026 | 536 | 0.32 |
| 284 | Z67902_clat_FR_U | ORC | plate 6 | 2 | 0.37 | 1248103 | 1247923 | 56577 | 16.37 | 0.0054 | 0.0005 | 20501 | 0.0028 | 508 | 0.35 |
| 285 | Z67903_clat_FR_U | ORC | plate 6 | 2 | 0.50 | 1662010 | 1661751 | 61678 | 17.48 | 0.0054 | 0.0005 | 21106 | 0.0027 | 541 | 0.33 |
| 286 | Z67904_clat_FR_U | ORC | plate 6 | 2 | 0.57 | 1902269 | 1901982 | 68508 | 16.13 | 0.0058 | 0.0005 | 21452 | 0.0027 | 491 | 0.35 |
| 287 | Z67905_clat_FR_U | PET | plate 6 | 2 | 0.38 | 1250712 | 1250536 | 57639 | 15.90 | 0.0054 | 0.0005 | 20126 | 0.0028 | 501 | 0.34 |
| 288 | Z67906_clat_FR_U | PET | plate 6 | 2 | 0.49 | 1636687 | 1636437 | 56501 | 20.40 | 0.0054 | 0.0005 | 20726 | 0.0028 | 530 | 0.31 |
| 289 | Z67907_clat_FR_U | PET | plate 6 | 2 | 0.56 | 1863480 | 1863215 | 65489 | 20.51 | 0.0056 | 0.0005 | 21931 | 0.0028 | 532 | 0.33 |
| 290 | Z67908_clat_FR_U | PET | plate 6 | 2 | 0.54 | 1793152 | 1792885 | 58099 | 21.98 | 0.0053 | 0.0005 | 20980 | 0.0029 | 501 | 0.34 |
| 291 | Z67909_clat_FR_U | PET | plate 6 | 2 | 0.48 | 1599771 | 1599530 | 63571 | 18.32 | 0.0054 | 0.0005 | 21862 | 0.0027 | 546 | 0.30 |
| 292 | Z67910_clat_FR_U | PET | plate 6 | 2 | 0.40 | 1332294 | 1332105 | 55769 | 16.13 | 0.0052 | 0.0004 | 19883 | 0.0028 | 534 | 0.31 |
| 293 | Z67911_clat_FR_U | PET | plate 6 | 2 | 0.55 | 1820184 | 1819929 | 66518 | 18.44 | 0.0051 | 0.0004 | 21563 | 0.0027 | 514 | 0.35 |
| 294 | Z67912_clat_FR_U | PET | plate 6 | 2 | 0.51 | 1689825 | 1689572 | 65304 | 18.61 | 0.0052 | 0.0004 | 21409 | 0.0026 | 528 | 0.32 |
| 295 | Z67913_clat_FR_U | PET | plate 6 | 2 | 0.60 | 1990655 | 1990381 | 63851 | 19.74 | 0.0055 | 0.0004 | 21455 | 0.0027 | 518 | 0.37 |
| 296 | Z67914_clat_FR_U | PET | plate 6 | 2 | 0.59 | 1974469 | 1974157 | 52792 | 27.31 | 0.0054 | 0.0005 | 20687 | 0.0029 | 545 | 0.32 |
| 297 | Z67915_clat_FR_U | PET | plate 6 | 2 | 0.49 | 1641525 | 1641263 | 68987 | 16.57 | 0.0054 | 0.0005 | 21963 | 0.0026 | 526 | 0.35 |
| 298 | Z67916_clat_FR_U | PET | plate 6 | 2 | 0.61 | 2048619 | 2048336 | 66611 | 21.57 | 0.0052 | 0.0004 | 22097 | 0.0026 | 513 | 0.35 |
| 299 | Z67951_clat_EE_R | PAL | plate 6 | 2 | 0.64 | 2122489 | 2122170 | 77208 | 15.85 | 0.0056 | 0.0004 | 23305 | 0.0026 | 531 | 0.33 |
| 300 | Z67952_clat_EE_R | PAL | plate 6 | 2 | 0.41 | 1381053 | 1380824 | 70909 | 12.01 | 0.0064 | 0.0005 | 24272 | 0.0025 | 546 | 0.29 |
| 301 | Z67953_clat_EE_R | PAL | plate 6 | 2 | 0.54 | 1794381 | 1794137 | 83660 | 14.74 | 0.0055 | 0.0005 | 29002 | 0.0022 | 541 | 0.31 |
| 302 | Z67954_clat_EE_R | PAL | plate 6 | 2 | 0.49 | 1641544 | 1641296 | 64578 | 16.97 | 0.0055 | 0.0005 | 21000 | 0.0026 | 504 | 0.34 |
| 303 | Z67955_clat_EE_R | PAL | plate 6 | 2 | 0.61 | 2034856 | 2034546 | 85582 | 16.32 | 0.0052 | 0.0004 | 23705 | 0.0025 | 509 | 0.34 |
| 304 | Z67956_clat_EE_R | PAL | plate 6 | 2 | 0.69 | 2315614 | 2315324 | 77725 | 19.33 | 0.0049 | 0.0004 | 23215 | 0.0025 | 498 | 0.36 |
| 305 | Z67957_clat_EE_R | PAL | plate 6 | 2 | 0.40 | 1347537 | 1347348 | 60460 | 15.58 | 0.0054 | 0.0005 | 20788 | 0.0027 | 546 | 0.30 |
| 306 | Z67958_clat_EE_R | PAL | plate 6 | 2 | 0.54 | 1812354 | 1812043 | 65506 | 16.71 | 0.0055 | 0.0005 | 24559 | 0.0025 | 517 | 0.33 |
| 307 | Z67959_clat_EE_R | PAL | plate 6 | 2 | 0.72 | 2385160 | 2384813 | 81922 | 17.92 | 0.0055 | 0.0005 | 23941 | 0.0025 | 526 | 0.34 |
| 308 | Z67960_clat_EE_R | PAL | plate 6 | 2 | 0.51 | 1715012 | 1714736 | 67173 | 18.39 | 0.0052 | 0.0005 | 22018 | 0.0026 | 547 | 0.29 |
| 309 | Z67961_clat_EE_U | HAA | plate 6 | 2 | 0.56 | 1851488 | 1851237 | 79991 | 12.93 | 0.0048 | 0.0005 | 20344 | 0.0025 | 451 | 0.41 |

| | | | | | | | | | | | | | | | |
|---|---|---|---|---|---|---|---|---|---|---|---|---|---|---|---|
| 310 | Z67962_clat_EE_U | HAA | plate 6 | 2 | 0.49 | 1620104 | 1619844 | 68777 | 14.23 | 0.0052 | 0.0006 | 20241 | 0.0027 | 528 | 0.31 |
| 311 | Z67963_clat_EE_U | HAA | plate 6 | 2 | 0.93 | 3096921 | 3096494 | 85087 | 21.73 | 0.0051 | 0.0004 | 35400 | 0.0018 | 515 | 0.34 |
| 312 | Z67964_clat_EE_U | HAA | plate 6 | 2 | 0.88 | 2927703 | 2927277 | 68601 | 24.57 | 0.0051 | 0.0005 | 31136 | 0.0020 | 512 | 0.35 |
| 313 | Z67965_clat_EE_U | HAA | plate 6 | 2 | 1.00 | 3344484 | 3344024 | 72717 | 23.49 | 0.0046 | 0.0005 | 32689 | 0.0019 | 521 | 0.35 |
| 314 | Z67966_clat_EE_U | HAA | plate 6 | 2 | 0.33 | 1095091 | 1094957 | 49566 | 13.45 | 0.0053 | 0.0005 | 18022 | 0.0029 | 545 | 0.29 |
| 315 | Z67967_clat_EE_U | HAA | plate 6 | 2 | 0.37 | 1223700 | 1223543 | 55850 | 16.29 | 0.0052 | 0.0005 | 18872 | 0.0029 | 557 | 0.29 |
| 316 | Z67968_clat_EE_U | HAA | plate 6 | 2 | 0.42 | 1404279 | 1404076 | 64324 | 15.06 | 0.0048 | 0.0005 | 19726 | 0.0025 | 512 | 0.35 |
| 317 | Z67969_clat_EE_U | HAA | plate 6 | 2 | 0.42 | 1398858 | 1398654 | 63594 | 15.28 | 0.0059 | 0.0006 | 20280 | 0.0026 | 510 | 0.36 |
| 318 | Z67970_clat_EE_U | HAA | plate 6 | 2 | 0.51 | 1701291 | 1701055 | 66875 | 17.02 | 0.0052 | 0.0005 | 20832 | 0.0028 | 533 | 0.32 |
| 319 | Z67971_clat_EE_U | TAR | plate 6 | 2 | 0.50 | 1678182 | 1677935 | 65259 | 18.62 | 0.0051 | 0.0005 | 24242 | 0.0024 | 486 | 0.36 |
| 320 | Z67972_clat_EE_U | TAR | plate 6 | 2 | 0.84 | 2798779 | 2798358 | 102881 | 10.79 | 0.0054 | 0.0005 | 32300 | 0.0018 | 435 | 0.42 |
| 321 | Z67973_clat_EE_U | TAR | plate 6 | 2 | 0.55 | 1834951 | 1834680 | 60544 | 18.53 | 0.0052 | 0.0005 | 24440 | 0.0024 | 525 | 0.34 |
| 322 | Z67974_clat_EE_U | TAR | plate 6 | 2 | 0.45 | 1494703 | 1494484 | 45284 | 17.11 | 0.0056 | 0.0005 | 19405 | 0.0028 | 544 | 0.29 |
| 323 | Z67975_clat_EE_U | TAR | plate 6 | 2 | 0.33 | 1089151 | 1088982 | 32715 | 17.60 | 0.0053 | 0.0006 | 14579 | 0.0022 | 476 | 0.40 |
| 324 | Z67976_clat_EE_U | TAR | plate 6 | 2 | 0.39 | 1299413 | 1299217 | 56577 | 16.37 | 0.0053 | 0.0005 | 23230 | 0.0025 | 551 | 0.29 |
| 325 | Z67978_clat_EE_U | TAR | plate 6 | 2 | 0.78 | 2592202 | 2591833 | 79484 | 21.72 | 0.0051 | 0.0005 | 29668 | 0.0022 | 527 | 0.34 |

**Table S6.** Estimated effective population sizes ($N_e$) of the studied *Co. pamphilus* populations. The estimates are derived by using the method based on linkage disequilibrium and setting the minor allele frequency threshold to 0.05.

| Population[a] | Country | Environment[b] | Sample size (N) | $N_e$ | 95% CI of $N_e$[c] |
|---|---|---|---|---|---|
| BUD | BE | urban | 16 | 62.8 | 30.5. 40.5 |
| JOS | BE | urban | 16 | 9.5 | 7.9. 11.6 |
| NIS | BE | rural | 16 | ∞ | 607.9. ∞ |
| DIV | CZ | urban | 13 | ∞ | ∞ |
| MIL | CZ | rural | 16 | ∞ | ∞ |
| POH | CZ | urban | 16 | 14.1 | 11.3. 18.1 |
| HEI | FI | urban | 16 | ∞ | ∞ |
| LOF | FI | rural | 16 | 30.7 | 21.3. 51.6 |
| PIK | FI | urban | 16 | 330.2 | 77.6. ∞ |
| TUR | FI | urban | 4 | ∞ | ∞ |
| VIR | FI | rural | 16 | 45.6 | 27.7. 111.1 |
| DEC | FR | urban | 16 | ∞ | ∞ |
| PAR | FR | urban | 16 | 135.1 | 56.9. ∞ |
| PET | FR | urban | 6 | ∞ | ∞ |
| SAV | FR | rural | 18 | ∞ | ∞ |
| GAL | GR | rural | 16 | 63.2 | 32.0. 609.2 |
| KAT | GR | rural | 11 | ∞ | ∞ |
| ARR | IT | urban | 17 | ∞ | 527.7. ∞ |
| MAN | IT | rural | 18 | ∞ | ∞ |
| BRO | SE | urban | 16 | ∞ | ∞ |
| GAR | SE | urban | 15 | ∞ | ∞ |
| SON | SE | rural | 16 | ∞ | 5033.5. ∞ |

[a] See Table S1 for a key of the population abbreviations.

[b] Populations are classified as urban or rural based on the threshold of 0.2 in the proportion of urban land cover within a 2500 m radius (urban populations have urban land cover ≥0.2). See accurate values of urban land cover in Table S1.

[c] When the estimated $N_e$ is infinite, the 95% CI of it is also given as infinite because a meaningful CI cannot be derived then.

**Table S7.** Estimated effective population sizes ($N_e$) of the studied *Ch. clathrata* populations. The estimates are derived by using the method based on linkage disequilibrium and setting the minor allele frequency threshold to 0.05.

| Population[a] | Country | Environment[b] | Sample size (N) | $N_e$ | 95% CI of $N_e$[c] |
|---|---|---|---|---|---|
| DRO | AT | rural | 15 | ∞ | 2729.5, ∞ |
| FEL | AT | rural | 13 | ∞ | ∞ |
| KLA | AT | rural | 2 | ∞ | ∞ |
| BUD | BE | urban | 8 | ∞ | 34.7, ∞ |
| JOS | BE | urban | 16 | ∞ | 215.9, ∞ |
| MOE | BE | urban | 6 | ∞ | 109.3, ∞ |
| NIS | BE | rural | 15 | ∞ | 83.3, ∞ |
| BIS | CZ | urban | 11 | ∞ | ∞ |
| CHL | CZ | urban | 16 | ∞ | ∞ |
| DIV | CZ | urban | 16 | ∞ | ∞ |
| MIL | CZ | rural | 16 | 1535.4 | 39.0, ∞ |
| POH | CZ | urban | 5 | ∞ | ∞ |
| HAA | EE | urban | 16 | ∞ | 44.3, ∞ |
| KAR | EE | rural | 15 | ∞ | ∞ |
| PAL | EE | rural | 15 | ∞ | 791.0, ∞ |
| TAR | EE | urban | 15 | ∞ | 96.8, ∞ |
| HAN | FI | rural | 16 | ∞ | ∞ |
| HEI | FI | urban | 16 | ∞ | 38.7, ∞ |
| KUN | FI | rural | 5 | ∞ | ∞ |
| MAL | FI | urban | 16 | 26.3 | 13.1, 132.4 |
| VIR | FI | rural | 14 | ∞ | 37.7, ∞ |
| ORC | FR | urban | 3 | ∞ | ∞ |
| PET | FR | urban | 16 | ∞ | 71.1, ∞ |
| SAV | FR | rural | 16 | ∞ | ∞ |
| BRO | SE | urban | 16 | 49.4 | 22.4, ∞ |
| KRI | SE | rural | 2 | ∞ | ∞ |
| SON | SE | rural | 2 | ∞ | ∞ |
| TIS | SE | rural | 3 | ∞ | ∞ |

[a] See Table S1 for a key of the population abbreviations.
[b] Populations are classified as urban or rural based on the threshold of 0.2 in the proportion of urban land cover within a 2500 m radius (urban populations have urban land cover ≥0.2). See accurate values of urban land cover in Table S1.
[c] When the estimated $N_e$ is infinite, the 95% CI of it is also given as infinite because a meaningful CI cannot be derived then.
